# Dissection of genotype-phenotype relationships in *Candida parapsilosis* uncovers drivers of clinically-relevant traits

**DOI:** 10.64898/2026.05.18.725882

**Authors:** Miquel Àngel Schikora-Tamarit, Elena López-Peralta, Alejandra Roldán, Cristina de Armentia, Alba Torres-Cano, Laura Alcázar-Fuoli, CAPAR Study Group, Óscar Zaragoza, Toni Gabaldón

## Abstract

Hospital outbreaks caused by the fungal pathogen *Candida parapsilosis* are of growing concern due to their increased drug resistance and high mortality rates. However, the genetic bases of clinically-relevant traits in this species remain poorly explored. Here, we mapped genotype–phenotype relationships across 189 isolates from a multi-hospital *Candida parapsilosis* outbreak, for which we measured 61 diverse clinical phenotypes and generated complete genome sequences. As variation in previously-known genes explained little of the observed phenotypic diversity, we leveraged convergence genome-wide association studies and interpretable machine-learning models that predict phenotypes from genetic variants. These approaches identified candidate drivers of virulence and antifungal resistance, confirming expected mechanisms while uncovering novel ones. Predictive models were accurate for key traits, including azole resistance and clinical features of infected patients. Our results shed light on the genetic bases of clinically-relevant traits in a major fungal pathogen, and pave the way towards sequence-based diagnostics for improved patient outcomes.

## INTRODUCTION

Fungal infections pose a serious global health threat, causing ∼4 million deaths each year and requiring better therapeutic and diagnostic tools^1–4^. Among other pathogens, *Candida* species are a major cause of hospital-acquired infections, having high mortality in immunocompromised patients^3,5^. Given the high dynamism of *Candida* genomes, a promising strategy to improve current therapies and diagnostics is to elucidate genomic changes underlying clinically-relevant traits such as virulence and antifungal drug resistance^6–11^. Recent efforts established genotype-phenotype relationships using directed evolution^12–14^ and genome-wide comparisons of clinical isolates^9,11,15,16^, but various relevant gaps remain. For instance, most studies assessed genetic variants independently, not accounting for polygenic effects or epistatic interactions^9,16,17^. Furthermore, the focus has been almost exclusively on drug resistance, leaving aside virulence traits. Finally, most studies centered on certain “mainstream” species (e.g. *Candida albicans*, *Candida glabrata* or *Candida auris*).

Here, we investigated genotype-phenotype relationships in *Candida parapsilosis*, an emerging pathogen frequently causing nosocomial outbreaks^3,18^. This opportunistic yeast can cause invasive infections in immunocompromised individuals such as neonates, bone marrow transplant recipients and patients receiving immunosuppressive therapies^3,19^. Moreover, infections are also associated with the use of catheters, such as central venous catheters and parenteral nutrition, and have high mortality rates ranging from 20% to 45% despite active treatment^3^. The incidence of candidemia caused by *C. parapsilosis* is particularly high, being the second to third cause of candidemia depending on the country^3,5,20^. While resistance towards echinocandins and polyenes remains very rare for this species, azole resistance is steadily increasing, reaching ∼10% in some regions^21–23,3,24–26^. Furthermore, compared to other *Candida* species, *C. parapsilosis* presents an intrinsically reduced susceptibility to echinocandins^27,28^, and has a high propensity to form biofilms on prostheses and medical implants, leading to increased drug tolerance^29,30^. These factors combined make *C. parapsilosis* a significant public health threat^3,22,31^.

To address these challenges, recent studies have investigated the evolutionary mechanisms generating clinically-relevant traits, such as virulence and antifungal resistance in *C. parapsilosis* (reviewed in ^23^ and ^32^). A common approach has been to investigate whether observed resistance can be attributed to non-synonymous mutations in a few genes implicated in drug resistance in other species. For instance, studies analyzing *C. parapsilosis* serial isolates from the same patient^33,34^ or from the same outbreak^35–39^ have consistently shown that protein altering mutations in the drug targets *ERG11* (encoding lanosterol 14α-demethylase) and *FKS1* (encoding 1,3-ß-D-glucan synthase) often explain azole and echinocandin resistance, respectively. These mechanisms have been further validated through CRISPR-based mutation reintroduction assays^33,40^.

While such hypothesis-driven approaches provide valuable insights, they can only confirm known mechanisms in additional species. To circumvent this and gain a more comprehensive understanding other studies have adopted more exploratory approaches. For instance, directed evolution experiments implicated 38 genes (including *FKS1*) associated with echinocandin resistance^41^. Similarly, systematic analyses of copy-number variants (CNV) in clinical isolates suggested a putative role of *ERG11*, *CDR1B* and *RTA3* duplications in azole and miltefosine resistance^42,43^. Other studies have identified correlations between biofilm formation and point mutations in certain genes (*ALS6*, *ALS7*, *SAPP3*, *SAP7* and *SAP9*^44^; or *EFG1*^45^). More recently, machine learning has been employed to pinpoint variants associated with micafungin tolerance and heteroresistance^46^. Also, Genome-Wide Association Studies (GWAS) have identified Single Nucleotide Polymorphisms (SNPs) linked to drug resistance, albeit with mixed results. For instance, one such recent GWAS on fluconazole resistance yielded no significant hits, a result attributed to limited statistical power stemming from a small sample size (n=42) and the clonal nature of *C. parapsilosis*^43^. Conversely, a recent study reports GWAS-based associations between *ALS3* (an adhesion protein) variants and resistance to fluconazole and 5-flucytosine^45^.

Despite these advances, several important gaps remain. First, many studies suffer from small sample sizes and often lack rigorous statistical testing for genotype-phenotype associations. Second, despite their established importance^42,43^, structural variants (SVs) like inversions or translocations are often overlooked. Third, previous research has almost exclusively focused on drug resistance, although genotype-phenotype analyses could potentially unveil underlying mechanisms for virulence traits or predisposition to infecting specific patient populations / body compartments^44,45^. Fourth, current GWAS approaches may be underpowered due to their reliance on allele-counting methods that treat each SNP independently^43,45^. This approach is unsuitable for clonal species like *C. parapsilosis* and for phenotypes exhibiting allelic heterogeneity, as expected for the traits under investigation^9,47^. Instead, GWAS approaches based on genotype-phenotype convergence and variant collapsing, as recently applied in other *Candida* species^9^, may be more appropriate. Fifth, most studies used univariate approaches that examine each variant independently, which do not account for genetic interactions or the expected polygenic nature of these traits^23,32^. Therefore, multivariate approaches, such as those based on machine learning, that consider all genes simultaneously could provide novel insights and enable sequence-based trait diagnosis. While recent efforts have utilized such tools to develop predictors for drug resistance and tolerance^46,48^, there is a clear need for a more comprehensive, multi-phenotype analysis in *C. parapsilosis*.

To address these critical gaps and gain novel insights on the genetic bases of relevant traits in *C. parapsilosis*, we sequenced the genomes and phenotyped hundreds of clinical isolates collected during the course of a multi-hospital outbreak in Spain^24,25,49,50^. This extensive dataset offers unprecedented statistical power to unravel the emergence of drug resistance, virulence, and other clinically important phenotypes in a *Candida* pathogen. We first assessed the contribution of previously known genes, establishing a baseline for subsequent analyses. Given that these genes alone were insufficient to explain most traits, we employed convergence GWAS and interpretable Machine Learning (ML) approaches. Collectively, these analyses offer an integrated view of the genetic determinants underlying numerous clinically-relevant traits in *C. parapsilosis,* serving as a blueprint for studying genotype-phenotype relationships in clonally-reproducing species

## RESULTS AND DISCUSSION

### Establishing a comprehensive genomic and phenotypic resource for *C. parapsilosis* clinical isolates

We generated whole-genome sequencing data for 233 representative isolates from a 2019-2023 multi-hospital outbreak in Spain^51^. To provide a broader context, we augmented this dataset with all available *C. parapsilosis* clinical isolates from the NCBI Sequence Read Archive (SRA)^52^ (214 as of Nov 2023), totalling 447 isolates. After rigorous quality control, our final dataset comprised 365 high-quality, high-coverage genomes (189 newly sequenced, 176 from SRA), from 13 countries across North America, Asia, and Europe (**Fig. 1a, Supplementary Table 1**). To our knowledge, this collection stands as one of the largest global genomic datasets for *C. parapsilosis*^35–38,42,43,45^, and the 189 outbreak isolates represent the largest single-outbreak genomic collection reported for this species^36–38^, making it highly valuable for genotype-phenotype analysis.

**Fig. 1.**
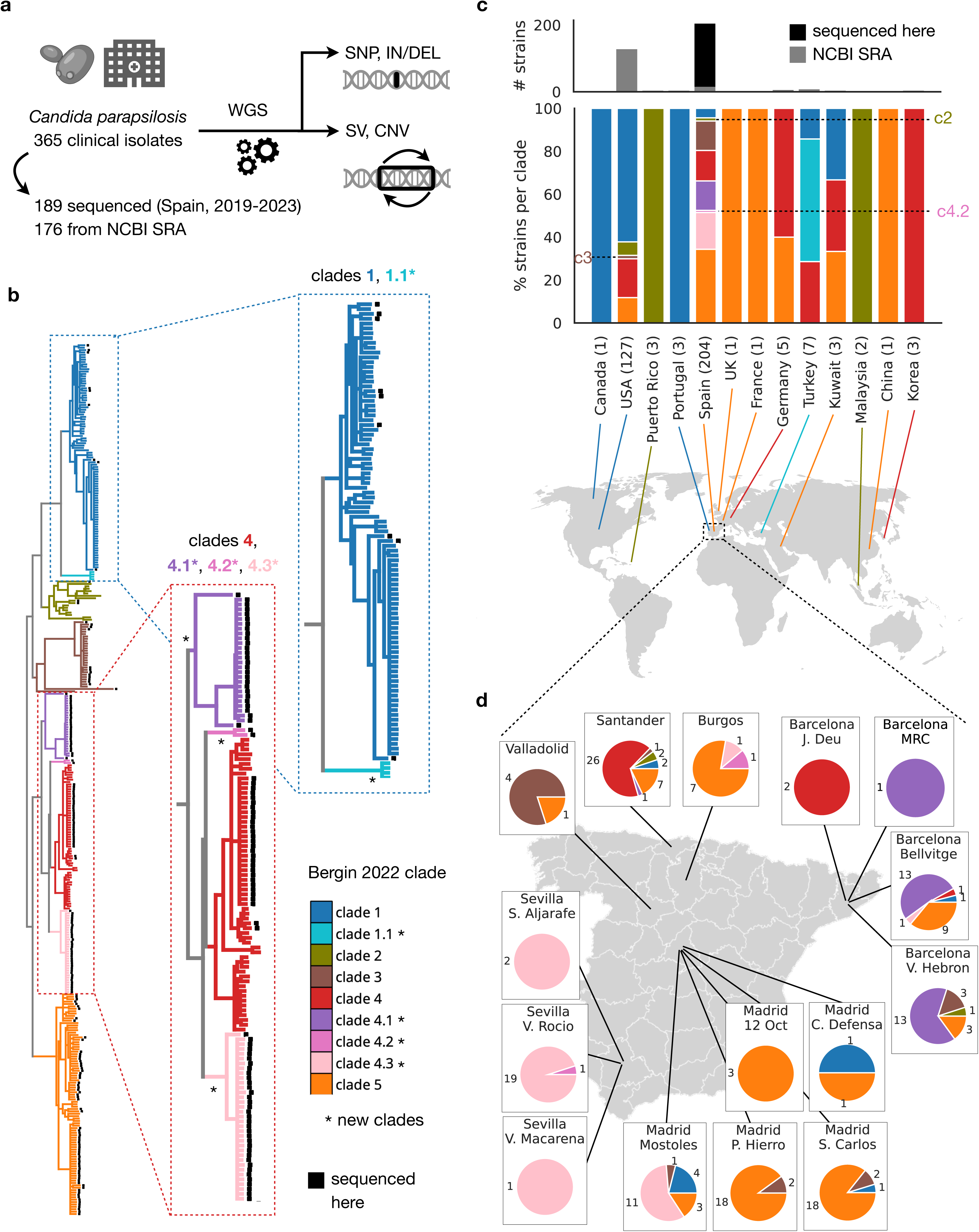
Overview of the genomic dataset produced. (a) Variant calling pipeline. For 189 isolates sequenced here and 176 available at NCBI SRA at the start of this study, we processed genomic reads to identify Single Nucleotide Polymorphisms (SNPs), small Insertions and Deletions (IN/DELs), coverage-derived Copy Number Variants (CNVs) and breakpoint-inferred Structural Variants (SVs). (b) SNP-based tree, based on bwa mem-mappings, of all the isolates analyzed here, where the black squares indicate those sequenced here. The colors correspond to different clades as defined in Bergin et. al.^42^, and the asterisks indicate new clades as compared to that study. Branches with a support < 90 are collapsed. (c) Country of origin of the strains analyzed, where the colors represent the clade as shown in (b). Note that there are 4 isolates from the SRA for which we could not get country information. (d) Geographic precedence of the 189 Spanish isolates sequenced here. The numbers indicate the strain count of each clade in each hospital. Also, the hospital names are just a reduced version, see **Supplementary Table 1** for the full hospital name.

We used PerSVade^53^ to call single-nucleotide polymorphisms (SNPs), small insertions and deletions (INDELs), structural variants (SVs) and copy-number variants (CNVs) (see **Methods, Fig. 1, Extended Data Fig. 1**). We reconstructed a SNP-based phylogenetic tree (see **Methods, Fig. 1b, Extended Data Fig. 2, Supplementary Results**), which likely accurately represents the species’ genetic structure given minimal signs of recombination (**Extended Data Fig. 3**). The resulting phylogeny provided an evolutionary framework for subsequent analysis and revealed distinct clades, largely consistent with the five previously defined (1 to 5 in ^42^). Additionally, we identified four new sub-clades (1.1, 4.1, 4.2, 4.3) (**Fig. 1b**). Sub-clade 1.1 includes isolates from a prior study^33^, while 4.1, 4.2, and 4.3 consist solely of our newly sequenced strains. Thus, while our data expand established clades, the overall genetic structure described by ^42^ remains robust. This underscores the representativeness of our dataset, supporting the generalizability of downstream genotype-phenotype analyses. These clade and sub-clade definitions and variants are a valuable resource for future epidemiological and population genomic studies. For instance, phylogeographic analysis of newly-sequenced isolates suggests both global and local propagation among Spanish clinical isolates (**Fig. 1c,d**, **Supplementary Results**).

To study genotype-phenotype relationships we retrieved various traits for the 189 newly-sequenced isolates (**Fig. 2**, **Extended Data Fig. 4,5, Supplementary Results**). For instance, we identified resistance in liquid media towards ten antifungals, based on previous susceptibility measurements on our strains^51^. Similarly, we retrieved measurements on our isolates regarding tolerance towards six disinfectants, agar invasion, pseudohyphal growth, biofilm formation, and behavior in a microfluidics catheter-like device^54^. Additionally, we performed fluconazole E-tests to infer heteroresistance, higher resistance in solid vs liquid media, and tolerance at 48h. Finally, we considered as relevant “traits” of an isolate the clinical information related to the isolation source or the age of the infected. While these “traits” may not solely reflect strain genetics, defining them as binary phenotypes would enable us to find variants predisposing strains to infect specific patient populations or body sites. Altogether, this comprehensive phenotypic collection offers an unprecedented capacity to study genotype-phenotype relationships, and our inclusion of virulence and clinical phenotypes makes it particularly unique.

**Fig. 2.**
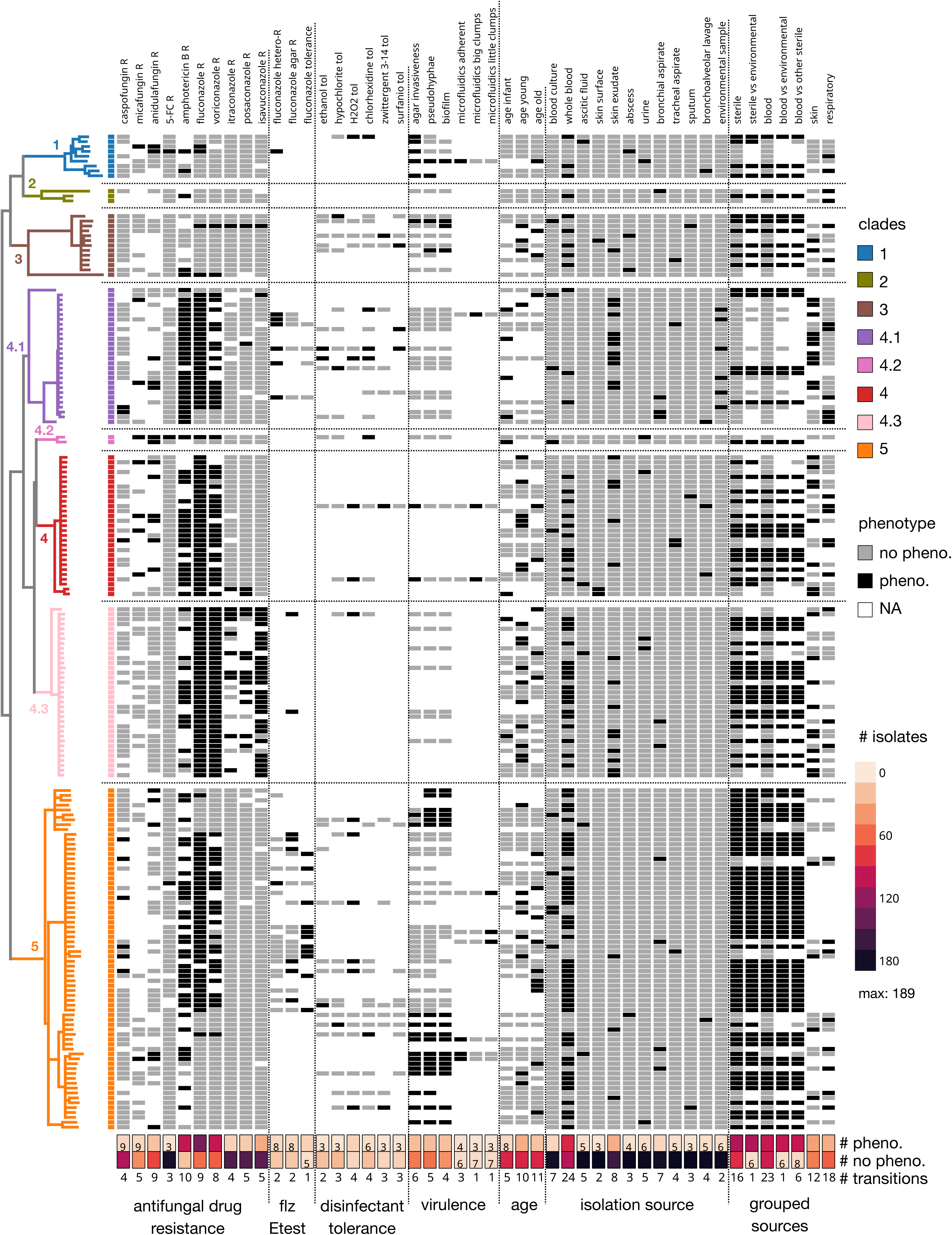
Overview of the measured phenotypes. Heatmap showing the binarized phenotypes for all strains sequenced here, including i) resistance towards ten antifungals, ii) fluconazole heteroresistance, tolerance and resistance in agar (based on E-tests performed here), iii) tolerance towards six disinfectants, iv) virulence properties (invasiveness, pseudohyphae, biofilm formation, and behavior in a microfluidics behavior), v) patient age groups, vi) isolation source, and vii) grouped isolation sources. The antifungal susceptibility data was generated in ^51^, while the disinfectant tolerance and virulence phenotypes were measured in ^54^. Note that we reduced the branch length of the long-branched outgroup of clade 3 (see Fig. 1b) to accommodate it to the plot size. Also, the number of transitions (# transitions) refers to the number of nodes of the bwa mem SNP tree (Fig. 1b) which, according to ancestral state reconstruction (ASR), have a phenotype to no phenotype change or *vice versa*. This number is calculated on the representative sample set (see **Extended Data Fig. 6a**), and in a way that all nodes are considered for the calculation of transitions (see **Supplementary Methods**). This explains why the estimates of the number of transitions shown here are different to those considered for ML and GWAS analyses (see Fig. 4d, **Methods**). For the isolation source, we only show sources that have >2 strains with and >2 strains without the source. Also, note that the NA (not applicable) category in which the phenotype was either i) not measured, ii) an intermediate phenotype (e.g. intermediate susceptibility) or iii) not related to the comparison. See **Extended Data Fig. 4,5** and **Methods** to better understand how we defined all these phenotypes. Also, note the following caveats for the virulence factors. For agar invasiveness we observed low formation in 3 isolates, which we set to “not applicable” (NA) in our analysis. Similarly, we set to NA 13 isolates that had a small capacity to form biofilms.

### Assessing genes previously related to drug resistance

Similarly to previous hypothesis-driven research, we assessed whether previously known resistance drivers in *C. parapsilosis* could explain the observed trait variability. We initially focused on established mechanisms of azole resistance, consisting of non-synonymous changes in ergosterol pathway genes (*ERG11* and *ERG3*) or regulators of efflux pumps (*TAC1*, *MRR1*, *UPC2* and *NDT80*)^23,32,40,49,55–66^. To assess their role, we inspected presence/absence patterns of gene variants across resistant and susceptible isolates (**Fig. 3, Extended Data Fig. 6a**).

**Fig. 3.**
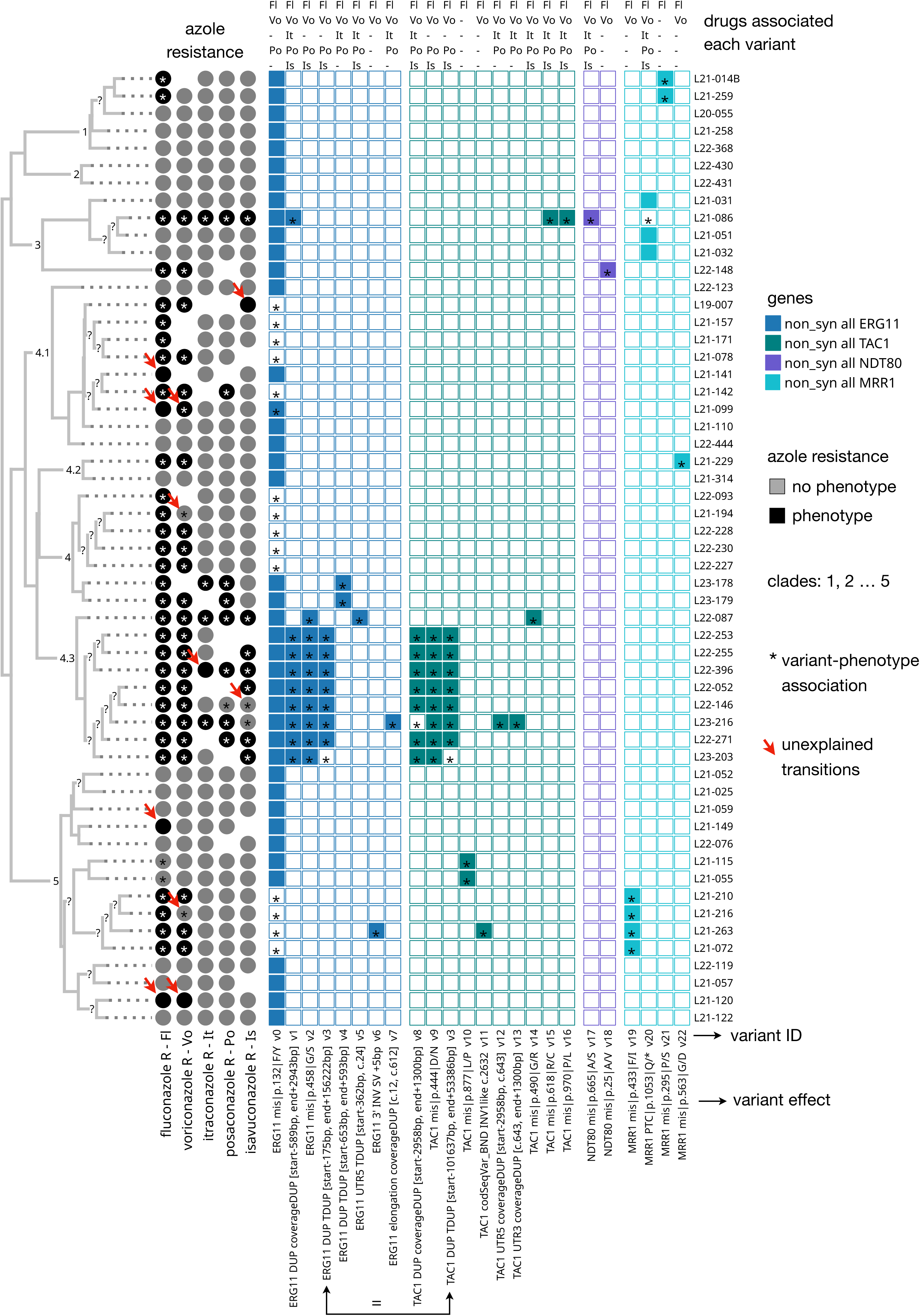
Evaluation of the known mechanisms of azole resistance. Presence absence pattern of all nonsynonymous variants in *ERG11*, *TAC1*, *MRR1* and *NDT80* which appear together with any of the azole resistance phenotypes, or in nodes that have azole resistance, as indicated by the asterisks. Only representative samples (see **Extended Data Fig. 6**) are shown for clarity (and we also removed some redundant samples manually). Note that we could not find any such variants for *UPC2* and *ERG3*, which have also been previously linked to azole resistance. The circles represent the resistance phenotypes as shown in Fig. 2. The red arrows were manually placed on samples for which a specific change in the phenotype (vs sister clades) is not related to any change in the four genes. The numbers in the tree reflect the clades as shown in Fig. 1, while the “?” indicates nodes with support <70. In addition, the table on top of the presence / absence matrix reflects the azoles to which each variant is associated at least once.

We confirmed the importance of *ERG11, TAC1, MRR1*, and *NDT80* in azole resistance, observing correlations between their variants and resistance across all tested azoles (**Fig. 3**). These include missense mutations, but also premature stop codons, segmental duplications and inversions, underscoring the importance of considering multiple variant types (**Fig. 3**). While most variants correlated with fluconazole and voriconazole resistance, only a subset influenced other azoles. Notably, we identified three pan-azole resistant isolate clusters (one in clade 3, two in 4.3), each with distinct variant combinations. For instance, a clade 3 isolate had an *ERG11* duplication, *TAC1* missense mutations (R618C, P970L), an *NDT80* missense (A665S) and a *MRR1* premature stop codon loss (**Fig. 3**). Conversely, both clade 4.3 clusters shared an *ERG11* G458S mutation and a *TAC1* missense (G490R or D444N), with additional cluster-specific duplications affecting either only *ERG11,* or both *ERG11* and *TAC1* (also described in ^51^) (**Fig. 3**). These observations suggest independent, possibly synergistic, mechanisms for azole and pan-azole resistance.

Remarkably, we observed numerous phenotypic transitions (resistant to susceptible or *vice versa*) unrelated to non-synonymous changes in these genes, including four transitions for fluconazole, four for voriconazole, one for itraconazole and two for isavuconazole (**Fig. 3**). Similarly, we found no clear association between variants in previously expected genes and resistance to echinocandins (*FKS1*, *FKS2* and *ERG3*^23,61,62,66–70^, **Extended Data Fig. 7a**), amphotericin B (*ERG2, ERG3, ERG5, ERG6* and *ERG25*^23,71,72^, **Extended Data Fig. 7b**) nor 5-flucytosine (*FCY1, FCY2, FUR1* and *CHS7*^73^). For echinocandins and amphotericin B this might be because our “resistance” phenotype reflects reduced susceptibility rather than clinical resistance (see **Methods**, **Extended Data Fig. 4**). For 5-flucytosine, mechanisms inferred for this drug in other species (e.g. *Candida albicans*) may not apply to *C. parapsilosis*.

Taken together, these results indicate that previously implicated genes offer only a partial understanding of resistance mechanisms. Additional limitations of this hypothesis-driven approach is its inability to identify novel genetic determinants and the lack of rigorous statistical support, as it ignores the possibility that some of the mutations may be associated with resistance by chance.

### Convergence-based GWAS reveal the landscape of genotype-phenotype relationships

To systematically investigate genetic changes underlying relevant traits, we performed a convergence-based GWAS across all phenotypes, which uses ancestral state reconstruction (ASR) to associate evolutionary transitions in phenotypes and genetic variants^9,74,75^. For this, we developed a pipeline similar to that in ^9,76^, which considers variants individually or in groups, to account for allelic heterogeneity and equivalent functional effects at the gene, domain or pathway level (**Fig. 4a**). To focus on high confidence associations, we only analysed 27 of 61 traits with at least three phenotype transitions, and used conservative filters (see **Methods, Extended Data Fig. 8, Supplementary Table 2, 3**).

**Fig. 4.**
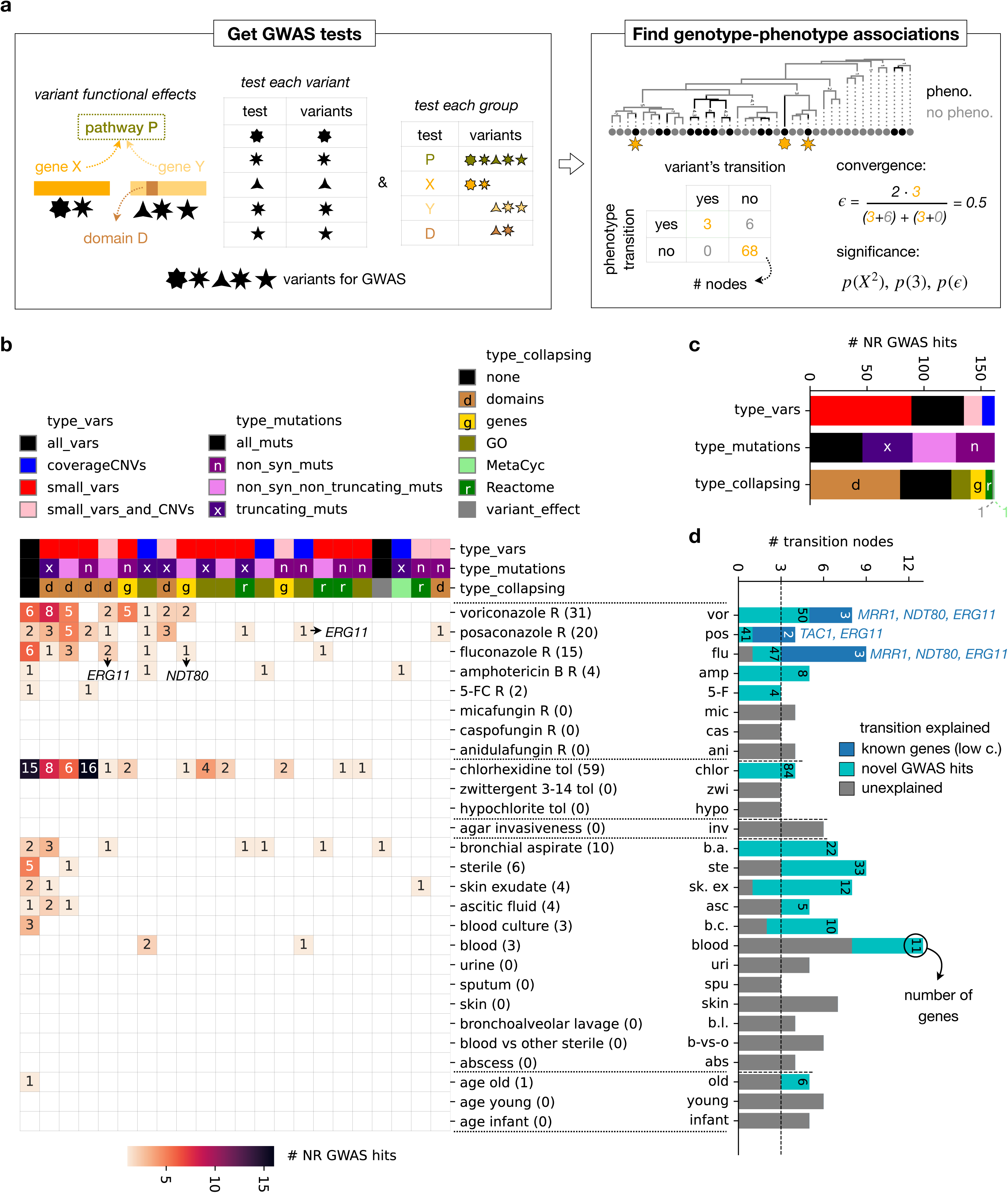
Convergence-based GWAS to identify the genetic determinants of each trait. (a) Overview of the GWAS pipeline applied. Flowchart adapted from ^9^ (see **Main text** and **Methods**). (b) Heatmap showing the number of high-confidence non-redundant (NR) GWAS hits found in each phenotype (rows), by each collapsing strategy (columns). The phenotypes are sorted by the number of hits, and also grouped by phenotype type. Also, the columns are sorted by number of hits. The arrows indicate hits including known azole resistance genes (see Fig. 3) (c) Barplot showing the numbers of hits across all phenotypes that used each collapsing strategy. (d) Barplot showing how different phenotype transitions are explained. For each transition, we evaluated whether it could be explained by change in the expected genes for azole resistance (blue) or, if this was not the case, whether it could be explained by any of the GWAS hits of the phenotype (cyan). The gray bar reflects the number of transitions that are not explainable by any of the latter mechanisms. Also, note that we defined as ‘transitions explained by expected genes’ those that had a genotype-phenotype association across any of 18,144 filtering parameter combinations (see **Methods**), so we had high confidence about the transitions unexplained by these genes. In addition, the numbers indicate the numbers of genes associated with the GWAS hits / known genes in each phenotype.

We identified high-confidence associations for 13 traits (**Fig. 4b, Supplementary Table 4**, see also **Supplementary Table 5** for results with less stringent filtering). These included not only resistance phenotypes (towards 5-FC (2 hits), amphotericin B (4), fluconazole (15), posaconazole (20) and voriconazole (31)), but also chlorhexidine tolerance (59), and patient-related traits (age old (1) and various isolation sources (blood (3), blood culture (3), ascitic fluid (4), skin exudate (4), sterile (6), and bronchial aspirate (10))). To assess genetic architecture we further inspected the type of hits obtained (**Fig. 4c**). This suggested mostly independent trait acquisitions except for fluconazole resistance and chlorhexidine tolerance, which may have also been shaped by (para)sexual recombination (**Supplementary Results**, **Fig. 4c**, **Extended Data Fig. 9, 10, 11**), as observed in other *Candida* species^9^.

We further assessed the explanatory power of these hits by visualizing the fraction of phenotypic transitions associated with i) expected genes (see above and **Methods**), or ii) high-confidence GWAS hits (**Fig. 4d**). Most hits explained transitions not attributable to expected genes, supporting that our approach uncovered novel mechanisms. Nevertheless, a large fraction of phenotypic transitions remained unexplained by either approach (**Fig. 4d**). While this unexplainability could be attributed to a lack of genetic heritability for some phenotypes (e.g. isolation sources), this is hardly the case for others (e.g. drug resistance or agar invasiveness). Additionally, our stringent GWAS thresholds and the limited sample size might limit statistical power, as we potentially missed known associations for *MRR1, TAC1* and *NDT80* in azole resistance (**Methods, Fig. 4b, Extended Data Fig. 8**). Finally, the numerous, often independent, hits found for many phenotypes also suggest polygenic effects, which univariate GWAS cannot fully resolve. Thus, GWAS likely presented an incomplete picture of genotype-phenotype relationships, necessitating further analyses.

### Machine learning classifiers uncover genotype-phenotype relationships

To further understand genotype-phenotype associations, including genetic interactions and polygenic effects, we developed machine learning (ML) classifiers to predict phenotypes from genetic variants. We focused on 13 phenotypes for which we had five or more sharp transitions across the strain phylogeny. These include body sources (skin exudate, urine, bronchial aspirate, blood, sterile, and blood vs other sterile), antifungal resistance (towards micafungin, fluconazole and voriconazole), patient age groups (infant, young, old) and agar invasiveness. For each phenotype, we built accurate and generalizable ML classifiers by evaluating various feature selection approaches, models and model hyperparameters, totalling 5,940-216,864 parameter combinations (‘train parameters’) for each phenotype. For each model, we used 80% of the data for training and evaluation (in 75% / 25% splits), independently testing performance on the remaining 20% (**Fig. 5a**).

**Fig. 5.**
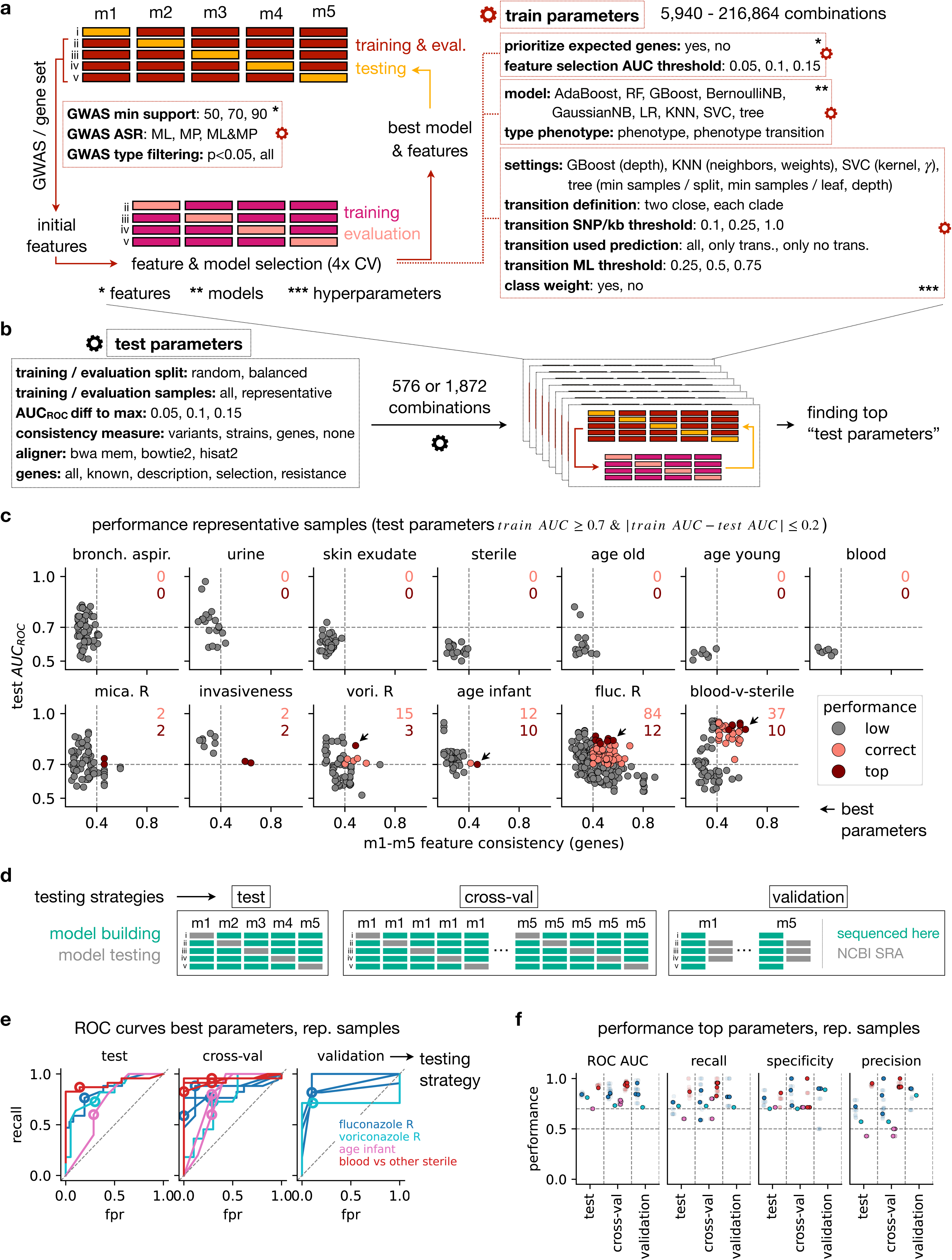
Building machine-learning classifiers for four phenotypes. (a) Schematic representation of the process used to build ML classifiers for a given phenotype. We tried to build ML classifiers for 13/61 phenotypes (see Fig. 2), for which we could find >=5 phenotype-no phenotype transitions by all aligners. To reduce overfit performance estimates, we split the data into ∼80% for training & evaluation, and ∼20% for testing, in a phylogenetically-balanced way (see **Extended Data Fig. 6**). Note that the approximate 80-20 splits is due to the fact that for different phenotypes the phylogenetically balanced splitting would result in slightly different percentages. We used the training & evaluation set to select optimal features, models and model hyperparameters, in two steps. First, we narrowed down the initial set of features (variants, genes, domains and/or pathways) to only include those potentially related to each phenotype. We did this by either i) applying a convergence GWAS pipeline to pinpoint those associated with the phenotype or ii) only analyzing those affecting genes that have been previously related to each phenotype (see **Methods**). Second, we applied 4-fold cross-validation (CV) to select features (with forward feature selection), models (e.g. logistic regression, random forest…) and model parameters (e.g. tree depth). Note that we tried to predict either i) directly the phenotype or ii) phenotype transitions between close strains (see **Methods**); which are related to the model / hyperparameter selection. We then selected the ‘best model and features’ as the one that had a high ROC AUC and high consistency of selected features across 5 different random train/evaluation CV runs. Finally, we tested the performance of this best model on the testing data for independent model evaluation. Given our small sample size, we ran this process 5 times, obtaining 5 models (m1, m2, m3, m4 and m5) for each train/test split in each phenotype. (b) To also check the effect of various parameters on final model performance we assessed the effect of varying i) the type of splitting the training & evaluation samples, ii) the training sample set, iii) the way to select the best models (e.g. what constitutes a high ROC AUC, or how to measure feature consistency), iv) the aligner, and v) the initial gene set evaluated when no GWAS was used for initial feature filtering (e.g. genes correlated to resistance or recent adaptation in other *Candida* species^9^ and/ or genes with functions potentially related to each phenotype). Note that all these are parameters that cannot be assessed using the train & evaluation set. We tried different numbers of parameter combinations for different types of phenotypes (i.e. 1872 for resistance traits, as these required testing more gene sets, and 576 for the others). Thus, we ran the pipeline from (a) 1872 or 576 times for each phenotype, varying these parameters. (c) For each of the test parameters with no clear sign of overfitting or underfitting (ROC AUC >= 0.7 on the training data and similar AUC in training and testing), the ROC AUC on test data (y axis) vs the consistency of the features yielded by each of the m1-m5 models (x axis, see **Methods**). The colors represent whether the parameters have a correct performance (test AUC >= 0.7 and consistency >= 0.4), and whether they are among the top performing parameters (they have an AUC >= max AUC - 0.05, and a consistency in the top ten of all correct models). The arrows point to the best performing parameters for the four phenotypes for which we consider to have sufficient numbers (>=10) of correct parameters (“voriconazole resistance”, “age infant”, “fluconazole resistance” and “blood vs other sterile”). (d) To assess model performance we used different testing strategies, each based on a distinct set of data for model building (green) and testing (gray). For instance, the ‘test’ strategy consisted in building models m1-m5 on the training & evaluation set data, and aggregating the results on the testing data from different models, which is equivalent to the y axis from (c). We only considered models that could be built (e.g. had AUC ROC>=0.7 and >1 predictor), ignoring cases with >1 model that could not be built. Also, to get an estimate of the performance across models we got ‘cross-val’ performance estimates for each model m1-m5 using CV on all data. We only considered models that had a ROC AUC close to the maximum ROC AUC across all m1-m5 (AUC >= max AUC - 0.05). Finally, to get an additional estimate on unseen data we considered independently-generated datasets, available in NCBI SRA. We assessed the performance of each model m1-m5, built on all the data sequenced here, and evaluated the performance on the independently generated samples. (e) Receiver operating characteristic (ROC) curves related to the best parameters from (d), for each of the testing strategies (see (c)). For ‘cross-val’ and ‘validation’ each line corresponds to a different model m1-m5. Note that ‘validation’ was only calculated for fluconazole and voriconazole resistance because we could not find sufficient public data to validate the other phenotypes. The circle represents an optimal discretization threshold, selected based on the ‘test’ curve for ‘test’ metrics, and on the ‘cross-val’ curve for ‘cross-val’ and ‘validation’ metrics. In all cases, we selected a threshold that would maximize recall while having a fpr <= 0.3. (f) Performance metrics for all top models. The points without transparency reflect the results of the best models. Note that for recall, specificity and precision these are based on a threshold selected as defined in (e).

Given the specific challenges of our dataset (see **Supplementary Results**) we incorporated several non-standard steps. As in the GWAS, we used both single and collapsed variant groups as features to address allelic heterogeneity (**Fig. 4a, Methods**). To ensure robust testing, we ran the process on five different train / test splits, generating five models (m1-m5). Also, to avoid misleading performance estimates due to sampling bias, we used a set of phylogenetically representative samples (**Extended Data Fig. 6a**) for final testing. Furthermore, to prevent models from predicting phylogeny instead of phenotype (an issue likely affecting previous *Candida* ML studies^46,48^), we defined train / test splits based on a phylogenetically-balanced approach, where clades used in training/evaluation and testing are different (**Extended Data Fig. 6b**). Finally, to increase the modelling sample size and minimize the number of possible features we explored predicting phenotypes from the classification of phenotype transitions between pairs of closely related strains.

Given the scarcity of similar studies in *Candida*, we explored different parameter choices (**Fig. 5b**), providing insights for future ML studies in these pathogens. We tested varying read aligners, feature selection approaches, training datasets (all vs representative isolates), the method for partitioning train and evaluation sets (randomly or phylogenetically balanced), performance metrics, and the criteria to select optimal ‘train parameters’ (**Methods**). For 576 or 1872 (depending on the phenotype) combinations of such ‘test parameters’ we evaluated model performance (Receiver operating characteristic (ROC) curve AUC on test data) and model feature consistency (feature overlap across m1-m5 models). For four phenotypes (fluconazole and voriconazole resistance, blood vs other sterile, and age infant), we found more than 10 (12-84) test parameters yielding acceptable predictive capacity and consistency (AUC >= 0.7, consistency >= 0.4) (**Fig. 5c**), demonstrating significant predictive performance. While this may result from assessing multiple parameters, we observed large performance differences among phenotypes (**Fig. 5c**) unrelated to sample size differences (**Fig. 2**), suggesting that high performances reflect models capturing true genotype-phenotype relationships.

To further validate our classifiers we analyzed ‘top parameters’ (i.e. those with an AUC >= max AUC - 0.05 and consistency within top 10) for these four traits (**Fig. 5c**), using various ‘testing strategies’ (**Fig. 5d**). First, two strategies involved training / testing on the datasets generated here, but either obtaining aggregate performance metrics for each parameter (‘test’ strategy) or one metric for each ‘top’ model m1-m5 (‘cross-val’ strategy). Furthermore, the ‘validation’ strategy involved training on all of our data, and testing on independent publicly available datasets overlapping most of our resistant clades (obtained for voriconazole and fluconazole resistance^33,37,38,77–79^, see **Extended Data Fig. 12**). ROC curves for all strategies suggested a good balance between specificity and recall in all models yielded by ‘top parameters’ (**Fig. 5e,f**). We evaluated recall, specificity, and precision using thresholds that balanced recall and specificity (maximum recall at fpr <= 0.3), and found that most models achieved a good balance between specificity and recall with slightly smaller precision (except ‘blood vs other sterile’) (**Fig. 5f**). In a clinical setting this trade-off might be acceptable, as having false negatives (e.g. missing a drug resistant strain) might be less desirable than having false positives (e.g. wrongly assuming resistance in a susceptible strain). Importantly, the accuracy of these classifiers could improve as additional data becomes available.

To provide interpretability and infer evolutionary mechanisms, we analyzed variant patterns and genes related to predictive features for the ‘top parameters’ (**Supplementary Results, Fig. 6a, Extended Data Fig. 13**). This suggested that fluconazole resistance is related to (mostly non-synonymous) changes in seven genes (*ERG11*, *Scer_MOH1* (i.e. ortholog of *Saccharomyces cerevisiae MOH1*)*, CDR1B, Scer_ATP10, MRR1, Scer_ERG24* and *CPAR2_204120*) which collectively explain all phenotypic transitions (**Fig. 6b,c, Extended Data Fig. 14**, **15**). Notably, we observe that variants in the novel genes *CDR1B*, *Scer_MOH1* and *Scer_ERG24* are necessary to explain transitions unexplained by changes *ERG11*, *TAC1*, *MRR1* or *NDT80* (**Fig. 3**, **Fig. 6b,c**), showing our model’s utility in leading to new results. Furthermore, our findings confirm the previously-proposed relevance of *CDR1B* in azole resistance^43^. Conversely, voriconazole resistance can be attributed to non-synonymous variants in *ERG11* and *STP4*, combined with regulatory changes around *Scer_CLU1* (**Extended Data Fig. 16**). Additionally, we infer that the predisposition towards infecting infants is influenced by combinations of regulatory variants around *Scer_PMI40* and non-synonymous changes in *ALS11* or *RBT1* (**Extended Data Fig. 17**). Finally, we find that the predisposition towards infecting non-sterile compartments outside the bloodstream is modulated by regulatory changes around *Scer_MDS3* combined with non-synonymous changes in *ALS11* or *ALS7* (**Extended Data Fig. 18**).

**Fig. 6.**
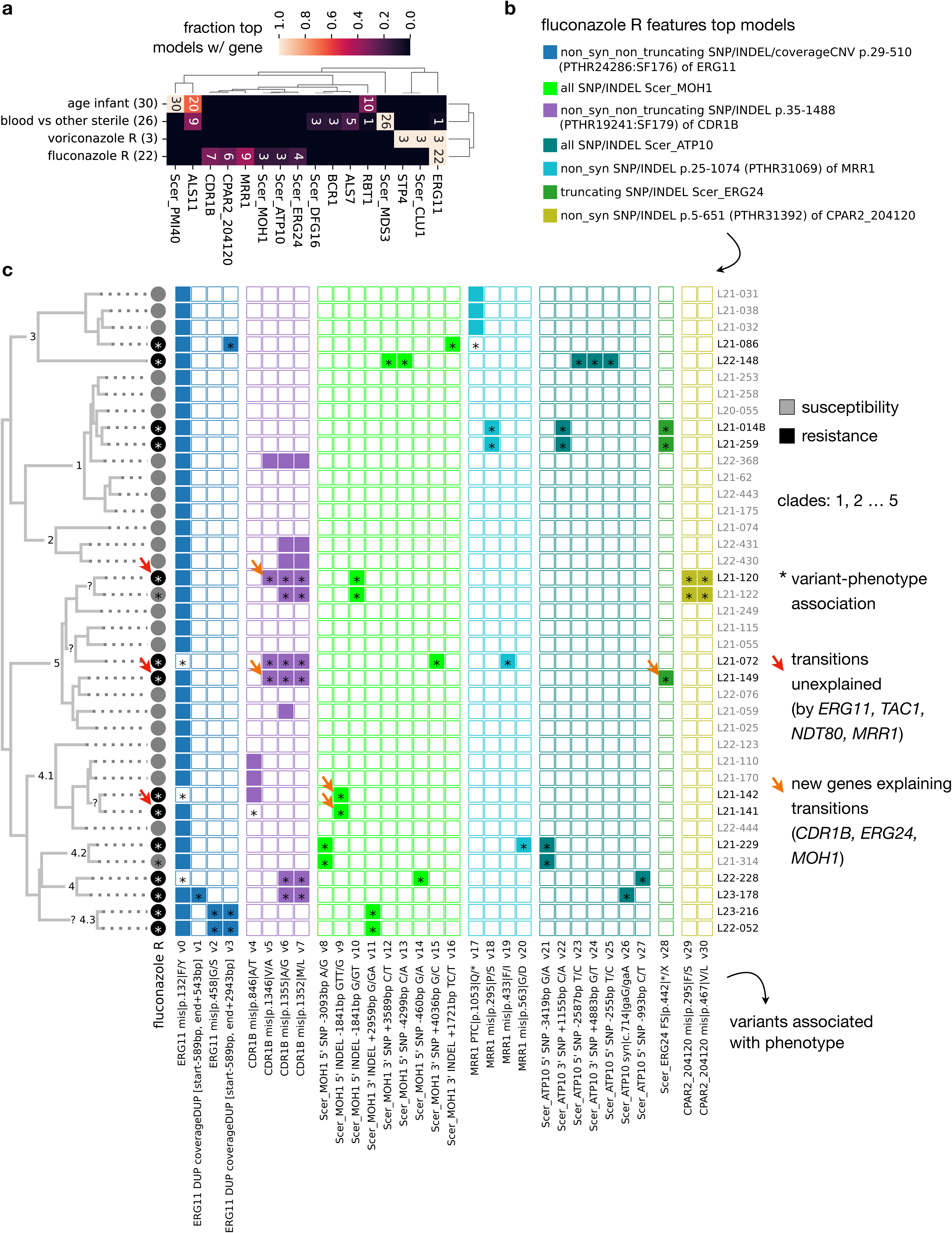
Features selected by the top models. (a) Heatmap showing the fraction of top models, for each phenotype, with some feature mapped to each gene. Only genes found in >=10% of models are shown. (b) Most common features used for each of the genes used for modelling fluconazole resistance in >=3 top models. (c) Presence / absence pattern of variants related to each feature, shown as in Fig. 3. Only variants appearing with the phenotype, or in nodes that have the phenotype, are shown. The orange arrows show instances where a resistance event cannot be explained by previously-known genes, but instead it is explained by some of the model features. See **Extended Data Fig. 16, 17** and **18** to see equivalent plots for the other three phenotypes.

Taken together, these results suggest that we can predict from sequence data fluconazole / voriconazole resistance, the capacity to invade non-sterile compartments different from blood (mostly ascitic and cerebrospinal fluids, **Extended Data Fig. 5b**) and a predisposition to infect infants with reasonably high confidence. Given the potential clinical applications, our findings constitute a major milestone in the study of adaptation mechanisms of *Candida* pathogens. Furthermore, our thorough parameter analysis informs on optimal strategies for building genotype-phenotype ML classifiers in *Candida* and other clonally-reproducing species (see **Supplementary Results, Extended Data Fig. 13, 19, 20**). Finally, our analysis of predictive features enables interpretability of classifier behavior, suggesting novel determinants of resistance and virulence that deserve further attention. The capacity to predict patient age and source compartment from strain genetics is particularly relevant, but it will require additional validation, as our limited metadata cannot discard that isolates affecting infants or non-sterile compartments may also be able to infect other age groups or body parts.

### GWAS and ML results reveal the drivers of phenotypic variation

Despite overlapping results on certain key genes (*ERG11, CDR1B, Scer_ERG24, MRR1, Scer_CLU1* and *Scer_MDS3*), we found that GWAS and ML approaches yielded mostly non-overlapping sets of genes (**Extended Data Fig. 21**), suggesting they provide complementary rather than redundant insights. Collectively, the analysis of expected genes, convergence GWAS and ML features suggested a total of 298 genes with variants associated to any of the 15 considered phenotypes (**Fig. 3, 5, Supplementary Table 4**). From those, we prioritized a subset of 79 candidate gene-phenotype mappings, including the (most likely) relevant genes for each phenotype (see **Methods, Fig. 7, Extended Data Fig. 22, Supplementary Table 6**). While we discuss all these mappings in **Supplementary Information**, we next provide a summary of the functions of these genes, which include mostly novel findings showing the complexity and polygenic nature of these genotype-phenotype relationships.

**Fig. 7.**
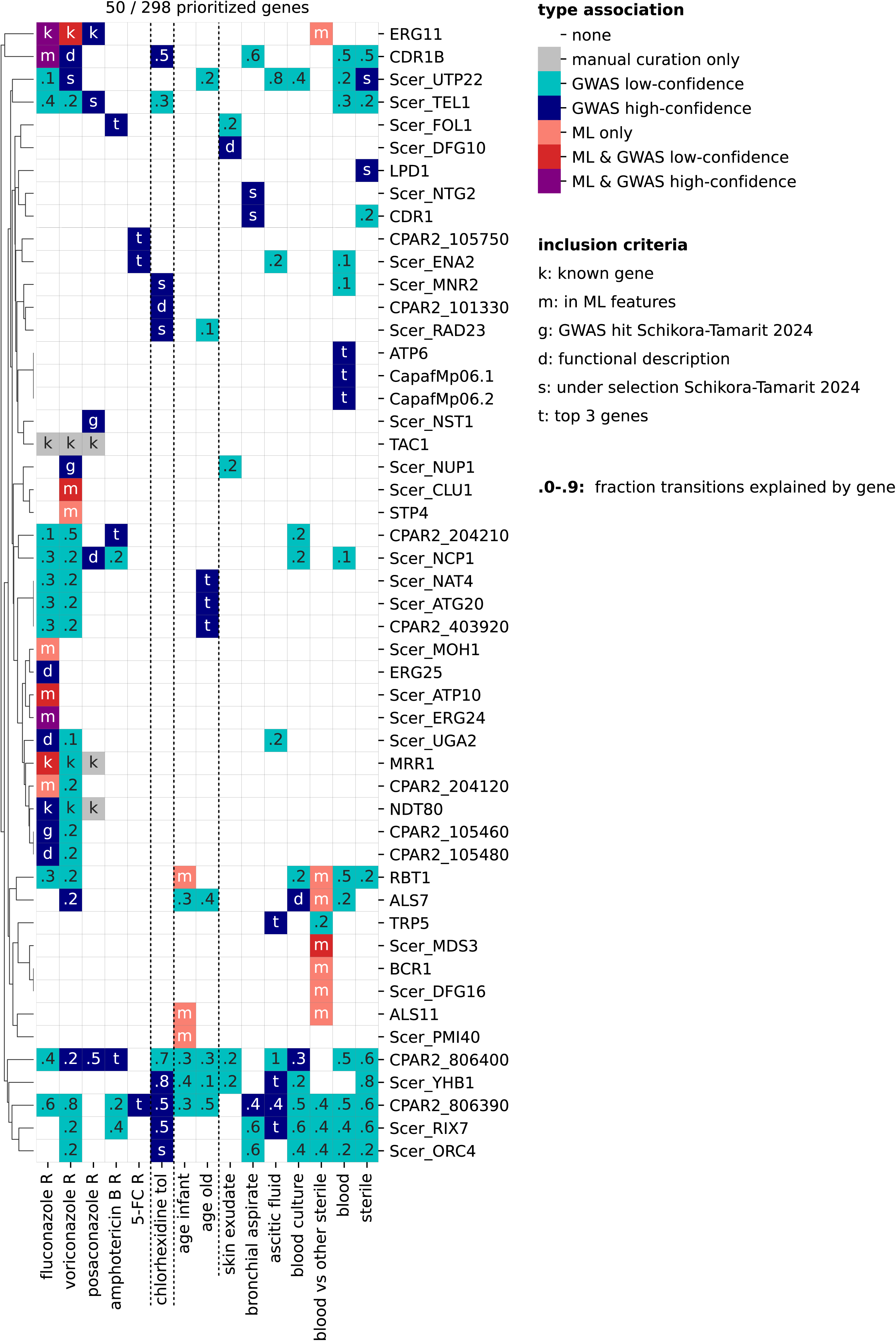
Prioritized functional genes. Genes prioritized among those that are expected (known azole resistance genes), or are related to GWAS or ML hits. The color reflects the source of the associations, and the symbols indicate the inclusion criteria that made a gene be prioritized for a given phenotype. Also, the floats indicate instances where a gene was not prioritized but had associated GWAS hits, and the float indicates the fraction of phenotype transitions that can be associated with such genes. See **Methods** section ‘Integration of ML and GWAS results’ for more information regarding the prioritization strategy. Note that **Extended Data Fig. 22** contains the same plot but with descriptions.

We find seven pan-azole resistance genes possibly related to impaired drug binding and/or titration (*ERG11*), regulation of azole efflux (*NDT80*, *TAC1* and *MRR1*), ergosterol biosynthesis (*Scer_NCP1*), biofilm formation and DNA-damage stress responses (*Scer_TEL1*), and unknown functions (*CPAR2_806400*). Additionally, we identified seven genes associated to only fluconazole and voriconazole resistance, possibly related to drug efflux (*CDR1B*), rRNA processing and tRNA export (*Scer_UTP22*), aminoacid, energy and xenobiotic metabolism (*Scer_UGA2, CPAR2_105460, CPAR2_105480* and *CPAR2_204210*), and protein glycosylation (*CPAR2_204120*). Furthermore, we find eight genes associated to resistance towards only one azole, possibly related to mitochondrial-activated drug efflux (*Scer_ATP10*), vacuolar transport (*Scer_MOH1*), ergosterol biosynthesis (*Scer_ERG24* and *ERG25*), osmotic stress responses (*Scer_CLU1* and *Scer_NST1*), transcriptional regulation of azole responses (*STP4*) and nuclear pore activity (*Scer_NUP1*).

We find that amphotericin B reduced susceptibility is related to four genes involved in ergosterol biosynthesis (*Scer_NCP1*), xenobiotic metabolism (*CPAR2_204210*), tetrahydrofolate biosynthesis (*Scer_FOL1*) and unknown function (*CPAR2_806400*). With the exception of *Scer_FOL1,* these genes were also related to azole resistance, which suggests common resistance mechanisms between the two drug families, possibly due to common effects on ergosterol biosynthesis. In addition, 5-flucytosine resistance is associated with three genes involved in regulation of carbohydrate metabolism (*CPAR2_105750*), transmembrane ion transport (*Scer_ENA2*) and unknown functions (*CPAR2_806390*). Furthermore, chlorhexidine tolerance is associated with changes in putative stress responses (*CPAR2_101330* and *Scer_RAD23*), DNA replication (*Scer_ORC4*), magnesium homeostasis (*Scer_MNR2*) and drug efflux (*CDR1B*).

Regarding clinical phenotypes, we find that predisposition towards infecting infants is associated to genes related to mannoprotein biosynthesis (*Scer_PMI40*), biofilm formation (*Scer_PMI40, RBT1*), invasion (*RBT1, ALS11*), hypoxic growth (*RBT1*) and adhesion to human cells (*ALS11*). Conversely, predisposition towards infecting older patients is associated with genes involved in protein glycosylation (*Scer_NAT4*), autophagy (*Scer_ATG20*), biofilm formation (*CPAR2_403920*) and morphogenesis (*CPAR2_403920*). Although the sets of genes related to each age group are non-overlapping, they affect certain similar functions (biofilm formation, protein glycosylation and adhesion), suggesting common properties of strains that infect such particularly susceptible patients.

Finally, the genes modulating predisposition towards infecting specific body compartments are possibly involved in pseudohyphal growth (*Scer_MDS3, Scer_DFG10*), host adhesion (*ALS7, ALS11*), biofilm formation (*Scer_MDS3, ALS7, Scer_NTG2, CDR1*), protein glycosylation (*Scer_DFG10*), DNA-damage responses (*Scer_NTG2*), xenobiotic transmembrane transport (*CDR1*), rRNA processing and tRNA export (*Scer_UTP22*) and antigenic properties (*LPD1*). These are all virulence-related functions that may be modulated through mutations, leading to selectivity for specific compartments in certain isolates.

Beyond the functional insights, the analysis of the involved variants suggested relevant trends. We find that 24 of the 79 prioritized gene-phenotype mappings involve only upstream, downstream or CDS synonymous variants (**Supplementary Table 6**), suggesting that non-coding and synonymous variants, perhaps through altering gene regulation, may play a significant role in shaping these clinically-relevant phenotypes. Additionally, 20 of 79 gene-phenotype mappings related to CNVs and SVs (**Supplementary Table 6**), showing the importance of considering these complex, often overlooked, variants. These findings highlight our approach’s ability for identifying all relevant variants in genotype-phenotype analyses. Also, for most hits we could find previous confirmatory evidence in other fungal species, showing the robustness of our approach and providing interesting hypotheses for further research (see **Supplementary Information**).

## CONCLUSIONS

Despite increasing clinical relevance of *Candida parapsilosis*, the genetic mechanisms driving key traits remain poorly studied in comparison to other *Candida* species. To address this gap we exploited the combined power of evolutionary genomics, Genome-Wide Association Studies (GWAS), and innovative machine learning (ML) tools. The effectiveness of these approaches relies on the availability of genetic and phenotypic data for a large and diverse set of isolates. Hence, we generated an unprecedentedly large and comprehensive collection of *C. parapsilosis* genomes and clinically-relevant phenotypes for a multy-hospital outbreak in Spain, establishing a valuable resource for the field. The analysis of this newly derived dataset in combination with publicly available genomes uncovered four novel sub-clades, and refined the phylogeography of *C. parapsilosis*, highlighting both global and local spreading.

Assessment of previously established resistance mechanisms (the only approach used in most previous studies) confirmed the crucial role of *ERG11, TAC1, MRR1*, and *NDT80* in azole resistance. However, numerous azole resistance transitions in our dataset remained unexplained, suggesting the existence of additional, uncharacterized mechanisms. Similarly, for echinocandins, amphotericin B, and 5-flucytosine, previously established mechanisms failed to adequately explain the breadth of observed phenotypic variation, underscoring the limitations of such hypothesis-driven approaches.

To obtain further insights, including for clinically-relevant traits with no candidate genes, we performed a convergence-based GWAS, which identified both known and novel genetic drivers of various traits. While our GWAS provided significant insights, our rigorous filtering strategy ensuring high-confidence associations, resulted in limited explanatory power. The persistence of unexplained transitions, coupled with the likely polygenic nature of many traits, highlights the limitations of univariate approaches in fully capturing the complex genetic architecture of *C. parapsilosis*.

Finally, we extensively explored the use of ML classifiers as an alternative genotype-phenotype discovery approach. We obtained highly accurate ML classifiers for four key phenotypes: fluconazole resistance, voriconazole resistance, predisposition to infect non-blood sterile compartments, and predisposition to infect infants. These classifiers represent a major step towards developing DNA-based profiling approaches to predict virulence and resistance traits, offering promising avenues for future clinical applications, including more precise diagnostics and personalized therapies. Furthermore, our comprehensive benchmarking of ML parameters and strategies provides crucial information for building robust ML models in *Candida* pathogens.

The use of interpretable ML models allowed us to elucidate underlying mechanisms of virulence and resistance. Notably, for fluconazole resistance, our models identified novel genetic determinants that collectively explain all phenotypic transitions previously unexplainable by known genes. This showcases the power of evolutionary informed multivariate analyses as the one proposed here. Although ML approaches have been previously used to predict drug resistance from genome data in *Candida* pathogens^46,48^, we propose here an innovative and rigorous implementation that maximizes the discovery of causal relationships through phylogeny-aware data splits, and enhances predictive performance by analyzing phenotypic transitions instead of phenotypes.

Altogether, our study provides relevant insights for understanding genotype-phenotype relationships, not only within the broader field of *Candida* pathogens but also for other clonally-reproducing species. Importantly, we have not only characterized genotype-phenotype relationships for drug resistance, as commonly done in earlier analysis, but also for non-drug resistance traits, identifying, for the first time, variants that may predispose or facilitate *C. parapsilosis* strains to infect specific body parts and patient age groups. To facilitate a broader adoption of our approach, we share the results of our extensive benchmarking to aid in the development of similar ML approaches for other traits, as well as easy-to-use and install toolkits for end-to-end GWAS and ML analyses.

In conclusion, our work significantly advances the understanding of genetic determinants underlying clinically relevant traits in *C. parapsilosis*, providing a robust framework and valuable resources for future research and clinical translation.

## MATERIALS AND METHODS

### Whole genome sequencing

We sequenced 233 representative strains from a 2019-2023 multi-hospital outbreak in Spain described in a recent study, which generated microsatellite typing and *ERG11* sequencing data for them^51^. We selected representative isolates from all the different genotypes based on microsatellite typing, covering a variety of *ERG11* mutations, body isolation sources and geographical origin. For DNA isolation, cells were cultivated in Sabouraud liquid medium at 30°C with shaking for 24h, and after centrifugation, the pellets suspended in PBS and disrupted with FastPrep (MP Biomedicals) and glass beads with three cycles of 20 seconds of rupture and 20 seconds of incubation in ice. Then, phenol-chloroform-isomyl alcohol (25:24:1) was added and centrifuged and the DNA in the supernatant was precipitated with 70% Ethanol. Finally, the pellet was dissolved in distilled water and treated with RNase. Concentration and purity of the DNA was obtained with Nanodrop (Thermo Scientific) and Quantifluor and Quantus Fluorimeter (Promega). For whole genome sequencing, libraries were prepared using DNA Prep. Tagmentation kit following the manufactureŕs protocol. Sequencing was done in a NovaSeq sequencer (NovaSeq 6000 SP Reagent kit v1.5, 300 cycles, double stranded).

We sequenced the 233 strains in three sequencing runs (‘OSCAR01’, ‘OSCAR02’ and ‘OSCAR03’). Due to lower coverage in the first run (‘OSCAR01’), we sequenced twice the samples L21-060 and L21-067 (in both projects ‘OSCAR01’ and ‘OSCAR02’), so we have a total of 235 strains sequenced here. However, the strain tree (see below) indicates that all of these samples belong to a single unresolved polytomy (when requiring node support > 90). Thus, we conclude that both runs were equivalent, and the runs from OSCAR02 were used for all analyses below for these two strains (except tree reconstruction).

### Strain phenotyping

Antifungal susceptibility in liquid medium was obtained from ^51^, generated with the EUCAST 7.4 protocol, for the following antifungals: amphotericin B, 5-flucytosine, fluconazole, itraconazole, voriconazole, posaconazole, isavuconazole, caspofungin, micafungin and anidulafungin. The Minimal inhibitory concentration (MIC) was the minimal concentration that inhibited 50% of growth compared to the control well, except for amphotericin B (90%). Similarly, we obtained various phenotypes from a previous study on our collection^54^, including percentage of viability in various disinfectants (ethanol, sodium hypochlorite, H_2_O_2_, Surfanios Premium, chlorhexidine digluconate, Zwittergent 3–14 detergent), biofilm formation (high-biofilm-forming, low-biofilm-forming and non-biofilm-forming), morphology (yeast or pseudohyphae), agar invasiveness (high-invasiveness, low-invasiveness or non-invasive) and microfluidics behavior (adherence, formation of big clumps, or formation of little clumps).

In addition, to infer antifungal susceptibility in solid medium we performed E-tests. In brief, *C. parapsilosis* cells were placed in Sabouraud plates and incubated overnight at 30 oC. Then, the cells were suspended at 10^6^ cells/mL, and 100 uL were spread on RPMI+2% glucose buffered at pH 7 with MOPS. E-test strips (Biomerieux) were placed on the middle of the plate. The plates were incubated at 35 °C. MIC was evaluated at 24 and 48 h as the concentration of fluconazole in the strip that crossed the inhibition halo formed in the plate. Heteroresistance was defined as the appearance of colonies inside the inhibition halo.

### Code and software environments

The code necessary to perform all analyses described in this work is available in the GitHub repository https://github.com/Gabaldonlab/Cparapsilosis_GenoPheno (latest version by the time of publication). In brief, we used the script run_pipelines.py to generate processed datasets through launching HPC jobs and figures_paper.py to generate plots and tables. Additionally, we ran various different scripts for specific analysis (get_genetic_distances_one_sample.py, downsample_reads.py and get_mappability_file_whole_genome.py), as described below. These scripts require Cparapsilosis_popgen_functions.py, a module comprising all necessary python functions. Below we refer to specific functions as CPfun.<FUNCTION name> which, unless specified otherwise, rely on custom python and pandas code. We ran all of this code on the conda environment Cparapsilosis_popgen_env, defined in the file Cparapsilosis_popgen_env.yml of the repository, which includes matplotlib (3.8.0)^80^, multiqc (1.17)^81^, numpy (1.26.0)^82^, python (3.9.7)^83^, sra-tools (3.0.5)^84^, scipy (1.11.3)^85^, seaborn (0.13)^86^, biopython (1.78)^87^, pandas (2.1.1)^88^, ete3 (3.1.3)^89^, jellyfish (1.0.1)^90^, openpyxl (3.0.10)^91^, gffread (0.12.1)^92^ and sciuality control, our final dataset comprised 365 high-quality, high-coverage genomes (189 newly sequenced, 176 from SRA), from 13 countries across North America, Akit-learn (1.5)^93^.

Additionally, we used other pipelines and conda environments, described next. For the variant analyses we used the published tool PerSVade (latest version by 24 November 2024)^53^, available at https://github.com/Gabaldonlab/perSVade. For some analysis steps we used the conda environments perSVade_env and perSVade_env_RepeatMasker_env, which were generated with the traditional PerSVade installation described in https://github.com/Gabaldonlab/perSVade/wiki/2.-Installation. The perSVade_env was used to run various perSVade modules, the perSVade script get_trimmed_reads_for_srr.py and our script get_mappability_file_whole_genome.py (see below). Also, for some specific analyses we used certain tools within this environment, including unpigz (2.4)^94^, bedtools (2.29.0)^95^, bedmap of the bedops suite (2.4.39)^96^ and samtools (1.9)^97^. Furthermore, note that perSVade’s modules internally use various additional conda environments, generated as explained in the aforementioned Installation page. In addition, the perSVade_env_RepeatMasker_env was used to run RepeatMasker (4.0.9_p2)^98^.

For tree reconstruction we used the tree_from_SNPs pipeline, used in ^9^, and available at https://github.com/Gabaldonlab/tree_from_SNPs (latest version by July 3 2025). We ran this pipeline on the tree_from_SNPs_env conda environment available in this repository, which includes r-phytools (0.7_90)^99^, r-base (4.1.1)^100^, python (3.6.11), biopython (1.78), bedops (2.4.41), pandas (1.1.1), mosdepth (0.3.1)^101^, ete3 (3.1.2), iqtree (2.1.2)^102^, picard (2.18.26)^103^, gatk4 (4.1.2.0)^104^, snp-sites (2.5.1)^105^, bedtools (2.29.0), datamash (1.1.0)^106^ and r-argparser (0.7)^107^.

For the GWAS and ML parts we developed the ancestral_GWAS toolkit, inspired by our previous work in *Candida*^9^, available at https://github.com/Gabaldonlab/ancestral_GWAS_toolkit (latest version by the time of publication). We ran the scripts included in this toolkit on the ancestral_GWAS_env conda environment available in this repository, which includes matplotlib (3.5.1), pip (21.3.1), python (3.10.0), seaborn (0.11.0), scikit-learn (1.5.1), biopython (1.79), ete3 (3.1.3), pandas (1.3.5), scipy (1.7.3), statsmodels (0.13.1)^108^, openpyxl (3.0.10), psutil (5.8.0)^109^, bedtools (2.31), bedops (2.4.41), samtools (1.6), gffread (0.12.1), goatools (1.1.12)^110^, pastml (1.9.34)^111^ and pythoncyc (2.0.2)^112^. Beyond these pipelines used for many steps in our analysis, we also ran other tools for more specific steps. First, for the quality control of the sequenced genomes we used the Kraken2 (2.1.1)^113^ and ktImportTaxonomy from Krona (2.7.1)^114^ tools installed within the MeTAline pipeline conda environment (meTAline), available at https://github.com/Gabaldonlab/meTAline. Second, for fetching metadata from the NCBI SRA database^52^ we used epost and efetch from entrez-direct (13.9)^115^, within the conda environment Candida_mine_env used in our previous work^9^, available at https://github.com/Gabaldonlab/Candida_Selection_DrugResistance (latest version by July 3 2025). Third, for running protein annotations we used Interproscan (5.52-86.0)^116^ within the conda environment InterProScan_env from the same ‘Candida_Selection_DrugResistance’ repository. Fourth, for processing MetaCyc gene annotations we used Pathway Tools (25.0)^117^.

In summary, we used various conda environments (Cparapsilosis_popgen_env, perSVade_env, perSVade_env_RepeatMasker_env, tree_from_SNPs_env, ancestral_GWAS_env, meTAline, Candida_mine_env and InterProScan_env) to run different parts of the analysis. In the sections below we refer to the specific environments used for each step, using the text “env:name” in parenthesis. Unless a specific environment is mentioned, scripts were run in the main Cparapsilosis_popgen_env.

### Obtention of genomes, annotations and databases

We used the reference genome of *C. parapsilosis* strain CDC317, obtained from the Candida Genome Database (CGD)^118^, from http://www.candidagenome.org/download/sequence/C_parapsilosis_CDC317/archive/C_parapsilos is_CDC317_version_s01-m06-r03_chromosomes.fasta.gz. Also, we used the corresponding gff annotations from http://www.candidagenome.org/download/gff/C_parapsilosis_CDC317/archive/C_parapsilosis_CDC 317_version_s01-m06-r03_features.gff. We obtained gene annotations from the CGD tabular file from http://www.candidagenome.org/download/chromosomal_feature_files/C_parapsilosis_CDC317/arc hive/C_parapsilosis_CDC317_version_s01-m06-r03_chromosomal_feature.tab.gz. Furthermore, to get a table with the genomic coordinates with simple repeats (e.g. satellites or low complexity regions) we used CPfun.get_simple_repeats_file. This function first runs RepeatMasker on the reference genome with arguments ‘-poly -html -gff -noint’ (env:perSVade_env_RepeatMasker_env), and then uses custom python code (CPfun.generate_df_repeat_masker_file) to convert the output into a tabular format that is compatible with perSVade’s modules.

To perform quality control of the genomic datasets we also used reference genomes for *Candidozyma auris*, *C. orthopsilosis*, *C. metapsilosis* and *C. albicans*. For *C. auris*, we used the GCA_002759435.2 assembly, and added the mitochondrial chromosome from ASM716870v1. For *C. orthopsilosis*, we used https://ftp.ncbi.nlm.nih.gov/genomes/all/GCF/000/315/875/GCF_000315875.1_ASM31587v1/GCF_000315875.1_ASM31587v1_genomic.fna.gz assembly, and then added the mtDNA chromosome from https://www.ncbi.nlm.nih.gov/nuccore/NC_006972. For *C. metapsilosis*, we obtained the assembly from ^119^. For *C.albicans*, we use haplotype A from http://www.candidagenome.org/download/sequence/C_albicans_SC5314/Assembly22/archive/C_albicans_SC5314_version_A22-s07-m01-r110_chromosomes.fasta.gz.

We retrieved gene pathway annotations from Gene Ontology (GO)^120^ and Reactome^121^. First, we obtained all GO annotations for the CDC317 reference genes (version s01-m06-r03 from CGD), available at http://www.candidagenome.org/download/go/archive/gene_association.20231003.cgd.gz. Second, to process GO annotations we downloaded the GO .obo file from https://purl.obolibrary.org/obo/go/go-basic.obo (latest version by 27 June 2024). Third, we obtained Reactome pathway annotations from https://reactome.org/download/current/ReactomePathways.txt, and their relationships from https://reactome.org/download/current/ReactomePathwaysRelation.txt (latest version by 28 June 2024).

To enable various Kraken runs on uncharacterized sequencing data we downloaded the standard database of reference genomes from RefSeq (latest version by 7 March 2021)^122^, hereafter referred to as KRAKEN2_DB_COMPLETE. Also, to perform a Kraken analysis only on fungi we used the fungal part of the B-GUT database^123^, called Broad_fungal_microbiome_db.

### Obtention of SRA datasets

To contextualize our sequenced isolates within the currently-described diversity of *C. parapsilosis* we used the NCBI SRA run selector website on 6 November 2023 to find available sequencing datasets for the species. Specifically, we looked for all runs matching the species taxon ID (5480), with the filters source=DNA, type=genome, layout=paired, strategy=genome, file type=fastq. Then, we removed a few BioProjects corresponding to *in vitro*-evolved strains (”PRJEB57083”, “PRJEB45347”, “PRJEB45642”, “PRJEB60076”, “PRJEB56880”) or genetically engineered strains (“PRJNA866533”). We kept runs that were i) pure isolates (with LibrarySource=”GENOMIC”), ii) coming from clinical or environmental sources (see CPfun.sra_run_is_clinical_or_enviromental_WGS) and iii) sequenced with Illumina (discarding a few “BGISEQ” and “DNBSEQ” samples). Finally, we discarded the run “ERR9707015” because we could not process it due to possible data truncation. This resulted in 214 runs from patients or human-related environments,hereafter referred to as “SRA runs context”.

Conversely, to root the phylogenetic trees with *C. orthopsilosis* outgroups (see below) we used the same NCBI SRA run selector strategy on 10 November 2023, but focusing on *C. orthopsilosis* datasets. Of these, we kept runs from bioprojects “PRJNA520893”, “PRJNA431439”, “PRJNA322245” and “PRJNA767198”, coming from previous studies sequencing this species^119,124–126^. Next, we kept only seven runs corresponding to highly homozygous strains (# heterozygous variants / kb < 1), as reported in the table Data S4 of ^126^, hereafter referred to as “SRA runs orthopsilosis”. The python code used for this selection is included in CPfun.get_homozygous_Corthopsilosis_srrs.

For all of these runs, we used CPfun.download_srr_with_prefetch to obtain pre-fetched SRR read files (with the tool prefetch of the sra-tools suite). Then, to get the raw fastq files we ran PerSVade’s script get_trimmed_reads_for_srr.py on these pre-fetched files with argument --stop_after_fastqdump (env:perSVade_env).

### Sequencing data mapping and quality control

Quality control and mapping was performed for all sequencing datasets analyzed here using the same pipeline. For this, we initially ran CPfun.get_df_reads_info with min_coverage=50 to process the reads and calculate various relevant statistics. First, this function uses downsample_reads.py to randomly sample 100,000 reads. Second, it runs the read depooling pipeline described in ^14,127^ (see **Supplementary Methods**) to infer the percentage of these 100,000 reads that map to the reference genomes of various *Candida* species (*C. parapsilosis*, *C. auris*, *C. orthopsilosis*, *C. metapsilosis* and *C. albicans*). Third, it calculates the number of total read pairs with wc. Fourth, it calculates the mean read length across the 100,000 read set with CPfun.get_read_length_from_fastqgz (based on perSVade_env). Fourth, it infers the expected depth of coverage as implemented in CPfun.get_chr_to_len, using the number of read pairs, the read length and the reference genome length, so that *expected coverage = (read length * 2 * number pairs) / genome length*.

Fifth, to ensure consistent coverage across samples, CPfun.get_df_reads_info runs the script downsample_reads.py to a number of reads that should generate 50x coverage, according to the formula *reads to downsample = (50 * genome length) / (read length * 2)*. Sixth, to get the trimmed 50x reads and an associated FastQC report^128^ it uses perSVade’s module trim_reads_and_QC (env:perSVade_env). Seventh, to get mapped trimmed reads with different aligners (bwa mem^129^, bowtie2^130^ and hisat2^131^), it runs perSVade’s module map_reads with arguments ‘--min_chromosome_len 5000 --aligner <bwa_mem, bowtie2_local, hisat2_no_spliced> --skip_marking_duplicates’ (env:perSVade_env). Finally, the function integrates all these calculations (number pairs, read length, expected coverage, percent of reads mapping to each *Candida* species) into a single table, useful for further quality control.

To allow for quality control based on the FastQC reports generated by perSVade we ran the function CPfun.run_multiqc_several_trimmed_reads. This arranges the FastQC reports of the trimmed reads in a folder structure and file naming that is compatible with multiqc, and then runs multiqc with arguments ‘--dirs-depth 1 -d -f -s --interactive’ to obtain quality control statistics.

To enable further filtering, we calculated the average depth of coverage per window for all sequencing datasets, different aligners (bwa mem, bowtie2 and hisat2) and various window sizes (200 kb, 100 kb, 50 kb, 20 kb and 1 kb), using CPfun.generate_df_coverage_per_window. This runs get_mappability_file_whole_genome.py on the reference genome (env:perSVade_env) to calculate the mappability per position (using perSVade’s function generate_genome_mappability_file). Next, for large windows (>10kb) it restricts the analysis to gDNA chromosomes, using CPfun.get_genome_excluding_chromosomes and CPfun.get_genomic_df_excluding_chromosomes (based on custom python code). Then, for each sample it runs perSVade’s module call_CNVs with arguments ‘--mitochondrial_chromosome no_mitochondria --skip_CNV_calling -p 2 --window_size_CNVcalling <WINDOW size> --min_chromosome_len 5000 --skip_coverage_correction --mappability_file <MAPPABILITY file> --average_cov_measure median --repeats_file

<REPEATS>’ (env:perSVade_env). This reports, for each window, the median coverage, the fraction of simple repeats, the GC content, the fraction of N bases, the percent of covered positions and the median mappability. Finally, CPfun.generate_df_coverage_per_window uses CPfun.get_cov_df_per_window_one_run to integrate all the coverage calculations into a single table, including the coverage relative to the median of the gDNA windows, which was further used to keep only high-quality datasets for the analysis (see below).

### Small variant calling and annotation

We called small variants (SNPs and small INDELs) using perSVade’s module call_small_variants with arguments ‘-r <*C. parapsilosis* reference> --min_chromosome_len 5000 --repeats_file skip -p 2 --callers HaplotypeCaller,bcftools,freebayes -c 12 --min_AF 0.25 --window_freebayes_bp 200000’ (env:perSVade_env). With these settings, this module calls small variants with three algorithms (GATK HaplotypeCaller^104^, bcftools call^132^ and freebayes^133^) and filters them with ‘best-practices’ parameters for each algorithm^53^. With ‘-c 12’ we keep variants that are in positions with coverage >= 12x. Finally, we define as ‘perSVade filtered variants’ those included in the output file ‘variants_atLeast2PASS_ploidy2.vcf’, which pass the filters for at least two algorithms, and have >= 25% of the reads supporting them (due to ‘--min_AF 0.25’). Note that we applied this exact pipeline also for the *C. orthopsilosis* samples, leading to small variants relative to *C. parapsilosis*, which enabled using them as outgroups (see section ‘Strain tree reconstruction’ below).

To get the integrated filtered small variants, for each aligner we ran CPfun.get_filtered_small_vars_df, using cov_windows_size=1000. This function calculates the median median coverage across 1kb gDNA windows for each sample, based on the per-window coverage calculations performed above. Then, it uses CPfun.get_df_small_vars_filt_one_run in parallel to load the perSVade filtered variants (defined above). The latter function filters out variants overlapping simple repeats using bedtools intersect (env:perSVade_env), and calculates the depth of coverage of variant positions relative to the median gDNA coverage of the sample (called ‘DP_rel_to_gDNA’). Furthermore, CPfun.get_filtered_small_vars_df, uses custom code and bedmap (env:perSVade_env) to transfer various coverage-related calculations of the 1kb windows overlapping the variant (median mappability, median coverage relative to gDNA and percent covered) to the filtered variants datasets. Next, this function adds to the dataset the number of small variants in a given position across all samples. Finally, it uses the output of get_mappability_file_whole_genome.py (described above) to calculate the mappability of each position with a variant. These steps generated all of the filtered high-confidence variants used in this study, together with various variant statistics used in further analyses.

To integrate all raw small variants into a single table and generate their annotations (i.e. impacts on nearby genes) we used fun.integrate_and_annotate_small_vars. For the variants called with each aligner, this function first runs perSVade’s module integrate_several_samples with arguments ‘--min_chromosome_len 100000 -p 2’ (env:perSVade_env), resulting in a single table with all raw variants across samples, called ‘integrated_small_variants.tab’. Next, it runs perSVade’s module annotate_small_vars on these integrated variants with arguments ‘--mitochondrial_code 4 --gDNA_code 12 --min_chromosome_len 100000’ (env:perSVade_env), generating the annotations in ‘small_vars_annot/annotated_variants_corrrectedGene.tab’. Note that we only ran this on a subset of 365 high-quality *C. parapsilosis* datasets (189 sequenced here, 176 from SRA), used in all analyses (except tree reconstruction), as described below.

### Whole genome sequencing datasets filtering

#### Calculation of quality control metrics for all datasets

To ensure high-quality datasets we analyzed several quality parameters. Initially, to be able to discard samples with unexpected SNP density patterns we ran CPfun.generate_variant_per_window_densities_df with aligner=”bwa_mem” and window_size=1000. This function calculates various variant density metrics for 1 kb windows per sample, for different filtered variants (INDELs or SNPs) called with bwa mem, and various zygosities (homozygous, heterozygous and unknown, which corresponds to GT=”.” in the vcf). Specifically, it records the i) set of variants, ii) number of variants, iii) number of variants per kb, iv) mean DP_rel_to_gDNA across variants, v) set of variant allele frequencies (VAF, i.e. the fraction of reads supporting the variant) and v) mean VAF across variants. Also, it adds coverage related metrics calculated previously with CPfun.generate_df_coverage_per_window (see above), including the median coverage, percent genome covered, fraction of repeats, GC content, median mappability, fraction of N bases and median coverage relative to gDNA.

Next, to integrate all these quality metrics into a per-sample dataset we ran CPfun.get_df_reads_info_with_multiqc_and_coverage_data with aligner_stats=”bwa_mem”, window_size_SNP_densities=1000. First, to enable filtering based on multiQC flags this function records relevant quality control flags including per_sequence_gc_content, per_base_n_content and adapter_content (manually curated). For sample ERR263539 we manually set per_sequence_gc_content=”fail”, since it showed an unexpected GC content distribution. Second, it records the expected coverage and % of reads mapping to *C. parapsilosis* during depooling, calculated with CPfun.get_df_reads_info as described above.

Third, to integrate the SNP density patterns into specific metrics for filtering, CPfun.get_df_reads_info_with_multiqc_and_coverage_data processes the 1kb SNP density windows explained above. Initially, it keeps windows with fraction of repeats <= 0.05, fraction of N bases <= 0.05 and median mappability >= 0.25. Then, it runs CPfun.get_r_df_reads_info_with_added_info on each sample to obtain relevant metrics, described next. For coverage-based filtering it records the mean percentage covered for gDNA windows (pct_cov_gDNA). To flag samples with a “smiley pattern”, where the distance to the telomere is negatively correlated to coverage^14^, it calculates the the spearman correlation (scipy.stats.spearmanr) between the distance to the telomere and the median coverage relative to gDNA median (r_distTelomere_vs_cov_gDNA). To identify strains with a noisy coverage pattern it calculates the standard deviation of the differences in median coverage (relative to gDNA median) between adjacent windows (std_rel_cov_diff_windows_gDNA). Finally, to flag possible strain mixes it calculates i) the number of heterozygous SNPs / kb in gDNA windows (hetero_SNPs_per_kb_gDNA) and ii) the median VAF of heterozygous SNPs in non-aneuploid gDNA regions, with median coverage relative to the gDNA median between 0.8 and 1.2 (gDNA_hetero_AF_median).

#### Obtention of high confidence datasets

To keep only high-confidence samples we ran CPfun.run_QC_and_filtering_samples with min_coverage=50, min_pct_reads_para=95 and min_pct_covered=90. This discards runs that have bad FastQC flags (per_sequence_gc_content = “fail”, per_base_n_content different than “pass” and/or adapter_content different than “pass”), including two *C. orthopsilosis* SRA runs, 11 *C. parapsilosis* SRA datasets and four runs sequenced here. To clarify low quality sources we used CPfun.run_kraken_subset_runs, which runs Kraken with a standard (KRAKEN2_DB_COMPLETE) or fungal (Broad_fungal_microbiome_db) database (env:meTAline), followed by Krona’s ktImportTaxonomy with arguments ‘ktImportTaxonomy -t 5 -m 3’ (env:meTAline) to visualize the taxonomy assignments. This showed that the runs sequenced here with bad FastQC flags have a mix of *C. parapsilosis* and other taxa (*Pseudomonas*, *Achromobacter xylosoxidans, C. albicans* or *Bacteria*), supporting our FastQC-based filtering strategy.

Next, CPfun.run_QC_and_filtering_samples filters out *C. parapsilosis* runs with “bad coverage”, having an expected coverage < 50, pct_reads_Cparapsilosis < 95 and/or pct_cov_gDNA < 90, including nine datasets from SRA and 18 runs sequenced here. Also, this function runs CPfun.run_kraken_subset_runs on the “bad coverage” runs sequenced here to identify the source of the unexpected low coverage. Most (15/18) of the datasets contained only *C. parapsilosis* with insufficient coverage, but we identified 3/18 with contamination or co-infection by *Meyerozyma guilliermondii* and *C. albicans*, which shows the importance of considering the read depooling statistics for quality control. Furthermore, our QC function records the datasets that have a “smiley effect” (r_distTelomere_vs_cov_gDNA < −0.1, 48 SRA runs), which is necessary for further analyses of structural variants, which are biased by this effect^14^. In addition, to avoid the effects of excessively noisy coverage patterns it removes 26 *C. parapsilosis* runs (8 sequenced here) with an excessive dispersion in relative coverage measures across windows (std_rel_cov_diff_windows_gDNA > 0.35).

To discard runs with strain mixes CPfun.run_QC_and_filtering_samples flags 14 *C. parapsilosis* runs sequenced here that have hetero_SNPs_per_kb_gDNA >= 0.1 or gDNA_hetero_AF_median < 0.4 or gDNA_hetero_AF_median > 0.6. Also, it uses CPfun.get_df_snps_specific_samples and CPfun.plot_distribution_AFs_specific_samples to generate plots that we inspected to identify, among these 14 runs, four than may be triploids and 10 that are more likely strain mixes, based on VAF distributions. Our QC function discards the mixes, resulting in a final set of 195 samples sequenced here (including four possible triploids and two re-sequenced strains, L21-060 and L21-067), 176 *C. parapsilosis* runs from SRA and 5 *C. orthopsilosis* runs from SRA. These are the so-called high-quality datasets used for further analyses (371 are *C. parapsilosis*). More specifically, for strain tree reconstruction we considered all of them except the possible triploids (367 *C. parapsilosis* isolates in total). Also, for all other analyses and **Supplementary Table 1** we kept only 189 strains sequenced here that were neither duplicates (keeping the latest run for L21-060 and L21-067) nor potential triploids, and the 176 *C. parapsilosis* runs from SRA (365 *C. parapsilosis* isolates in total), called ‘final high-confidence set’.

Note that across all these filtering steps we set quality control thresholds (on expected coverage, pct_reads_Cparapsilosis, pct_cov_gDNA, r_distTelomere_vs_cov_gDNA, std_rel_cov_diff_windows_gDNA, hetero_SNPs_per_kb_gDNA and gDNA_hetero_AF_median) based on manual exploration of the distribution of these metrics, using CPfun.plot_distribution_AFs_specific_samples at various steps of CPfun.run_QC_and_filtering_samples.

#### Generation of metadata table

To integrate all the relevant genomic, clinical and phenotypic metadata for the 371 high-confidence *C. parapsilosis* isolates analyzed here we ran CPfun.get_df_metadata_all_samples. For the high-confidence isolates sequenced here, this function first runs CPfun.get_df_metadata_final to integrate, into a single table, the measured phenotypes (described above) and clinical metadata about the infected patients (hospital, city, patient age, isolation source, year of collection). This function harmonizes the data and translates relevant terms from Spanish to English. CPfun.get_df_metadata_all_samples also parses the NCBI SRA run selector results to add to the table the country, year and city of isolation for the SRA datasets analyzed. Also, for those SRA runs that were analyzed in a recent phylogenetic study of *C. parapsilosis*^42^, the function records the assigned clade (**Fig. 1**).

### Analysis of recombination

To find evidence of genome-wide recombination and/or parasexual diversification we tried to identify pairs of isolates, out of the final high-confidence set of 365 runs, with genomic windows of low pairwise SNP divergence as compared the average pairwise divergence, as done before^10,15^. For this, we ran CPfun.get_stats_pairwise_SNP_distances_per_window to calculate such pairwise divergent estimates, explained next. First, for each aligner (bwa mem, hisat2 and bowtie2) and various window sizes (200 kb, 100 kb, 50 kb, 20 kb and 1 kb) we inferred variant densities per window in each run with CPfun.generate_variant_per_window_densities_df (explained above). To tailor the window sizes to different genome types, we considered 20-200 kb sizes for gDNA windows and 1 kb size for mtDNA windows.

Then, to calculate the per window pairwise variant distances between each ‘query’ sample and all the others (‘targets’), for each ‘query’ sample, aligner and window size we ran the script get_genetic_distances_one_sample.py. This loads the per-window variant densities datasets, and then keeps high-confidence windows with i) fraction of repeats <= 0.05, ii) fraction of N bases <= 0.05, iii) median mappability >= 0.9 and iv) median coverage relative to gDNA >= 0.25 in all samples. Furthermore, for each variant type (SNP, INDEL), zygosity (homozygous, heterozygous and unknown) and ‘target’ sample it runs CPfun.get_df_stats_distances_one_pair_samples_and_types_vars to infer the number of shared and different variants for each high-confidence window.

Next, to integrate these per window comparisons into per isolate pair metrics we used CPfun.get_dfs_stats_pairwise_SNP_distances_per_window_one_run, resulting in a dataset with various metrics for each aligner, window size, zygosity (only homozygous or heterozygous), type of variant (SNP or INDEL), ‘query’ and ‘target’ runs. Most importantly, it calculates, for each pairwise isolate comparison, the i) number of variants per kb across the genome (whole_genome_diff_vars_per_kb) and ii) fraction of the genomic windows with variants per kb below a threshold (0.001, 0.005, 0.01, 0.05, 0.1, 0.15) (fraction_diff_vars_per_kb_below_<THRESHOLD>).

Finally, for finding strain pairs with evidence of recombination we plotted all of the values of whole_genome_diff_vars_per_kb vs fraction_diff_vars_per_kb_below_<THRESHOLD> for different aligners, window sizes, zygosities and thresholds, as shown in **Extended Data Fig. 3** for a few examples based on bwa mem-called SNPs. However, we could not find any pair of strains with an unexpectedly high fraction_diff_vars_per_kb_below_<THRESHOLD> given their whole_genome_diff_vars_per_kb (**Extended Data Fig. 3**), suggesting that this species underwent mostly asexual propagation.

### Structural variant calling and annotation

To call, integrate and annotate the breakpoint-inferred Structural Variants (SVs) and coverage-derived Copy Number Variants (CNVs) (**Extended Data Fig. 1**) we used CPfun.get_cmds_perSVade_SVs_and_CNVs_calls (with SV_parms_file=”auto”) on 317 runs that have no “smiley pattern”. Due to SV_parms_file=”auto”, this function initially identifies optimal SV filtering parameters using various steps. To identify a subset of representative isolates for parameter benchmarking it first runs perSVade’s module get_stats_optimization with arguments ‘-min_chromosome_len 100000 --overlap_coverage 25 --overlap_insert_size 25 --overlap_insert_size_sd 25 --overlap_read_len 25 --overlap_coverage_mtDNA 100’ (env:perSVade_env). This suggested 15 runs that have distinct gDNA coverage, read length and insert size, within > 25% range of each other.

Then, to identify adequate parameters, CPfun.get_cmds_perSVade_SVs_and_CNVs_calls runs CPfun.run_perSVade_parameter_optimization_several_samples on these 15 representative datasets. This runs perSVade optimize_parameters with arguments ‘--min_chromosome_len 5000 --regions_SVsimulations random --simulation_ploidies diploid_hetero --nvars 50 --nsimulations 2 --range_filtering_benchmark theoretically_meaningful --keep_simulation_files’ (env:perSVade_env), leading to a set of optimized parameters (based on SV simulations) for each run. Then, it runs perSVade analyze_SV_parameters with argument --min_chromosome_len 5000 (env:perSVade_env) to check the performance of these optimized parameters on the simulated SVs of all representative runs. After inspecting the graphical outputs of this module we conclude that all 15 optimized parameters work equally well on all representative runs, so we selected the most conservative ones (from sample L21-190) as the single set of optimal SV filtering parameters to be used in all samples.

Next, to get all called and filtered SVs and CNVs for all datasets, CPfun.get_cmds_perSVade_SVs_and_CNVs_calls runs perSVade run_several_modules with arguments ‘call_CNVs,call_SVs,integrate_SV_CNV_calls -p 2 --min_chromosome_len 100000 --cnv_calling_algs HMMcopy,AneuFinder --window_size_CNVcalling 300 --skip_coverage_correction --mappability_file <MAPPABILITY file> --average_cov_measure median --max_fraction_N_bases 0.1 --max_fraction_repeats 0.1 --min_median_mappability 0.75 --SVcalling_parameters <OPTIMAL L21-190 parameters>’ (env:perSVade_env). Most of these arguments (except --min_chromosome_len and --SVcalling_parameters) are set to configure the call_CNVs module, so that CNVs are called on windows of 300 bp, in a diploid mode, using HMMcopy (1.32.0)^134^ and AneuFinder (1.18.0)^135^ callers, skipping coverage corrections and ignoring windows with fraction N bases > 0.1, fraction simple repeats > 0.1 and/or median mappability < 0.75. Also, note that ‘run_several_modules’ is run differently depending on the aligner used. Specifically, for bwa_mem we run all modules, but with hisat2 and bowtie2-mapped reads we only call CNVs (skipping the call_SVs module and the --SVcalling_parameters argument), since these aligners do not produce outputs that are directly compatible with perSVade’s SV calling capacities.

Furthermore, to integrate all SVs and CNVs generated by integrate_SV_CNV_calls in each dataset, CPfun.get_cmds_perSVade_SVs_and_CNVs_calls runs perSVade integrate_several_samples with arguments ‘--min_chromosome_len 100000 -p 2 --tol_bp 50 --pct_overlap 75 --CNV_overlap_only_based_on_pct’ (env:perSVade_env), resulting in a single table with all SVs and CNVs across samples, called ‘integrated_SVs_CNVs.tab’. Due to the arguments ‘--tol_bp 50 --pct_overlap 75 --CNV_overlap_only_based_on_pct’, SVs of a given type are the considered same if they reciprocally overlap by >=75% and have their breakends <=50 bp apart. Also, CNVs of a given type are considered the same if if they reciprocally overlap by >=75%. Finally, CPfun.get_cmds_perSVade_SVs_and_CNVs_calls runs perSVade’s module annotate_SVs on these integrated variants with arguments ‘--mitochondrial_code 4 --gDNA_code 12 --min_chromosome_len 100000’ (env:perSVade_env), generating the variant annotations (i.e. impacts on genes) in ‘SV_CNV_annotation/annotated_variants_corrrectedGene.tab’.

### Strain tree reconstruction

#### General strain tree reconstruction pipeline

To reconstruct a SNP-based phylogenetic tree for a given set of isolates we used the tree_from_SNPs pipeline (based on ^9^). For different tree reconstruction steps, described below, we used CPfun.get_cmds_tree_reconstruction_one_configuration, which acts as a wrapper to i) prepare the inputs for and run the script get_tree.py from the tree_from_SNPs pipeline and ii) generate various quality control calculations on the tree. It uses ete3 (env:tree_from_SNPs_env) for all tree manipulation steps.

This function takes various relevant arguments, described next. First, “runIDs_para” refers to the list of *C. parapsilosis* strains to consider. Second, “aligner” indicates the set of variants to be used, obtained based on the reads mapped with the selected aligner (bwa mem, hisat2 or bowtie2). Third, min_coverage_pos, passed to –-min_coverage_pos in get_tree.py, indicates the minimum coverage across all isolates for a position to be considered for tree reconstruction. Fourth, “min_support_trees” indicates the minimum branch support for a node to be considered “properly resolved”. Fifth, “outgroup” refers to a run ID to be set as an outgroup for rooting the tree, passed to --outgroup in get_tree.py. It can be a single *C. orthopsilosis* run, a single *C. parapsilosis* run or a comma-separated list of *C. parapsilosis* runs reflecting an outgroup clade; the latter being set as outgroup if found in the unrooted tree(s), or else the most similar found clade would be set as outgroup. Sixth, “mode” reflects the mode of tree reconstruction, passed to --mode in get_tree.py. In ‘diploid’ mode, this script first constructs 100 resampled trees based on all homozygous SNPs and a random selection of heterozygous SNPs^136^, and then integrates them into a consensus, which is a common approach for diploid *Candida* species^9,42^. Conversely, in ‘diploid_homozygous’ mode the script generates a single tree based on homozygous SNPs while discarding positions that have some heterozygous SNP in any isolate, as done previously for haploid *Candida* species^9^.

The function CPfun.get_cmds_tree_reconstruction_one_configuration first prepares the files that are necessary for running get_tree.py for each run in runIDs_para and the chosen *C. orthopsilosis* run (if set as outgroup). These include the aligned reads and perSVade-filtered small variants (variants_atLeast2PASS_ploidy2.vcf) obtained with the indicated aligner. Then, it runs get_tree.py with arguments ‘--mode <MODE< --min_coverage_pos <MIN_COVERAGE_POS< --mode_fasta_gen python --outgroup_resampled_trees <OUTGROUP< --batch_mode --repeat_tree_integration’ (env:tree_from_SNPs_env). Next, it uses CPfun.add_tree_cmds_to_list to handle the running of the resampled trees in “diploid” mode in a computationally efficient way. Furthermore, it obtains the outgroup-rooted tree for further calculations, which may be i) the one generated by get_tree.py if mode=”diploid” or ii) the one generated by CPfun.get_outgroup_rooted_tree based on the unrooted output of get_tree.py if mode=”diploid_homozygous”. Finally, it manipulates this raw rooted tree with CPfun.get_correct_tree, resulting in a high-confidence “corrected” rooted tree where nodes with a branch support below min_support_trees are collapsed into their parents.

Beyond the tree itself, CPfun.get_cmds_tree_reconstruction_one_configuration performs various quality control calculations. First, it calculates the number of parsimony informative sites used to generate the trees (nsites_parsimony), parsing the iqtree output generated by get_tree.py with CPfun.get_n_parsimony_informative_sites_from_iqtreefile, for either i) all 100 resampled trees if mode=”diploid” or ii) the only one generated tree if mode=”diploid_homozygous”. Second, if the outgroup is a list of strains (clade), it calculates the fraction of trees (out of 100 resampled for mode=”diploid” or 1 for mode=”diploid_homozygous”) having i) said clade as outgroup (fraction_trees_expected_outgroup) or ii) an alternative, most frequent clade as outgroup (fraction_trees_expected_most_frequent_outgroup). Third, from the high-confidence “corrected” tree this function calculates the percentage of nodes that are collapsed due to low branch support (i.e. <min_support_trees) (pct_collapsed_nodes). Fourth, from the raw rooted tree it calculates the percentage of nodes that would be collapsed due to low branch support (i.e. <min_support_trees) and have a long branch length (i.e. >= 1% of the maximum pairwise distance between any pair of tree nodes) (pct_collapsed_long_nodes). Fifth, from the high-confidence “corrected” tree this function calculates the percentage of nodes that constitute “diverse politomies”, i.e. having some of the children nodes with >=3 leaves (pct_politomies_diverse). Sixth, from the corrected tree this function records the list of *C. parapsilosis* isolates that belong to the *C. parapsilosis* outgroup clade (Cpara_outgroup_leaves), which is particularly useful to identify such an outgroup when the outgroup used for rooting is a *C. orthopsilosis* strain.

We used this CPfun.get_cmds_tree_reconstruction_one_configuration for various analysis steps, referenced below, setting specific arguments for runIDs_para, mode, outgroup, aligner, min_coverage_pos and min_support_trees.

#### Tree reconstruction for all isolates

We reconstructed a SNP-based tree for all isolates with high-confidence sequencing datasets except the possible triploids (see above), including 191 sequenced here (where L21-060 and L21-067 are sequenced twice) and 176 *C. parapsilosis* from SRA (367 in total), running CPfun.generate_SNPs_trees. We used CPfun.get_cmds_tree_reconstruction_one_configuration setting these runs as runIDs_para, min_support_trees=90 and min_coverage_pos=12. To assess the effect of different tree modes (diploid and diploid_homozygous) and aligners (bwa mem, hista2 and bowtie2) we generated six trees using combinations of these parameters, set as arguments mode and aligner in CPfun.get_cmds_tree_reconstruction_one_configuration.

We lacked a clear outgroup clade in *C. parapsilosis* to root the trees, as previous studies reported unrooted trees^42^ or used suboptimal midpoint rooting^9^. To identify the *C. parapsilosis* root we first reconstructed trees including both the *C. parapsilosis* 367 isolates and each of the five high-confidence homozygous *C. orthopsilosis* strains obtained from SRA (described above), which can be used as outgroups. Specifically, we ran CPfun.get_cmds_tree_reconstruction_one_configuration with aligner=”bwa_mem” 10 times, trying all possible combinations of the arguments mode (“diploid” or “diploid_homozygous”) and outgroup (each of the five *C. orthopsilosis* runs, one for each tree); all within the function CPfun.generate_SNPs_trees. Although these trees were poorly resolved (all with pct_collapsed_nodes > 50 and nsites_parsimony < 6,000), they suggest the same *C. parapsilosis* outgroup clade of 146 isolates being the most common for both modes (3/5 for mode=diploid, 4/5 for mode=diploid_homozygous).

Finally, we obtained the six *C. parapsilosis*-only trees for variants called with different aligners and modes using this 146 isolate clade as outgroup, using CPfun.get_cmds_tree_reconstruction_one_configuration on the 367 isolates analyzed here set as runIDs_para. To select the optimally-reconstructed tree out of the six we checked their quality control metrics (nsites_parsimony, fraction_trees_expected_outgroup, pct_collapsed_nodes, pct_collapsed_long_nodes, pct_politomies_diverse) and also compared them in a pairwise manner (**Extended Data Fig. 2**). Specifically, to get the differences between two high confidence “corrected” trees we used CPfun.get_pct_different_nodes_two_trees. Given two trees A and B, this function calculates the percentage of nodes in A that have a set of leaves not found in B, and *vice versa*, yielding two pairwise percentual distances: *d_AB_*and *d_BA_*. Then, we calculated the ‘minimum percent of equal nodes’ as 100 - max(*d_AB_*, *d_BA_*). Based on the analysis of these metrics we used mode=diploid and aligner=bwa_mem as the optimal parameters for tree reconstruction of all isolates (see **Supplementary Results**).

#### Clade inference

To define clades and sub-clades for the SNP-based tree (**Fig. 1b**) we transferred the clade definitions (1, 2, 3, 4 or 5) from Bergin et. al.^42^ to the generated tree. Initially, we used CPfun.get_trees_df_row_with_Bergin2022_clade_info to record the clade for isolates falling within previously defined clades. However, this left out 69 isolates, which belong to new sub-clades (**Fig. 1b**). To assign new sub-clade IDs we first ordered the tree to match the clade order 1-5 from Bergin et. al. with CPfun.get_tree_ordered_clades, and then assigned the new sub-clades (1.1, 4.1, 4.2 and 4.3) with CPfun.get_leaf_to_clade_Bergin2022_added_clades. We used Cfun.get_cladeID_info_file_Bergin2022 to transform clade definitions into a tabular file called ‘cladeID_Bergin2022_info.tab’ (used for GWAS as described in **Supplementary Methods**).

### Definition of binary phenotypes for genotype-phenotype analyses

To define binary phenotypes we ran CPfun.get_df_metadata_binary_phenotypes_curated on the metadata table generated by CPfun.get_df_metadata_all_samples (described above). This processes phenotypic and clinical metadata for the 189 high-confidence isolates analyzed here, resulting in 61 binary variables, including 1) resistance towards the antifungals amphotericin B, 5-flucytosine, fluconazole, itraconazole, voriconazole, posaconazole, isavuconazole, caspofungin, micafungin and anidulafungin, 2) tolerance towards ethanol, hypochlorite, H_2_O_2_, chlorhexidine, zwittergent 3-14 and surfanio, 3) fluconazole heteroresistance, 4) higher fluconazole resistance in solid vs liquid media, 5) fluconazole tolerance at 48h, 6) invasiveness in agar, 7) pseudohyphae formation, 8) biofilm formation, 9) various adherence behaviors in a microfluidics device, 10) patient age group (infant, young or old) and 11) various body isolation sources.

To enable comparisons between isolates with and without each of these traits, for a given trait each isolate may have a value of 1 (i.e. it has the trait), 0 (i.e. it lacks the trait) or -1 (i.e. the trait was either not measured or unclear). For instance, isolates with intermediate drug susceptibility, unmeasured biofilm formation or low agar invasiveness would have a −1 value. Below, we detail how we processed various datasets to get these traits in a binary format, while the corresponding code is in CPfun.get_df_metadata_binary_phenotypes_curated.

#### Antifungal resistance and disinfectant tolerance

To transform antifungal susceptibility Minimum Inhibitory Concentration (MIC) data to binary phenotypes, we set custom MIC thresholds to define resistant / susceptible strains, based on established clinical resistance breakpoints (EUCAST v10)^137^ and the observed MIC distributions (**Extended Data Fig. 4a**). To enable relevant comparisons these thresholds did not always match the clinical breakpoints. Specifically, for amphotericin B, anidulafungin and micafungin our dataset did not contain any isolates with resistance over the breakpoint, so that we set a lower threshold (one concentration below the breakpoint) to capture the difference between high or low susceptibility (**Extended Data Fig. 4a**). This means that for these drugs the “resistance” phenotype is below the clinical breakpoint of that species, but still meaningful with respect to the observed MIC distribution of the species.

Conversely, for four azoles (fluconazole, voriconazole, itraconazole and posaconazole) we had isolates with resistance over the clinical breakpoint, but the MIC distribution for each drug made us set the thresholds differently. For fluconazole and voriconazole we considered that the clinical breakpoints were adequate to distinguish clearly binary resistant (1) / susceptible (0) strains for various reasons. First, for both drugs the breakpoints consider a range of intermediate susceptibility (resistance=-1), as there is one breakpoint for susceptible and one for resistant strains. Second, these breakpoints separate a bimodal MIC distribution into two clear halves, indicating two distinct phenotypes (**Extended Data Fig. 4a**). However, for itraconazole and posaconazole the clinical breakpoint falls around the median of a unimodal MIC distribution, which means that adopting it would mean having no isolates with intermediate susceptibility (resistance=-1). As the distribution is unimodal, we consider that the lack of intermediate susceptibility category does not reflect the true biological nature of itraconazole and posaconazole susceptibility phenotypes. Also, given the absence of technical replicates in our measurements, not having this category may lead to some noise in the susceptible / resistant classification. Thus, we assign “intermediate susceptibility” to those isolates with the lowest MIC that is considered clinically resistant (**Extended Data Fig. 4a**).

Finally, for 5-flucytosine, isavuconazole and caspofungin there are no clinical breakpoints^137^, so we set the thresholds based on the MIC distribution (**Extended Data Fig. 4a**), leaving always a range of intermediate susceptibility to ensure clearly binary susceptible / resistant assignments. Similarly, we to define disinfectant tolerance binary traits (towards ethanol, hypochlorite, H_2_O_2_, chlorhexidine, zwittergent 3-14 and surfanio) we set custom thresholds based on the distribution of the percent of viability in each of them (**Extended Data Fig. 4e**). All of the binarization of drug resistance and disinfectant tolerance phenotypes was performed with CPfun.get_dicothomic_resistance.

Although it may have been tempting to perform genotype-phenotype analyses directly on the continuous phenotypes (MIC, percent of viability), we considered antifungal susceptibility and disinfectant tolerance as binary due to several reasons. First, as we discarded isolates with intermediate phenotypes (see **Extended Data Fig. 4a,e**) we made sure that the comparisons made reflect clear phenotypic changes, less affected by small experimental errors. Second, treating all variables as binary allowed us to apply a single pipeline to all analyzed phenotypes (for GWAS and ML analyses), ensuring consistency and comparability of different traits. Third, it simplified the reconstruction of ancestral states used in GWAS / ML analyses. Fourth, this approach allowed us to predict clear phenotype transitions, useful for ML analyses. Admittedly, our approach has the limitation that we ignore isolates with intermediate phenotypes in our analyses. However, for the biological mechanisms learnt here this should not be an issue, as it is preferable to ensure that compared samples have truly different phenotypes. However, this limitation should be considered if the ML classifiers are deployed in a clinical setting.

#### E-test phenotypes

To extract relevant binary phenotypes from the E-test (agar solid media) measurements (described above) we explored the distributions of i) fluconazole MIC in liquid media (FLC_MIC), ii) fluconazole MIC in E-test at 24h and 48h (MIC_Etest_24h, MIC_Etest_48h) and iii) whether there are colonies within the E-test inhibition halo. Based on these, we identified three phenotypes that are variable across strains: fluconazole heteroresistance (heteroresistance), higher fluconazole resistance in solid vs liquid media (resistance_higher_solid) and fluconazole tolerance at 48h (tolerance). First, among isolates that showed mild fluconazole resistance (FLC_MIC between 8-32), we identified some forming colonies in the inhibition halo (heteroresistance=1) and others that didn’t (heteroresistance=0) (**Extended Data Fig. 4b**). In this case, heteroresistance=-1 (not available) refers to isolates for which the E-test was not performed or with a FLC_MIC outside the 8-32 range.

Second, among isolates with fluconazole resistance in liquid (FLC_MIC between 16-64), we found some with much higher resistance in solid media (MIC_Etest_24h=512, resistance_higher_solid=1) and others with comparable resistance in solid media (MIC_Etest_24h between 6-48, resistance_higher_solid=0) (**Extended Data Fig. 4c**). Thus, resistance_higher_solid=-1 refers to isolates for which either the E-test was not performed, with a FLC_MIC outside the 16-64 range or with MIC_Etest_24h different from 512 and outside the 6-48 range. Third, among isolates with mild fluconazole resistance in solid media at 24h (MIC_Etest_24h between 6-16), we could find some with very high MIC at 48h (MIC_Etest_48h=512) (tolerance=1) and others with an intermediate one (tolerance=0) (**Extended Data Fig. 4d**). Thus, tolerance=-1 refers to isolates for which either the E-test was not performed or with a MIC_Etest_24h outside the 6-16 range. All of the code to set these binary traits can be found in CPfun.get_df_metadata_binary_phenotypes_curated.

#### Virulence traits

To convert the virulence measurements of invasiveness, morphology, biofilm formation and microfluidics behavior (described above) into binary traits (**Fig. 2**) we performed a few simple transformations. On the one hand, we one-hot-encoded the values of morphology and microfluidics behavior to generate binary variables. Specifically, we defined the binary variable ‘pseudohyphae formation’ to encode morphology information (which may be ‘yeast’ or ‘pseudohyphae’). Also, we created three new variables based on the one-hot-encoding of the microfluidics behavior values ‘adherent’, ‘big clumps’ and ‘little clumps’ (described above), called ‘microfluidics adherent’, ‘microfluidics big clumps’ and ‘microfluidics little clumps’. On the other hand, for agar invasiveness and biofilm formation, which may be ‘high’, ‘low’ or ‘no’, we considered isolates with ‘high’ as 1, ‘no’ as 0, and ‘low’ as −1 (not considered), ensuring the considered isolates have clearly distinct phenotypes. Also, note that for all these virulence binary phenotypes we set values of −1 for missing values. All related code can be found in CPfun.get_df_metadata_binary_phenotypes_curated.

#### Features of the infected patients

Beyond these experimentally determined phenotypes, we considered some clinical features of the source patients as relevant ‘traits’. This enabled us to study whether specific genetic variants predispose isolates towards infecting certain patient age groups and/or body parts. On the one hand, based on the patient age distribution (**Extended Data Fig. 5a**) we defined the phenotypes ‘age infant’, ‘age young’ and ‘age old’ for each isolate. For this, we first filtered out two isolates infecting patients of (registered) 124 years, resulting from missing age information which was registered as birth date of 01/01/1900. Then, we binned isolates according to the age group of the infected patient: ‘age_infant’ refers to <10 years, ‘age_young’ refers to 20-50 years and ‘age_old’ refers to 80-110 years. Also, for each of these ‘traits’ we define as ‘negative controls’ (e.g. age_infant=0) the isolates infecting patients of the most common age range (60-75 years), which would enable relevant comparisons. For each of these age group traits, isolates with value=-1 would be those infecting patients with an age outside the range of the group and outside the negative control range.

On the other hand, we defined each of the isolation sources (i.e. body compartment) as binary traits, where isolates from a given source have value=1 and those of other sources have value=0 (**Fig. 2**). Also, to gain power we grouped them in clinically-meaningful ways (**Extended Data Fig. 5b**) and defined the following ‘traits’ (each is a comparison). First, ‘source sterile’ refers to sterile compartments (blood culture, whole blood, vascular tissue, heart valve, cerebrospinal fluid, ascitic fluid, bile fluid) (value=1) vs all other compartments (value=0). Second, ‘source sterile vs environmental’ refers to sterile compartments (value=1) vs the samples from the hospital environment (value=0), where all other sources have value=-1. Third, ‘source blood’ includes blood-related compartments (blood culture, whole blood, vascular tissue) (value=1) vs all other compartments (value=0).

Fourth, ‘source blood vs environmental’ refers to blood-related compartments (value=1) vs the samples from the hospital environment (value=0), where all other sources have value=-1. Fifth, ‘source blood vs other sterile’ refers to blood-related compartments (value=1) vs other non-blood-related sterile compartments (cerebrospinal fluid, ascitic fluid, bile fluid) (value=0), where all other sources have value=-1. Sixth, ‘source skin’ refers to skin-related sources (skin exudate, wound exudate, wound, skin surface) (value=1) vs all other compartments (value=0) except the blood-related ones (blood culture, whole blood, vascular tissue) (value=-1) and samples from the hospital environment (value=-1). Seventh, ‘source respiratory’ refers to compartments within the respiratory system (bronchial aspirate, tracheal aspirate, lung biopsy, oropharyngeal sputum, sputum, oropharyngeal exudate, upper respiratory exudate, bronchoalveolar lavage, respiratory disease) (value=1) vs all other compartments (value=0) except the blood-related ones (blood culture, whole blood, vascular tissue) (value=-1).

All relevant code can be found in CPfun.get_df_metadata_binary_phenotypes_curated.

### Convergence Genome Wide Association Study (GWAS)

We attempted to run a convergence GWAS for the 61 binary phenotypes, based on the variants generated with each aligner (bwa mem, hisat2, bowtie2). For this, we developed the ancestral_GWAS toolkit, which builds on our previous work (see Methods of ^9^) but improves usability and computational performance. Most importantly, we ran run_GWAS.py from ancestral_GWAS, a pipeline using Ancestral State Reconstruction (ASR) to find variant transitions associated with phenotypic transitions in their reconstructed evolutionary histories. In addition, to take into account that different variants may change the phenotype by altering the same feature (a protein domain, gene or pathway), run_GWAS.py tests the association between groups of collapsed variants and the phenotype, in addition to the single variant tests (**Fig. 4**). Specifically, each of these groups refers to a combination of type_vars (combinations of small variants, SVs and/or CNVs), type_mutations (all, non synonymous, non synonymous non truncating or truncating), type_collapsing (at the level of variant effect (e.g. *ERG11* variant G458S), domain, genes, GO, Reactome, MetaCyc or none for single-variant groups) and group ID (e.g. gene CPAR2_600430).

To benchmark parameters we ran run_GWAS.py with varying settings. First, to address biases generated by clonal redundancy (**Fig. 2**) we ran the analysis on all isolates (sample_set = all_samples), on a subset of them where we only keep three isolates for each clade that is monophyletic for a given phenotype value (e.g. **Extended Data Fig. 6a**) (sample_set = representative_samples), and on an equivalent subset with one isolate per monophyletic clade (sample_set = balanced_samples). Second, to evaluate node filtering parameters we changed i) the minimum support that a node should have to be considered for GWAS, called ‘min_support’ (10, 50, 70 or 90) and ii) the ASR method used, called ‘ASR_method’ (MPPA, DOWNPASS or MPPA,DOWNPASS consensus). Third, we varied the aligner (bwa mem, bowtie2 or hisat2) used to generate the variants.

For each of the tested variant groups, our pipeline calculates various metrics used for filtering. First, ‘epsilon’ (*ε*) refers to the convergence level between genotype and phenotype calculated as proposed before^9,76^, which is 1.0 if all variant transitions in the tested group are associated to a phenotype transition and *vice versa*. Second, *n_G&P_* indicates the number of nodes with genotype and phenotype transitions. Third, *X*^2^ refers to the chi-square statistic of a contingency table associating genotype and phenotype transitions across nodes. Fourth, *p(X*^2^) and *p(n_G&P_)* reflect the empirical probability of observing a *X*^2^ or *n_G&P_*, respectively, equal or higher than the observed one across 10,000 GWAS based on resampled phenotypes. Fifth, we also include the bonferroni and FDR-corrected *p(X*^2^), *p(n_G&P_)* values, and the maxT-corrected *p(X*^2^*)_maxT_*, *p(ε)_maxT_*ones as defined in ^9^. Additionally, we applied these corrections across either i) all tested groups (for a given combination of aligner, ASR method, min support, and sample set) (conservative ‘correction scope’) vs ii) all groups with the same variant grouping strategy (certain values for ‘type_vars’, ‘type_mutations’ and ‘type_collapsing’) (relaxed ‘correction scope’).

To analyze phenotypes with sufficient variation across traits we focused on those with >=3 phenotype transitions by all aligners, in either all or representative samples, according to the GWAS run with ASR_method = “DOWNPASS” and min_support = 70. We calculated these numbers of transitions by parsing GWAS results with CPfun.get_integrated_GWAS_df_stats_pheno_transitions, and then integrating them with CPfun.get_min_ntransitons_across_aligners_one_phenotype. Also we discarded the trait ‘whole blood source’, which was highly redundant with ‘source blood’. Finally, we analyzed GWAS results for 27 phenotypes, including those related to i) antifungal resistance (towards 5-flucytosine, amphotericin B, anidulafungin, caspofungin, micafungin, fluconazole, voriconazole, posaconazole), ii) disinfectant tolerance (for chlorhexidine, hypochlorite, zwittergent 3-14, iii) virulence (agar invasiveness), iv) patient age (old, young and infant) and iv) isolation sources (abscess, bronchial aspirate, sputum, skin exudate, blood culture, bronchoalveolar lavage, ascitic fluid, urine, blood, blood vs other sterile, skin, sterile) (**Fig. 4b**). Also, to focus on stringent genotype-phenotype associations we used the results of the gwas_method = ‘synchronous’, referring to the GWAS in which we measure convergence between phenotype and genotype transitions (see **Supplementary Methods**).

Furthermore, to identify high and low confidence hit filtering strategies (see **Supplementary Results**) we obtained various quality control metrics (set of expected genes found, fraction of phenotype transitions that can be attributed to some protein-altering hit and number of ‘false positive’ variant hits) for different filter combinations with CPfun.bechmark_GWAS_parameters. This initially defines the filter combinations with CPfun.get_gwas_filters_df, and outputs the quality control metrics for each filter set and phenotype with CPfun.get_GWAS_benchmarking_df_one_result. Also, we used CPfun.get_df_GWAS_parameter_benchmark_filtered to further process this output for visual parameter exploration (see next step), adding fields about the types of p values used by different filtering sets (**Extended Data Fig. 8a**). Finally, we visually assessed different filterings using CPfun.explore_GWAS_parameters_plots, and generated the final so-called ‘raw’ list of filtered hits (with high or low confidence) using CPfun.get_filtered_GWAS_dfs.

To generate the raw list of high-confidence hits we used min_support = 70, aligner = hisat2, ASR method = MPPA,DOWNPASS, sample_set = balanced_samples, min *ε* = 0.3, min *n_G&P_* = 2, (only) bonferroni-corrected *p(n_G&P_)* < 0.05 and a relaxed correction scope. Conversely, to generate the raw list of low-confidence hits we used min_support = 10, aligner = hisat2, ASR method = MPPA, sample_set = all samples, min *ε* = 0.2, min *n_G&P_* = 2, (only) uncorrected *p(n_G&P_)* < 0.05. Then, we removed redundancy across these raw hits with CPfun.get_NR_GWAS_tables, resulting in a list of hits where each variant is only mapped to a single tested group, as found in **Supplementary Table 4,5**. Also, we used plot_tree_variants_multiple_phenos.py from the ancestral_GWAS toolkit to visualize the variants related to these hits along the strain tree. For additional details on GWAS analyses see **Supplementary Methods**.

### Building Machine Learning (ML) classifiers

To build the ML classifiers for each phenotype we also used various scripts within the ancestral_GWAS toolkit. In brief, we evaluated the performance of various i) predictive features (i.e. variant groups such as those used for GWAS) selected with forward feature selection, ii) ML models (e.g. gradient boosting or decision trees) and iii) model hyperparameters (e.g. tree depth). Specifically, for different combinations of ‘test parameters’ (described below) that we wanted to evaluate, we built five models (m1-m5) using 80% of the data for training / evaluation using Cross-Validation (CV), and 20% for testing, each m1-m5 corresponding to a different split. Note that the ‘evaluation’ set is often also referred to as the ‘validation’ set in ML studies, but we call it ‘evaluation’ to differentiate it from the ‘external validation’ set (see below). Also, for simplicity, across the output files and scripts of our ML pipeline we refer to the ‘training / evaluation set’ as the ‘CV set’.

For each phenotype, this process of training and evaluation involved benchmarking thousands of parameter combinations, called ‘train parameters’, related to the selection of features, models and hyperparameters. Specifically, we identified a set of optimal ‘train parameters’ with high predictive performance (ROC curve AUC) and feature consistency across multiple train / evaluation splits (described below). Finally, we assessed the performance and m1-to-m5 feature consistency of such optimal ‘train parameters’ on the testing data, leading to one such testing for each combination of ‘test parameters’ (**Fig. 5 a,b**).

To only build ML classifiers for phenotypes with sufficient phenotypic variation we analyzed 13 traits with at least 5 phenotypic transitions by all aligners. Specifically, for each phenotype and aligner we calculated the number of transitions as the maximum found across GWAS runs on all or representative sample sets with ASR_method = “DOWNPASS” and min_support = 70 (described above), as calculated in CPfun.get_integrated_GWAS_df_stats_pheno_transitions. These phenotypes include i) several body sources (skin exudate, urine, bronchial aspirate, blood, sterile, and blood vs other sterile), ii) antifungal resistance (towards micafungin, fluconazole and voriconazole), iii) the patient age groups (infant, young, old) and iv) invasiveness in agar.

A key aspect of our ML pipeline is related to the ‘train’ and ‘test’ parameter distinction, explained next. On the one hand, we wanted to evaluate how various so-called ‘test parameters’ influence i) model performance on the testing data and ii) potential for overfitting the training data. These include ‘aligner’, ‘training_type_split’, ‘sample_set’, ‘AUC_diff_threshold’, ‘consistency_measure’ and ‘gene_set’, described next. First, ‘aligner’ refers to the mapper algorithm from which the variants come (bwa mem, bowtie2, hisat2). Second, ‘training_type_split’ indicates the way how we perform the train / evaluation splitting, which can be random or phylogenetically-balanced (see **Extended Data Fig. 6b**). Third, ‘sample_set’ refers to the sample set to use for training and evaluation (see **Extended Data Fig. 6a**), which may be ‘all’ or ‘representative’ samples, as generated for the GWAS. Fourth, ‘AUC_diff_threshold’ configures the balance between performance (ROC AUC) and feature consistency in the selection of ‘optimal train parameters’ (see **Supplementary Methods**), and we considered the values 0.05, 0.1, 0.15 and 0.2. Fifth, ‘consistency_measure’ refers to how we measure feature consistency, which can be based on the variants, strains or genes related to these features, or ‘none’ to disregard feature consistency as a metric to optimize during training and evaluation.

Sixth, ‘gene_set’ indicates the subset of genes considered for classifier building, related to the initial set of features on which to perform training and evaluation. Specifically, we defined such features by either i) performing a convergence GWAS to pinpoint groups associated to the phenotype with the ‘synchronous’ method in the training / evaluation set (gene_set = ‘all’), or ii) only considering features related to the gene set of interest (other values for gene_set). We tried various combinations of expected genes, described below. All of these ‘test parameters’ configure the training / evaluation process, which explains why we cannot use it to select them.

On the other hand, the ‘train parameters’ are those that can be treated as true model hyperparameters, i.e. they can be optimized through the train and evaluation process. These include ‘min_support’, ‘ASR_method’, ‘type_filtering’, ‘prioritize_expected_genes’, ‘feature_selection_AUC_threshold’, ‘model’, ‘type_phenotype’, ‘transition_definition’, ‘transition_SNPkb_threshold’, ‘transition_used_prediction’, ‘transition_ML_threshold’ and ‘class_weight’, described next. First, ‘min_support’ reflects the minimum branch support used in the initial GWAS filtering when gene_set = ‘all’, and we tried the values 50, 70 and 90. Second, ‘ASR_method’ indicates the method used for ASR in this same GWAS, and we tried MPPA, DOWNPASS and MPPA,DOWNPASS (defined above). Third, ‘type_filtering’ indicates how the such initial GWAS hits were filtered, which may be i) ‘p < 0.05’, where filtered features should have a raw *p_FISHER_* < 0.05 or ii) ‘all’, so that filtered features are those with any convergence with the phenotype (*n_G&P_* >= 1). Fourth, ‘prioritize_expected_genes’ (yes / no) reflects whether we prioritize models that have some of the expected genes related to the selected features.

Fifth, ‘feature_selection_AUC_threshold’ indicates the ‘tol’ parameter used in the sklearn.feature_selection.SequentialFeatureSelector package, reflecting the minimum ROC AUC increment required for a new feature to be included during forward feature selection (it may be 0.05, 0.1 or 0.15). Sixth, ‘model’ refers to the ML modelling approach used for building the classifier, which we chose based on previous ML studies in bacteria and fungi^46,48,138^. It may be AdaBoost, random forest (RF), gradient boosting (GBoost), K-nearest neighbors (KNN), Bernoulli naive bayes (BernoulliNB), Gaussian naive bayes (GaussianNB), logistic regression (LR), support vector classifier (SVC) or decision tree (tree). Also, note that for KNN, GBoost, SVC and tree models we tried various combinations of them varying the configuration hyperparameters. Seventh, ‘type_phenotype’ refers to the way the phenotype was predicted. This may be ‘phenotype’ (the phenotype is directly modelled based on the features) or ‘phenotype_transition’ (we model transitions in phenotype between several pairs of isolates, and then use the prediction of transition / no-transition to infer the actual phenotype). For ‘phenotype_transition’, we i) predict transition / no-transition instances (according to the parameters ‘transition_definition’ and ‘transition_SNPkb_threshold’, see below), ii) keep certain types of predicted transitions / no transitions (according to the parameter ‘transition_used_prediction’), and iii) infer the phenotype probability based on the transition probabilities of a given sample (according to the parameter ‘transition_ML_threshold’).

Eighth, ‘transition_definition’ indicates how we define pairs of compared isolates to define transition / no-transition instances when type_phenotype = ‘phenotype_transition’. This parameter may be ‘two_close_balanced’ (compare each isolate with the two closest ones with and without the same phenotype) and ‘each_pheno_clade’ (identify clades monophyletic for the phenotype and compare each isolate with one representative of each clade). Ninth, ‘transition_SNPkb_threshold’ refers to the threshold on the genetic distance to select pairs of isolates, when type_phenotype = ‘phenotype_transition’. We tried the values 0.1, 0.25, 1.0. Tenth, ‘transition_used_prediction’ indicates the types of predicted transitions used for the phenotype inference. It may be ‘all’, ‘only_transitions’ (only use actually predicted transitions) and ‘only_no_transitions’ (only using predicted no-transitions). Eleventh, ‘transition_ML_threshold’ refers to the ML probability threshold used to convert transition probabilities to predicted transitions (0,1 values), and we tried the values 0.25, 0.5, 0.75. Twelfth, ‘class_weight’ (‘balanced’ or ‘none’) indicates whether a balanced class weighing is applied, which we only considered applying for phenotypes and sample sets with >= 80% class imbalance.

For each phenotype, we evaluated 576 or 1872 ‘test parameters’ (depending on the phenotype) and 5,940 - 216,864 ‘train parameters’ (depending on the test parameters and phenotypes). We used CPfun.run_ML_predictors_phenotypes to build all training, evaluation and testing datasets. For each of these test parameters we selected the optimal combination of training parameters, and we evaluated the resulting model performance (Receiver operating characteristic (ROC) curve AUC on testing data) and model feature consistency (overlap of features chosen across the m1-m5 models). Then, we focused on models with i) no clear sign of overfitting or underfitting (ROC AUC >= 0.7 on the training data and a small difference (up to 0.2) between the training / testing ROC AUCs), ii) acceptable performance on testing data (ROC AUC >= 0.7), and iii) high feature consistency (consistency >= 0.4). For each of the phenotypes with such acceptable models, we analyzed the performance of the ‘top testing parameters’, i.e. those with an AUC >= max AUC - 0.05 and consistency within top 10, using two testing strategies involving the datasets sequenced here. First, the ‘test’ strategy involved obtaining aggregate performance estimates (ROC AUC, precision, recall, specificity and false positive rate (fpr)) across m1-m5 splits, yielding one estimate for each set of testing parameters. Second, the ‘cross-val’ strategy involved getting similar performance estimates for each m1-m5 split and set of testing parameters. Note that the precision, recall, specificity and fpr were calculated using optimal probability thresholds that yielded the maximum recall with a fpr<=0.3. Also, we used plot_tree_variants_multiple_phenos.py of the ancestral_GWAS toolkit to visualize ML features along the strains tree. For additional details see **Supplementary Methods**.

### Validation of ML classifiers on external data

To perform the external validation of our ML classifiers we downloaded all available WGS paired-end run metadata for *C. parapsilosis* on 5 December 2024, using the NCBI SRA run selector strategy described in the section ‘Obtention of SRA datasets’ above. Then, we used CPfun.get_df_sra_runs_validation to filter the runs and keep those that come from BioProjects including isolate collections with many genomes sequenced and antifungal susceptibility phenotypes measured^33,37,38,77–79^. Also, we ran CPfun.get_df_phenotypes_validation to generate a table with the MIC values for the different antifungals reported in these studies. This function mostly relies on custom python and pandas code to parse different tables and get MIC values for each run, and also performs various quality control steps. First, it discards “SRR25837524” because the MICs were wrongly reported for this run. Second, it runs CPfun.fetch_sra_runs_metadata to get SRAStudy metadata for our runs through epost and efetch (env:Candida_mine_env), which was necessary to discard runs from study “SRP233635” (*C. auris* isolates) that were included in one of our BioProjects. Finally, we obtained 114 runs with antifungal susceptibility phenotypes measured in previous studies hereafter referred to as “SRA runs validation’.

Among these datasets, we found 99 extra ones as compared to the previously assembled ‘SRA runs context’ (see ‘Obtention of SRA datasets’), which is reasonable given that almost one year passed between the assembly of both strain collections leading to new strains being added to SRA. For these extra runs we downloaded the raw reads with the same pipeline as for the others (see ‘Obtention of SRA datasets’ above). Similarly, we used CPfun.get_df_reads_info to trim the reads, map them, and calculate various statistics for quality control (see ‘Sequencing data mapping and quality control’). Furthermore, we performed the small variant calling as mentioned in ‘Small variant calling and annotation’. Next, we filtered these 99 extra datasets to keep the high-confidence ones, with adequate coverage and quality (44/99 in total), following the criteria described in ‘Whole genome sequencing datasets filtering’, as implemented in CPfun.get_df_reads_info_analysis_extra_validation. Note that none of these isolates had a VAF distribution suggesting strain mixing, following the criteria implemented within CPfun.run_QC_and_filtering_samples (defined above). All these strains are in **Supplementary Table 1.**

Then, for 409 strains (365 initial + 44 extra) we obtained integrated small variants, CNVs and SVs as described in the sections ‘Small variant calling and annotation’ and ‘Structural variant calling and annotation’ above, as implemented in CPfun.get_integrated_variant_calls_all_datasets_with_validation. Also, for each of the phenotypes where we could find sufficient data (voriconazole and fluconazole resistance), we prepared various files including all strains with binary 0/1 phenotype values (sequenced here or from the SRA), used for subsequent validation of ML models. These files include i) the binarized phenotypes based on our resistance thresholds (**Extended Data Fig. 4a**), ii) the strains tree built with CPfun.generate_tree_one_phenotype_and_aligner (see **Supplementary Methods**), and iii) the convergence GWAS input files necessary for ML runs, obtained with CPfun.get_cmds_convergence_GWAS (see **Supplementary Methods**) run with skip_GWAS_run = True. Notably, the latter generated integrated variants (called ‘variants_df_all.tab’), variant annotations (‘variant_annotations_df_all.tab’), a corrected tree (‘corrected_rooted_tree.nw’) and the phenotypes file (‘phenotypes_all.tab’). All these files were generated as implemented in CPfun.prepare_files_for_geno_pheno_validation. Note that all isolates phenotyped in other studies fall within the clades described here (**Fig. 1**, **Extended Data Fig. 12**), validating our 189 strains as representative to build generalizable ML models.

Next, to obtain the ‘validation’ ML metrics (**Fig. 5d**) we further prepared various files. For each of the models obtained with the ‘top testing parameters’ (defined above) for fluconazole and voriconazole resistance, we first defined a set of training samples that may be i) all samples sequenced here with phenotypic information if sample_set = ‘all_samples’, or ii) all representative samples for a given phenotype if sample_set = ‘representative_samples’. Additionally, we defined a set of testing samples including the representative ones out of the phenotyped SRA isolates, defined as explained in ‘Convergence Genome Wide Association Study (GWAS)’. Then, we created a merged dataset (called ‘sample_info.tab’) including these training and testing samples and their phenotypes (from ‘phenotypes_all.tab’). Furthermore, we pruned the corresponding ‘corrected_rooted_tree.nw’ to only include such training and testing samples, resulting in ‘treefile_prunned.nw’. In addition, we put into ‘thresholds.txt’ the list of probability thresholds considered for the top model, enabling calculation of threshold-dependent metrics. Additionally, we wrote ‘model_info.tab’ including information about the top model to test; where each row is one combination of testing parameters and test_split_ID, and includes i) the features used, ii) various relevant training parameters (model, class_weight, type_phenotype, transition_definition, transition_SNPkb_threshold, transition_used_prediction, transition_ML_threshold), iii) various relevant testing parameters (aligner andsample_set).

Finally, we ran evaluate_ML_models.py from the ancestral_GWAS toolkit (env:ancestral_GWAS_env) to get the validation metrics, using arguments “--variants_file variants_df_all.tab --variant_annotations_file variant_annotations_df_all.tab --samples_file sample_info.tab --treefile treefile_prunned.nw --models_file model_info.tab --gff annotations.gff --ref_genome genome.fasta --target_geneIDs −TARGET genes− --pathway_annotations_dir pathway_annotations/ --CGD_chromosomal_features CGD_fieatures.tab --interproscan_output interproscan_annotation.out --gDNA_code 12 --mtDNA_code 4 --mtDNA_chrom mito_C_parapsilosis_CDC317 --default_pheno_probability 0.5 --thresholds_file thresholds.txt”. Note that the various arguments (--gff, --ref_genome, --pathway_annotations_dir, --CGD_chromosomal_features, --interproscan_output, --gDNA_code, --mtDNA_code, --mtDNA_chrom) are related to the processing of functional annotations, as further described in **Supplementary Methods**. Also, to --target_geneIDs we passed a comma-separated list of genes mapped to each of the selected features of each top model, used for filtering variants.

Most importantly, this script trains each of the models using the ‘top’ parameters and features specified in ‘model_info.tab’ on the training samples, and calculates performance metrics on the testing samples. For models where type_phenotype = phenotype_transition, the transitions of the testing samples are defined based on comparing each testing sample to the corresponding training samples (depending on transition_definition, transition_SNPkb_threshold, transition_used_prediction, transition_ML_threshold). Note that evaluate_ML_models.py could be used further to directly run our models on newly sequenced datasets. This script already provided the ‘validation’ ROC AUC (**Fig. 5e,f**), and to get the ‘validation’ specificity, recall and precision we used the optimal probability thresholds defined above for the ‘cross-val’ testing strategy (yielding maximum recall with fpr<=0.3). All these ‘validation’ ML metrics were calculated as implemented in CPfun.get_df_ML_results_with_validation_accuracy_metrics.

### Integration of ML and GWAS results

To integrate the results of the analysis of expected genes, high-confidence convergence GWAS hits and ML classifier features, we assessed the 298 genes yielded by these approaches in 15 phenotypes with either GWAS high-confidence hits or built ML classifiers. These included the four expected genes (*ERG11*, *TAC1*, *MRR1* and *NDT80*) manually associated with fluconazole, voriconazole and posaconazole resistance (**Fig. 3**). Also, we considered the 285 genes related to high-confidence GWAS hits across 13 traits (**Fig. 4d, Supplementary Table 4**). Furthermore, we included the 16 genes related to ML features for the four phenotypes in which the classifiers could be built (**Fig. 6a**).

To prioritize relevant candidates across these 298 genes we kept, for each phenotype, a subset of genes based on various sequential inclusion criteria. First, we kept the expected ‘known’ genes for azole resistance phenotypes (*ERG11*, *TAC1*, *MRR1* and *NDT80*) (criterion = ‘known gene’). Second, we included the genes related to ML features (criterion = ‘in ML features’). Third, for azole and amphotericin B resistance we kept genes whose orthologs in other *Candida* species have been associated to equivalent resistance phenotypes (azoles and amphotericin B, respectively) in a recent GWAS study^9^ (criterion = ‘GWAS hit Schikora-Tamarit 2024’). Fourth, we included genes with a functional description in CGD that is related to the phenotype of interest, which we defined based on manual curation as implemented in CPfun.get_phenotype_to_all_possible_genes_for_ML (criterion = ‘functional description’). Fifth, we kept genes with orthologs (in other *Candida* species) having signs of recent genomic selection by SNPs, as reported in a recent study^9^ (criterion = ‘under selection Schikora-Tamarit 2024’). Sixth, for phenotypes with none of the previous types of genes we included the top three genes with highest fraction of phenotypic transitions associated to a variant in the gene, based on the high-confidence GWAS hits, as implemented in CPfun.get_df_gwas_one_pheno_with_per_gene_stats (criterion = ‘top 3 genes’). This pipeline resulted in 50 genes.

We visualized the type of analysis (manual curation, ML, high-confidence GWAS or low-confidence GWAS) that yields each of these genes in a given phenotype (see **Fig. 7**). Also, for each gene in each phenotype we visualized either i) the first inclusion criteria of the sequential list described in the previous paragraph (e.g. ‘known gene’ has priority over ‘in ML features’ or ‘functional description’) or ii) the fraction of phenotypic transitions associated to a variant in the gene by high or low-confidence GWAS hits in case that the gene was not included in the prioritized subset of genes for that phenotype. This allowed us to pinpoint relevant trends across the genotype-phenotype associations. See CPfun.plot_ML_GWAS_overlap_heatmap for additional information.

Also, to identify the most relevant genes for each phenotype we focused on i) genes that were prioritized in each phenotype and ii) for ergosterol-related phenotypes (fluconazole R, voriconazole R, posaconazole R and amphotericin B R), genes prioritized in other ergosterol-related phenotypes with GWAS hits (high or low confidence) in the phenotype of interest (criterion = ‘ergosterol phenotypes’). For each of these 79 gene-phenotype mappings we then defined the relevant variant effects on the gene in different ways, depending on the type of associations that yielded them (‘GWAS low-confidence’, ‘GWAS high-confidence’, ‘ML & GWAS high-confidence’, ‘manual curation only’, ‘ML only’ or ‘ML & GWAS low-confidence’, see **Fig. 7**). First, for ‘GWAS low-confidence’ associations we kept the variant effects from **Supplementary Table 5**, which include all variants with transitions associated to phenotype transitions. Second, and similarly, for ‘GWAS high-confidence’ and ‘ML & GWAS high-confidence’ associations we kept the variant effects from **Supplementary Table 4**. Third, for ‘manual curation only’ associations (i.e. *ERG11*, *TAC1*, *MRR1* and *NDT80* in azoles that have no related GWAS or ML features) we kept nonsynonymous variants with transitions associated to phenotype transitions when using aligner = ‘bwa mem’, ASR_method = ‘MPPA,DOWNPASS’, min_support = 70, sample_set = ‘representative_samples’ (as in **Fig. 3**), inferred with the GWAS redundancy reduction pipeline (see **Supplementary Methods**) as implemented in CPfun.get_df_var_info_NR_certain_variant_groups.

Fourth, for ‘ML only’ and ‘ML & GWAS low-confidence’ associations we kept variants related to the ‘top models’ ML features of the gene of interest as yielded by a certain aligner (prioritizing hisat2, then bowtie2, then bwa mem) and a certain sample_set (prioritizing representative_samples over all_samples). To get the variants of such features we used CPfun.get_df_var_info_NR_certain_variant_groups, and then we kept either i) only those with transitions associated to phenotype transitions or ii) all variants if none of them had such correlated transitions. These final 79 gene-phenotype mappings are found in **Supplementary Table 6**, and the code to generate them is in CPfun.generate_GWAS_and_ML_integrated_table.

## ACKNOWLEDGEMENTS

The authors acknowledge support from the Spanish Ministry of Science and Innovation (grant numbers PID2021-126067NB-I00, PID2023-148686OB-I00 by MICIU/AEI/10.13039/501100011033 and by FEDER, UE, and PLEC2023-010225) cofounded by ERDF “A way of making Europe”, as well as support from the Gordon and Betty Moore Foundation (grant number GBMF9742); the Catalan Research Agency (AGAUR) (grant number 2022 INNOV 00065, 2024 PROD 00175 and 2024 PROD 00043); “La Caixa” foundation (grant number LCF/PR/HR21/00737 and CI23-20260); Fundació La Marató de TV3 (202328-31); AECC (PRYGN234923GABA and 290059); Instituto de Salud Carlos III (CIBERINFEC CB21/13/00061 and CB21/13/00105 - ISCIII-SGEFI/ERDF and DTS25/00141, ESI-2024 PI24CIII/00051); European Commission, Horizon Europe-HORIZON-MSCA-2023-DN-01-01 (grant number 101168618) and European Union’s Horizon 2020 research and innovation programme under the Marie Skłodowska-Curie grant agreement N° 101226544 (grant number 101227078). This work was also partially sponsored by a non-restricitive grant by Gilead Sciences. Also, MAST acknowledges his AI4S fellowship within the “Generación D” initiative by Red.es, Ministerio para la Transformación Digital y de la Función Pública, for talent attraction (C005/24-ED CV1), funded by NextGenerationEU through PRTR. Finally, we want to thank the members of the CAPAR Study Group for their feedback regarding the clinical interpretation of our results.

## AUTHOR CONTRIBUTIONS

OZ and TG conceived the study and obtained funds. MAST performed all computational analyses and prepared all the figures. TG supervised the computational analysis. OZ supervised the experimental work and the clinical information. MAST, TG and OZ wrote a first draft of the manuscript. ELP, AR, CA, LAF and ATC performed experimental work. The members of the CAPAR Study Group provided the clinical isolates and their related clinical information.

## DATA AVAILABILITY

All of the sequencing datasets generated here are available under project code PRJNA1439216 in SRA. Also, **Supplementary Table 1** contains all strains analyzed, with the measured phenotypes.

## CODE AVAILABILITY

All of the code and software environments used to generate the datasets, results, tables and figures presented here are in https://github.com/Gabaldonlab/Cparapsilosis_GenoPheno. Also, the code and software environments to run the tree reconstruction and GWAS / ML pipelines are in https://github.com/Gabaldonlab/tree_from_SNPs and https://github.com/Gabaldonlab/ancestral_GWAS_toolkit, respectively. The latter may be particularly useful beyond this project.

## STUDY GROUP INFORMATION

The CAPAR Study Group was integrated by the following authors: Maite Ruiz Pérez de Pipaón^1,14^, Marta López-Lomba^2^, Teresa Durán Valle^2^, Paloma Merino Amador^3^, Fernando González Romo^3^, Mireia Puig Asensio^4,14^, Carmen Ardanuy^5,15^, María Teresa Martín-Gómez^6^, Daniel Romero^6^, Gregoria Megías^7^, María Ángeles Mantecón Vallejo^7^, Ana Pérez de Ayala^8^, Isabel Sánchez Romero^9^, María Pía Roiz^10,14^, Isabel Lara Plaza^10^, Teresa Nebreda^11^, María Simón Sacristán^12^, Eneritz Velasco-Arnaiz^13^, Julià Gotzens^13^, Manuel Monsonís-Cabedo^13^.

1. Hospital Virgen del Rocío, Sevilla, Spain
2. Servicio de Microbiología. Hospital Universitario de Móstoles, Madrid, Spain.
3. Servicio de Microbiología. Hospital Clínico San Carlos. Instituto de Investigación Sanitaria Hospital Clínico San Carlos (IdISSC), Madrid, Spain
4. Servicio de Microbiología. Hospital Universitari de Bellvitge-IDIBELL-UB, Barcelona, Spain
5. Microbiology Department, Bellvitge University Hospital. Biomedical Research Institute of Bellvitge Institut d’Investigació Biomèdica de Bellvitge, IDIBELL), Barcelona, Spain.
6. Microbiology Department. Vall d’Hebron University Hospital, Barcelona, Spain
7. Complejo Asistencial Universitario de Burgos, Burgos, Spain
8. Hospital Universitario 12 de Octubre, Madrid, Spain
9. Hospital Universitario Puerta De Hierro, Majadahonda, Spain
10. Servicio de Microbiología. Hospital Universitario Marqués de Valdecilla-IDIVAL, Santander, Spain
11. Hospital Clínico Universitario de Valladolid, Valladolid, Spain
12. Hospital Central de la Defensa Gomez Ulla, Madrid, Spain
13. Infectious Diseases and Systemic Inflammatory Response in Pediatrics, Pediatric Infectious Diseases Department, Institut de Recerca Sant Joan de Déu. Hospital Sant Joan de Déu, Barcelona, Spain.
14. CIBER de Enfermedades Infecciosas, Instituto de Salud Carlos III, Madrid, Spain
15. CIBER de Enfermedades Respiratorias, Instituto de Salud Carlos III, Madrid, Spain

## EXTENDED DATA

**Extended Data Fig. 1.**
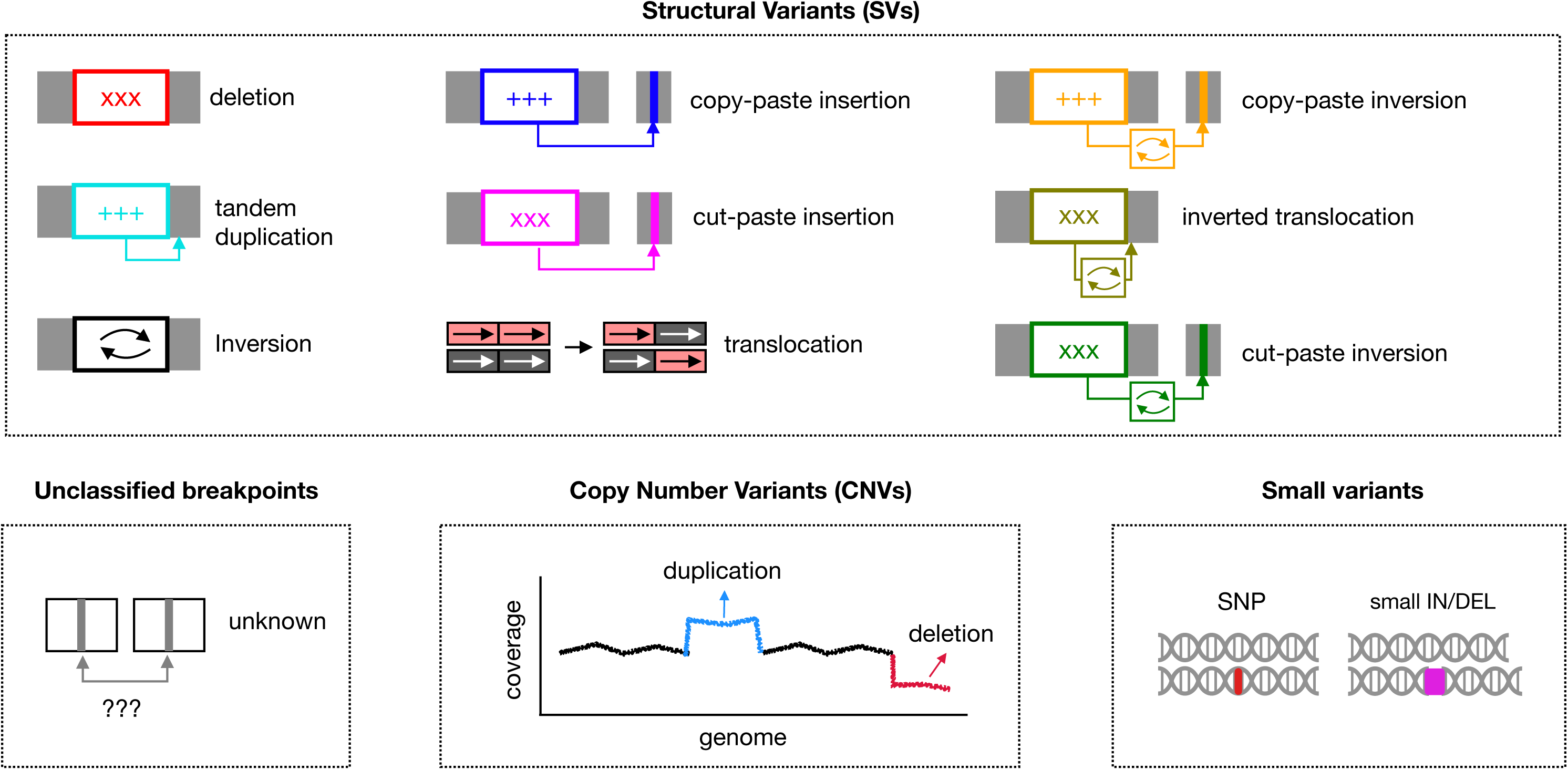
Variant types analyzed in this work. These are the variant types inferred with perSVade^53^. The distinction between SVs and CNVs is technical. While all CNVs are technically a type of SVs, the way how they were inferred is different. SVs are found based on split read, discordantly aligned read pairs and *de novo* assembly evidence. Conversely, CNVs are inferred based on changes in read coverage. This figure is adapted from ^9^.

**Extended Data Fig. 2.**
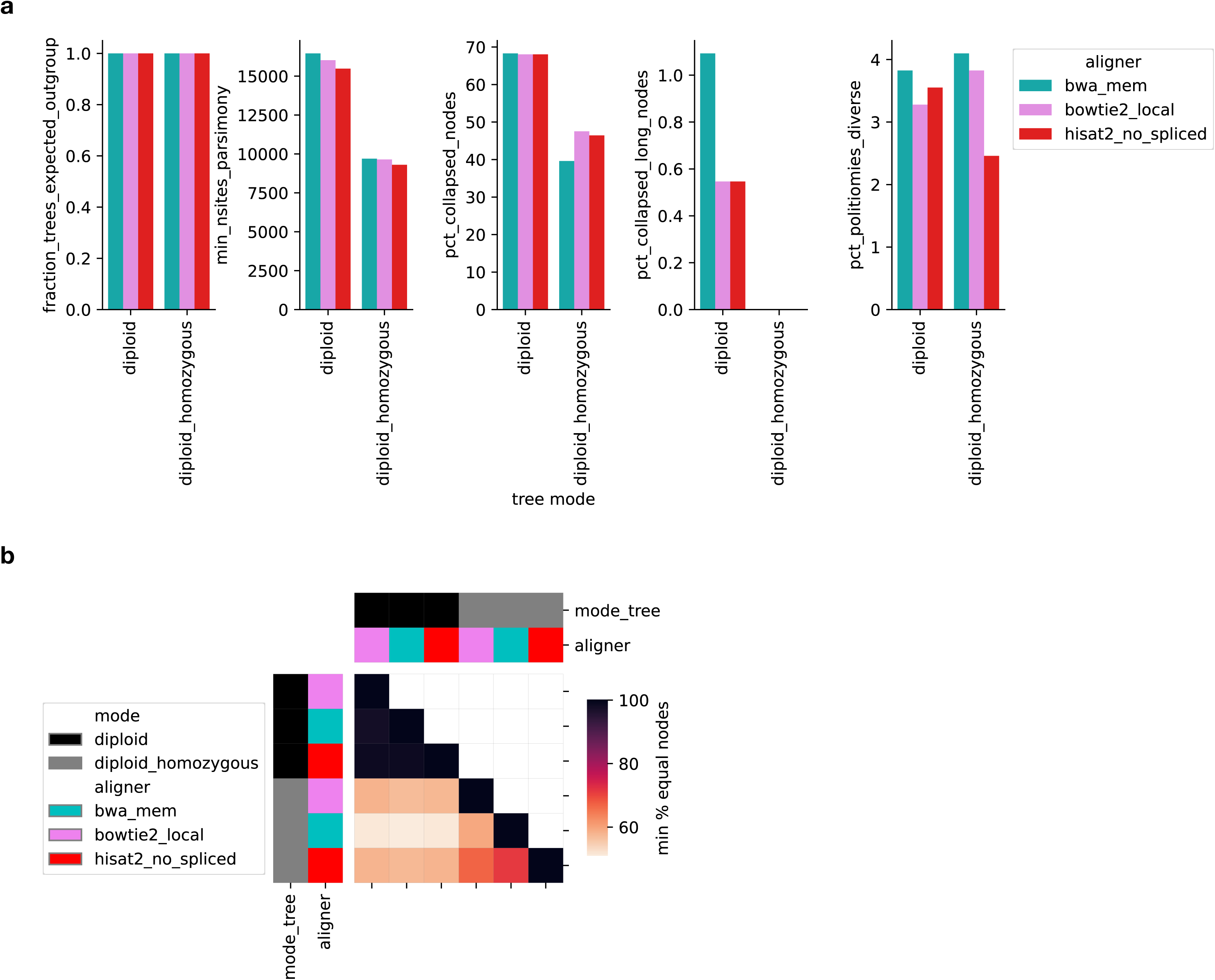
Parametrization of strain trees. (a) Various metrics for *C. parapsilosis*-only trees built using i) SNPs called from reads mapped with different aligners (colors), ii) various modes of handling of heterozygous SNPs (x-axis). For mode = diploid, ‘fraction_trees_expected_outgroup’ reflects the fraction of resampled unrooted trees that include a clade with the expected *C. parapsilosis* outgroup; while for mode = diploid_homozygous it indicates whether the final tree includes such outgroup. For mode = diploid, ‘min_nsites_parsimony’ indicates the minimum number of parsimony-informative sites in the alignment used to build each of the resampled trees; while for mode = diploid_homozygous it indicates the number of parsimony-informative sites used to build the final tree. The value ‘pct_collapsed_nodes’ indicates the percentage of nodes with support < 90, while ‘pct_collapsed_long_nodes’ indicates the percentage of nodes with support < 90 and long branches (see **Methods**). In addition, pct_politomies_diverse indicates the percent of nodes that have support < 90 and have children nodes with support >= 90 and >=3 strains. (b) Heatmap representing, for each pair of A / B trees compared as indicated in rows / columns, 100 - max(*d_AB_*, *d_BA_*), where *d_AB_* is the percentage of nodes in A that have a set of leaves not found in B, and *vice versa* for *d_BA_*.

**Extended Data Fig. 3.**
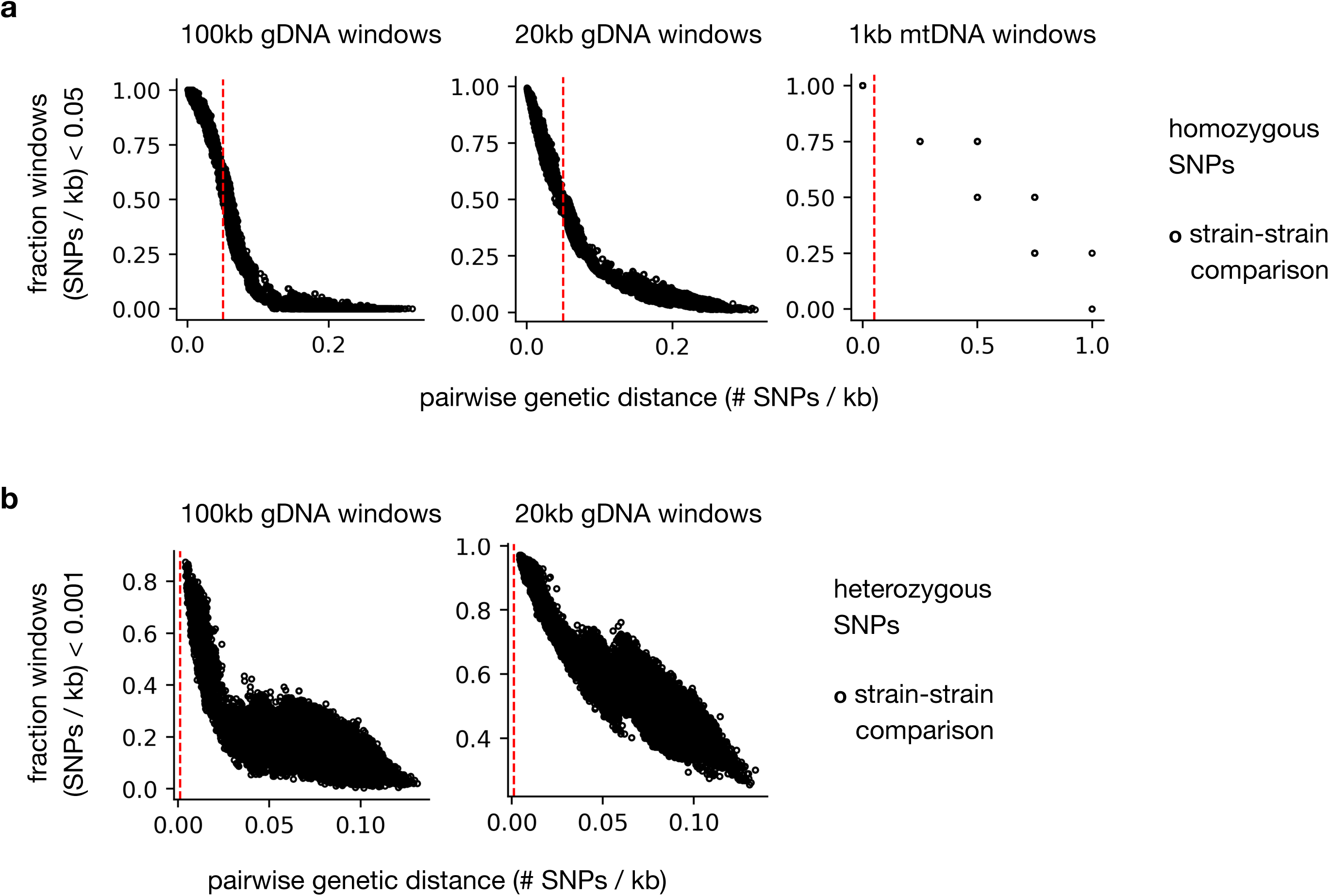
Analysis of genome-wide parasexual diversification. (a) To detect signs of parasexual diversification we calculated, for each pair of our 365 isolates i) the genetic distance (number of different homozygous SNPs / kb) and ii) the fraction of genomic windows (of either 100kb, 20kb or 1kb) with a low genetic distance (SNPs / kb < 0.05). Each point corresponds to one strain-strain comparison. We used windows of 100kb and 20kb for the gDNA regions (left and middle panels), and 1 kb for the mtDNA (right panel). (b) The same as in (a), but for heterozygous SNPs. Note that we do not show data for the mtDNA, where the calling of heterozygous SNPs is not plausible.

**Extended Data Fig. 4.**
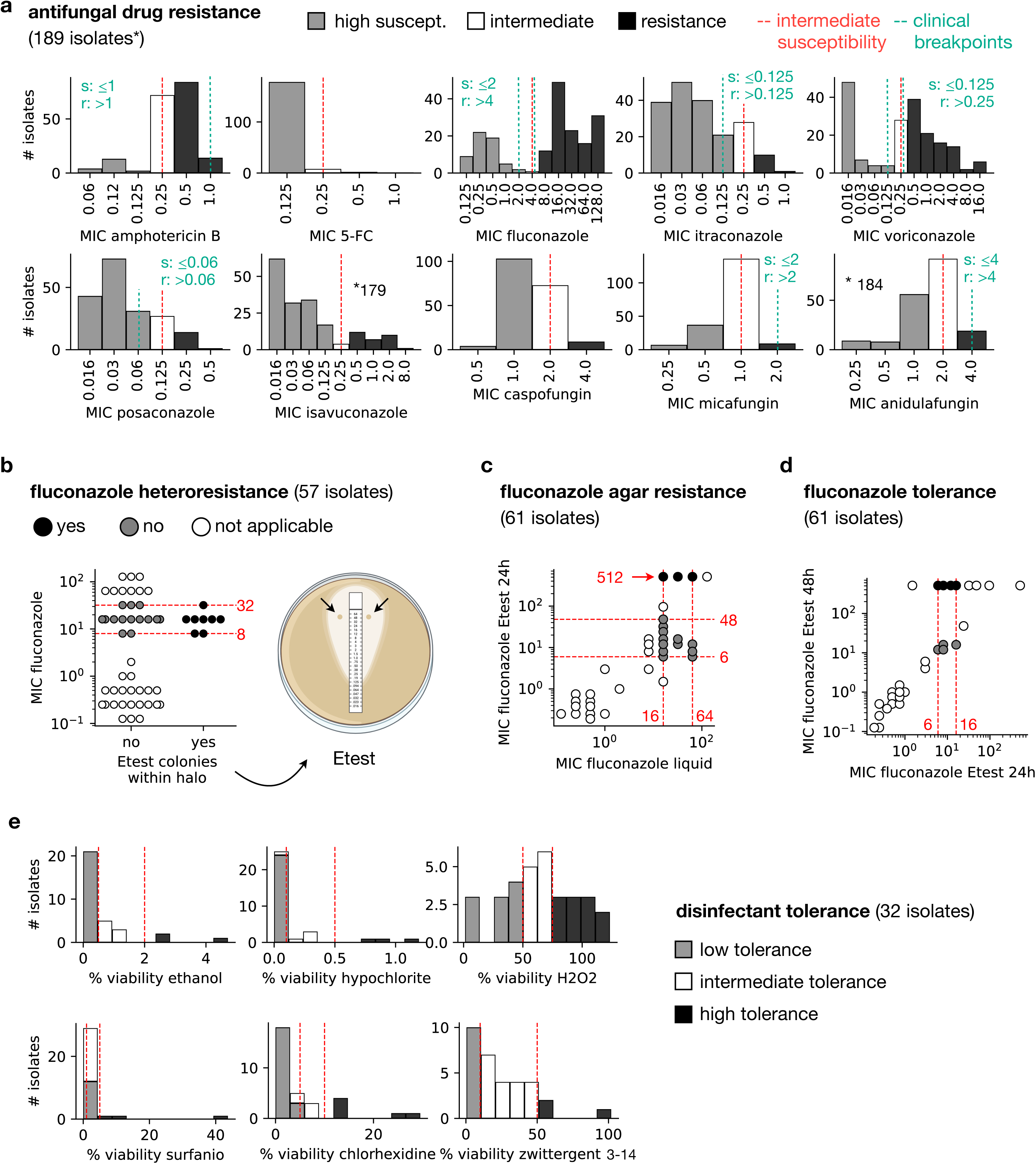
Definition of binary phenotypes measured in the lab. (a) Distribution of MIC values for all ten antifungals considered in this work, with bar colors reflecting the binary high susceptibility / intermediate susceptibility / resistance phenotypes. The red lines indicate the intermediate susceptibility concentration, while the green ones indicate the clinical breakpoints from https://www.eucast.org/astoffungi/clinicalbreakpointsforantifungals (EUCAST v10.0). Check **Methods** for more details on why the clinical breakpoint does not always match the used threshold that defines intermediate susceptibility. The numbers of isolates refer to those where we have antifungal susceptibility data. To see the numbers of strains with / without each phenotype, refer to Fig. 2. (b-d) Binary definition of fluconazole heteroresistance, agar resistance and tolerance, according to E-test measurements. The red lines indicate MIC thresholds used to focus comparisons on meaningful isolates (see **Methods**). (e) The same as in (a), but for the disinfectant tolerance phenotypes.

**Extended Data Fig. 5.**
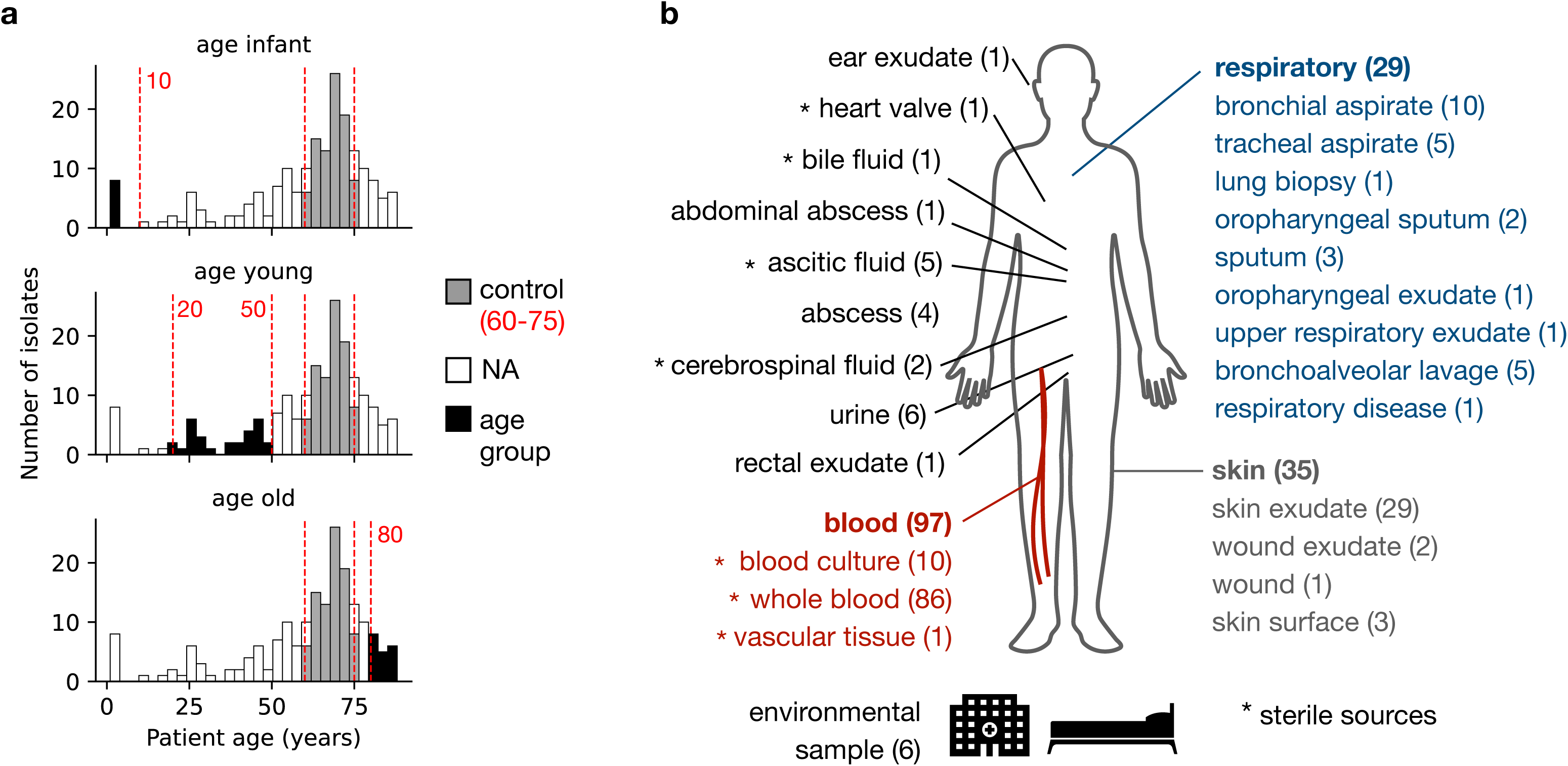
Definition of binary clinical features. (a) Patient age distribution, showing the thresholds used to define each of the age groups (infant, young, old; black bars) vs the 60-75 age of control patients (gray bars). Note that this distribution includes 187/189 analyzed samples, as for two patients we had no age information. (b) Isolation source of all strains sequenced here. To gain power to detect associations, we grouped certain sources together (respiratory, blood, skin and sterile), as shown in the color groups and asterisk. See **Methods** for more details.

**Extended Data Fig. 6.**
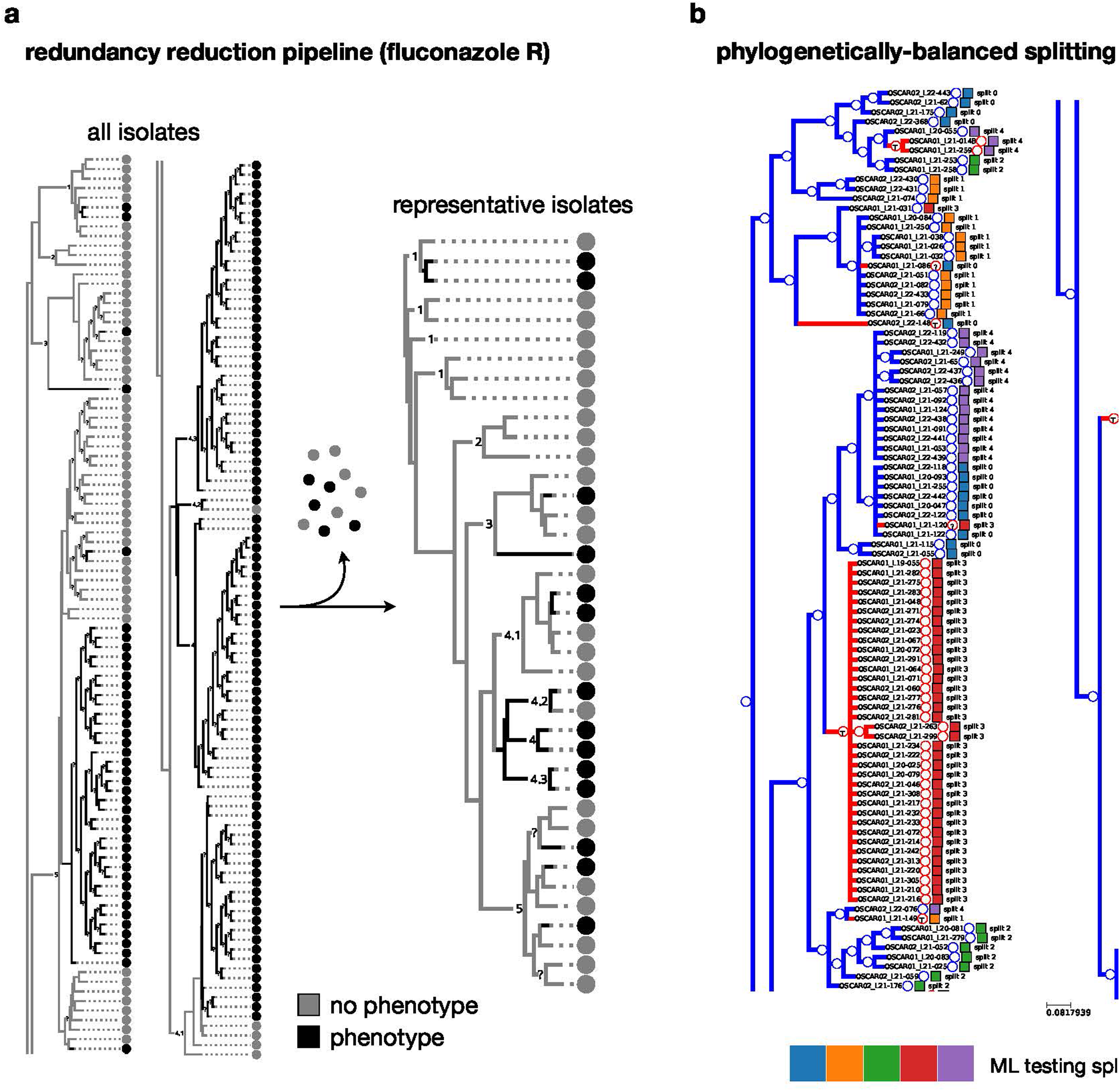
Pipelines for phylogenetic balancing in GWAS and ML analyses. (a) To perform a phylogenetically balanced GWAS and ML analyses we designed a redundancy reduction pipeline that takes all strains for which a phenotype is measured (left tree) and removes some strains to keep the representative ones (right tree). The example shown is the one for fluconazole resistance. This pipeline is implemented in the get_balanced_samples.py script of the ancestral_GWAS toolkit (see **Methods**). (b) We used this balancing pipeline to perform phylogenetically-balanced splitting in our ML analyses of all samples. The squares show, for fluconazole resistance, to which 1-5 split each strain was assigned. Note how this ensures that training / testing data are from different clades, as a way to reduce overfitting. The red / blue circles indicate susceptible / resistant strains.

**Extended Data Fig. 7.**
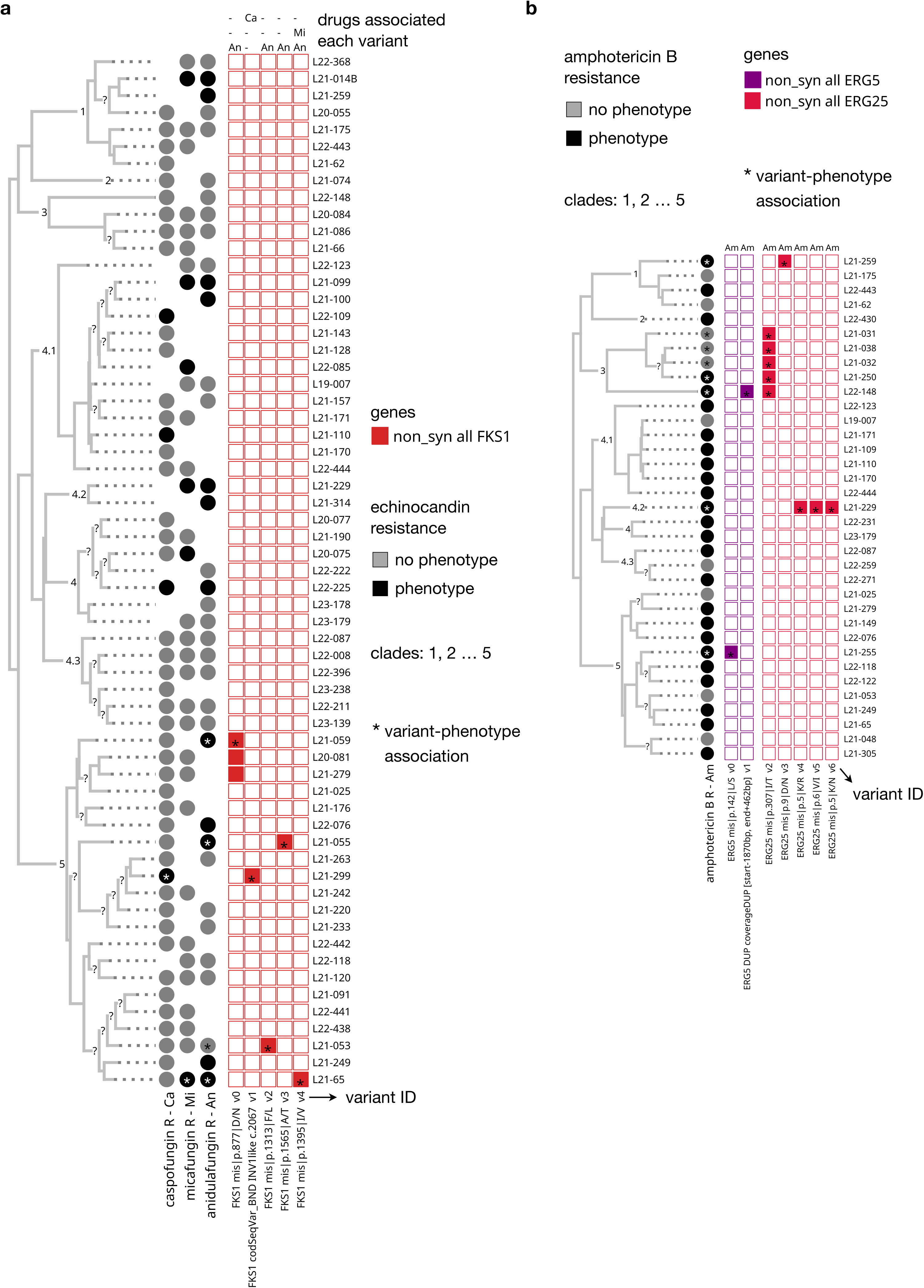
Evaluation of the known mechanisms of echinocandin and amphotericin B resistance. Equivalent to Fig. 3, but for echinocandins (a) and amphotericin B (b). Note that we could not find any such variants for *FKS2* or *ERG3* in echinocandins, nor for *ERG2*, *ERG3* or *ERG6*.

**Extended Data Fig. 8.**
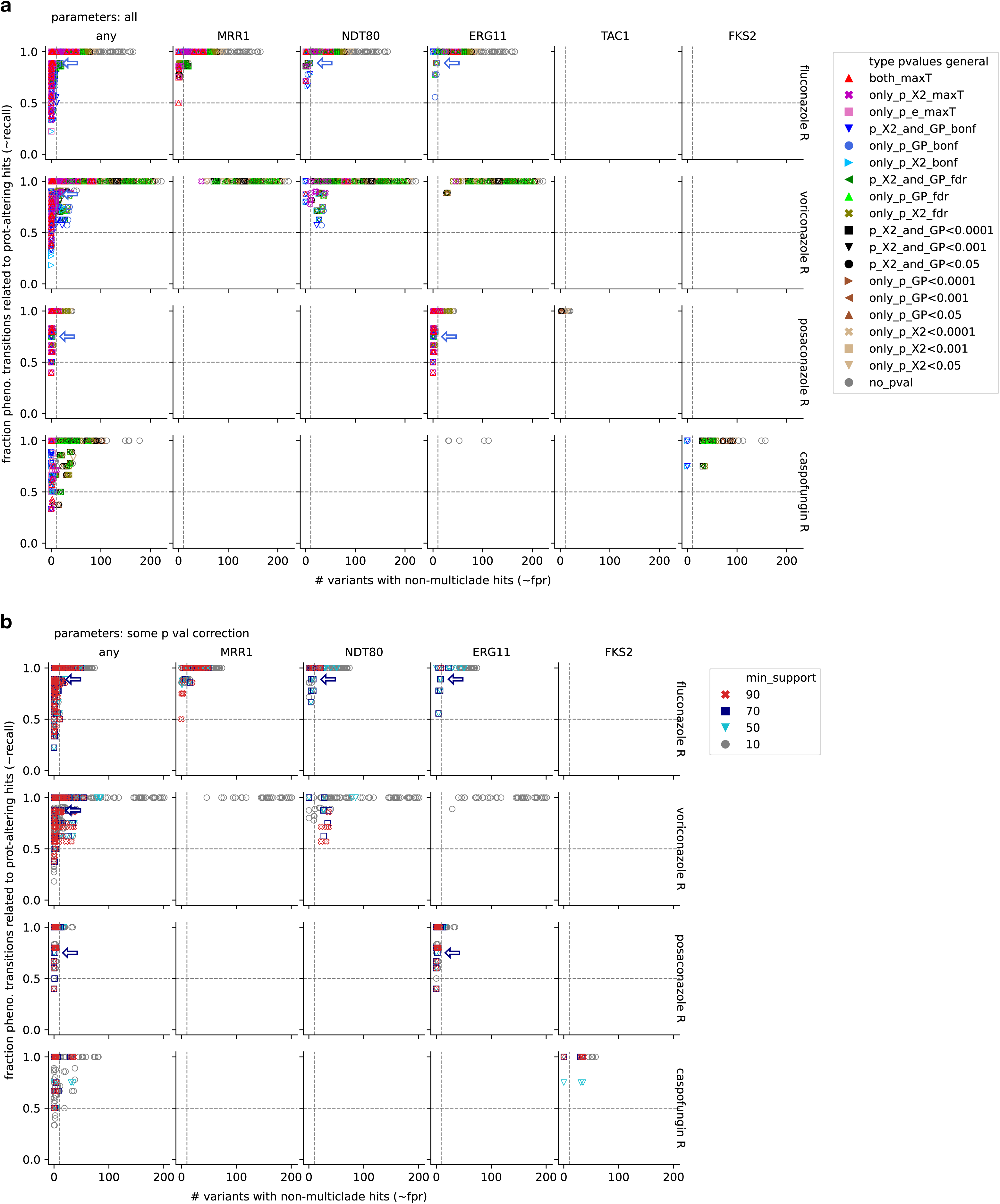
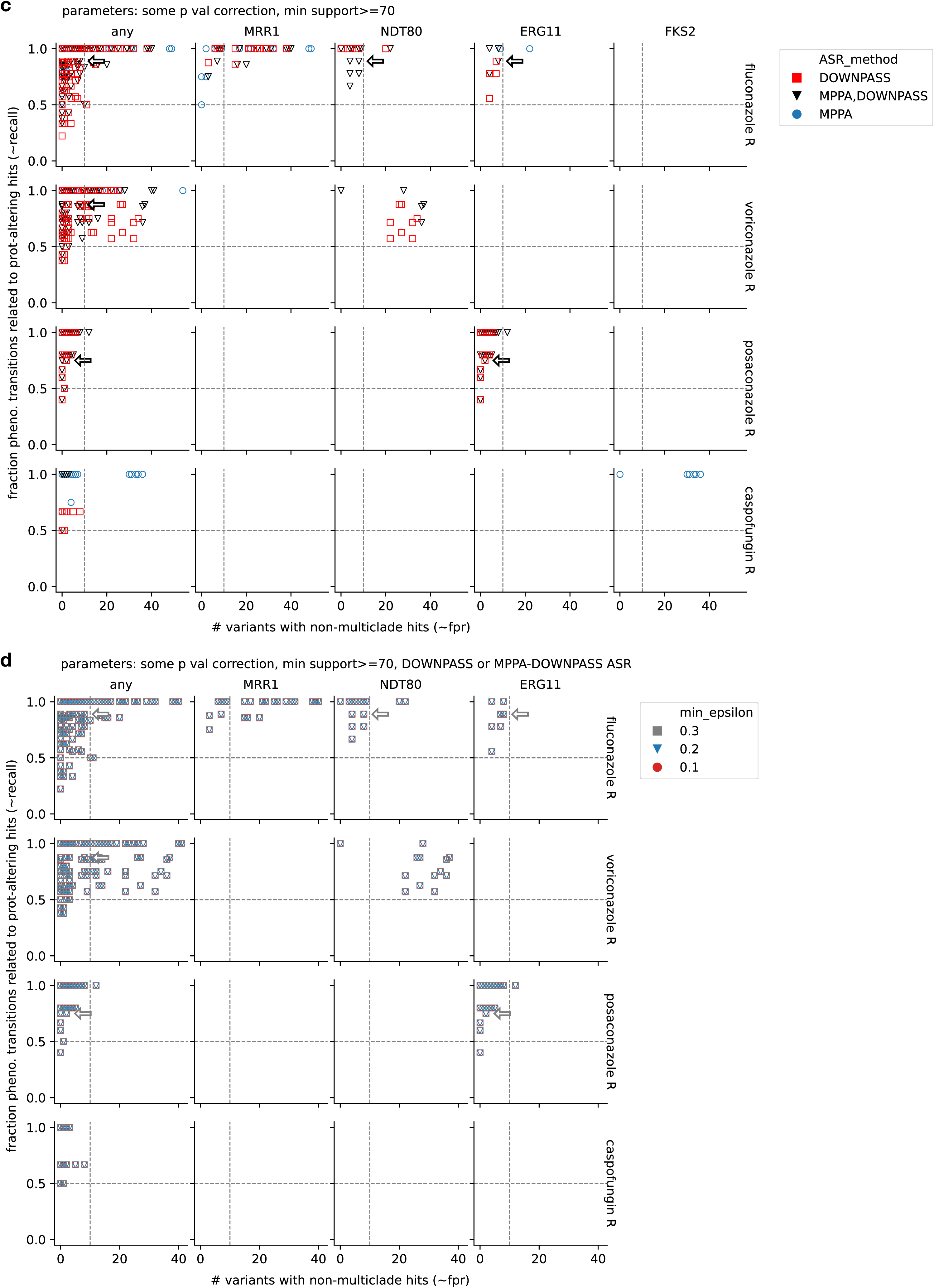
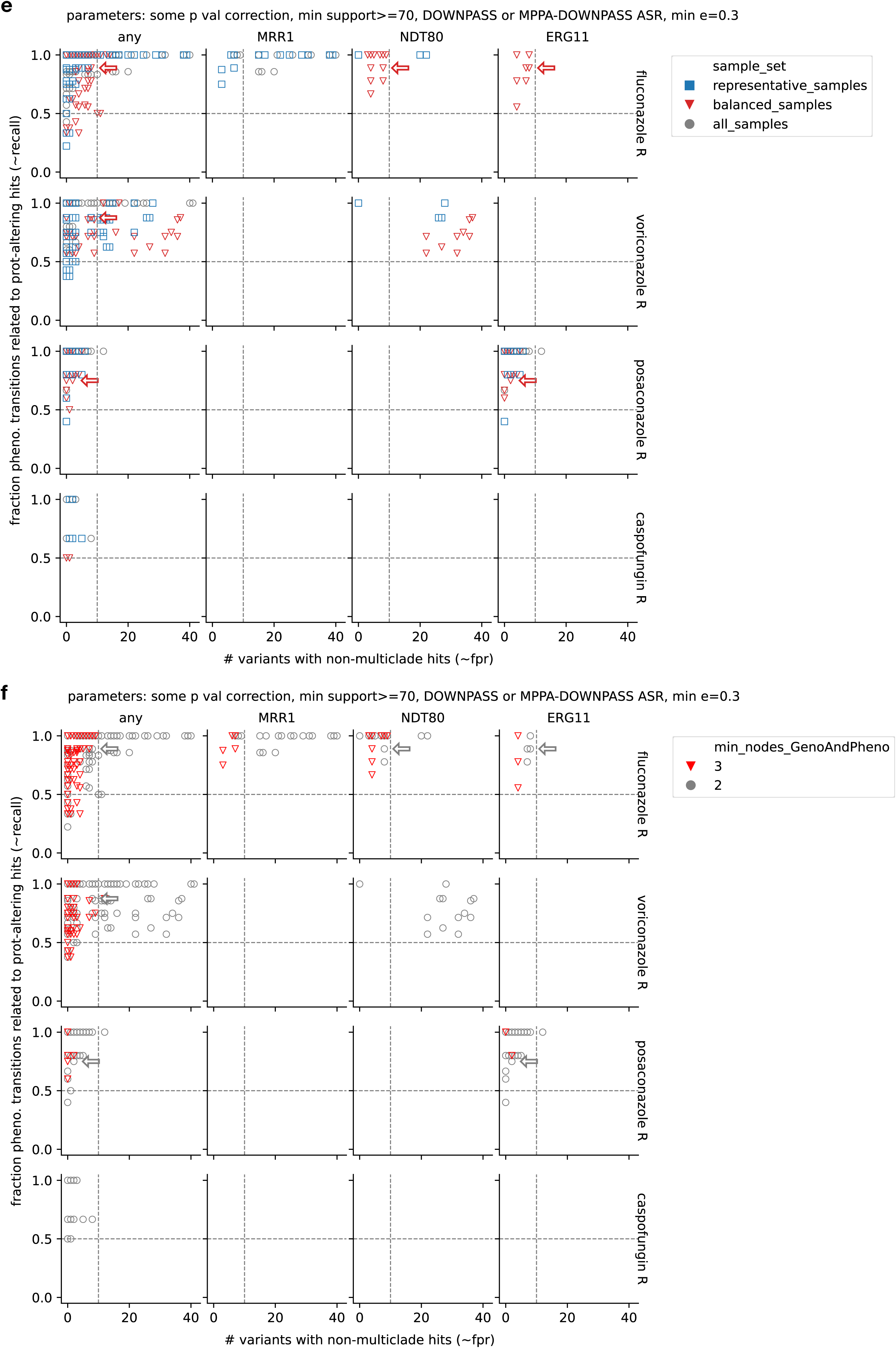
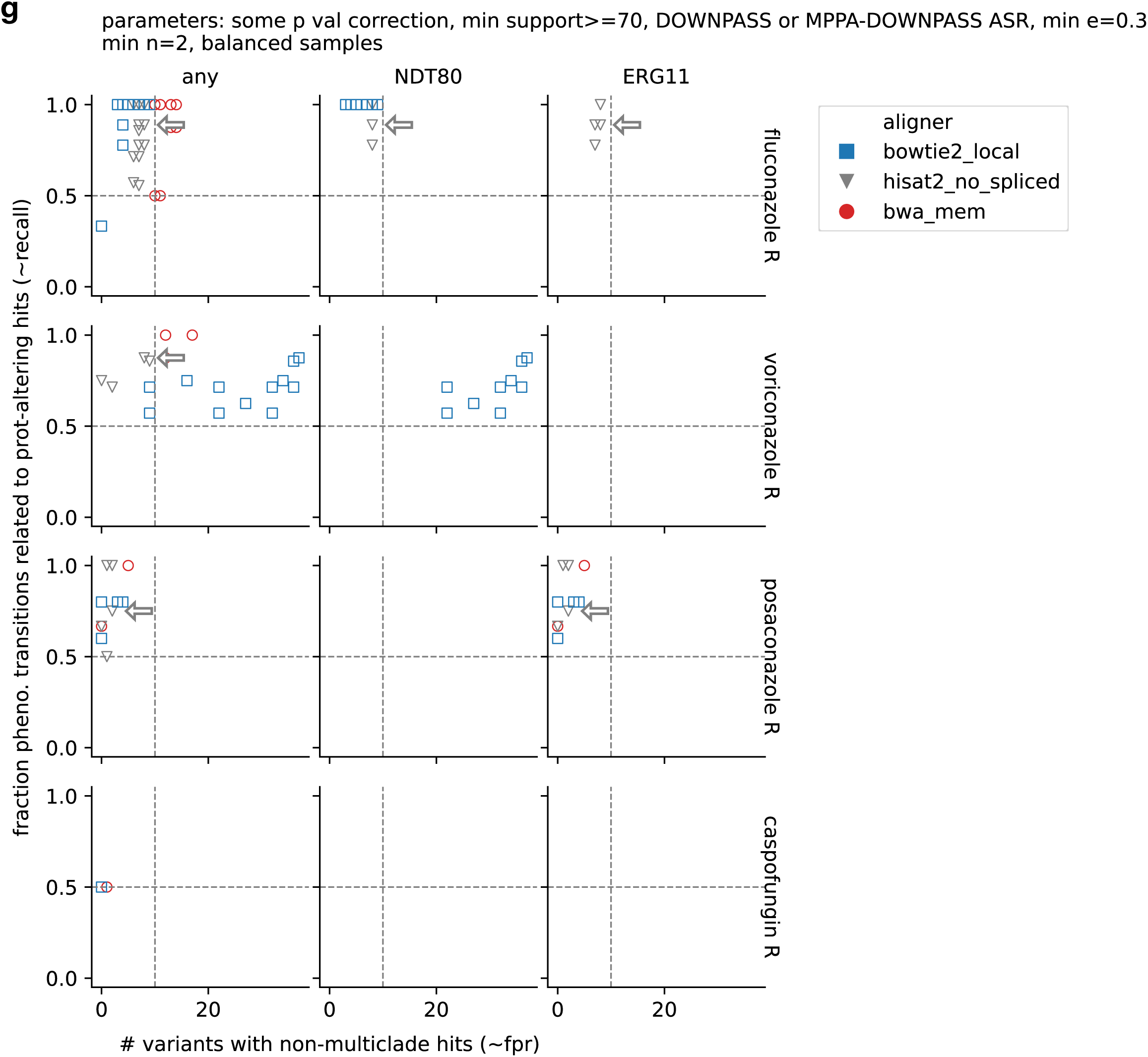
Exploration of different GWAS parameters. We applied sequentially filters to keep the important ones. The panels show this sequential filter exploration. The scatter points are plotted putting on top the preferred parameters, ordered as in the legend. The arrow represents the location of the final benchmarked parameters (see **Methods** and **Supplementary Results**).

**Extended Data Fig. 9.**
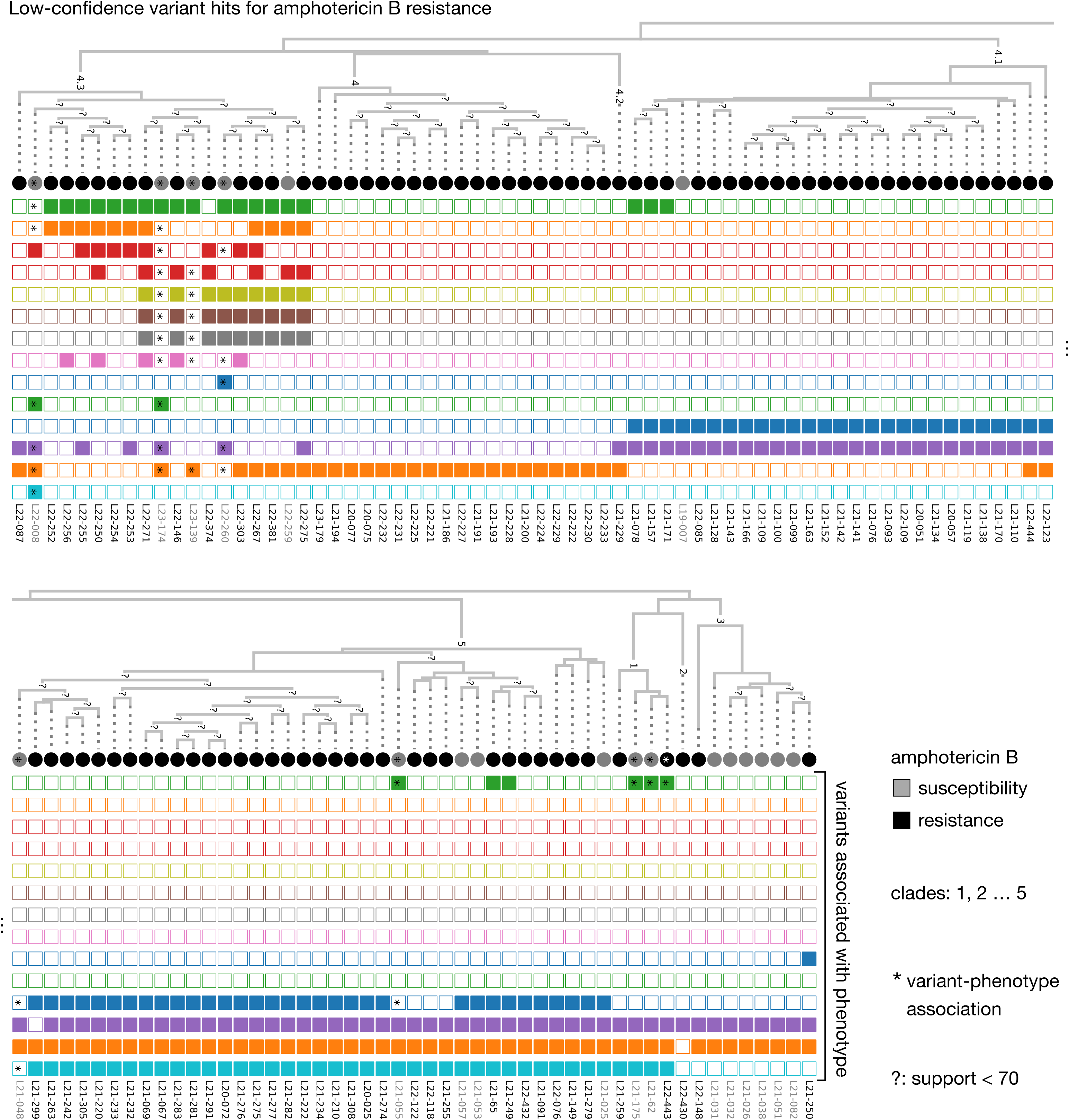
Example low-confidence GWAS variant hits. To justify our approach of treating GWAS variant hits as false positives we visualized those passing low-confidence filters for amphotericin B resistance as in Fig. 3, which include 14 variants (**Supplementary Table 5**). The asterisks represent variants and samples for which one of the variants’ transitions is correlated to a transition in the phenotype. The “?” refers to nodes with a support <70.

**Extended Data Fig. 10.**
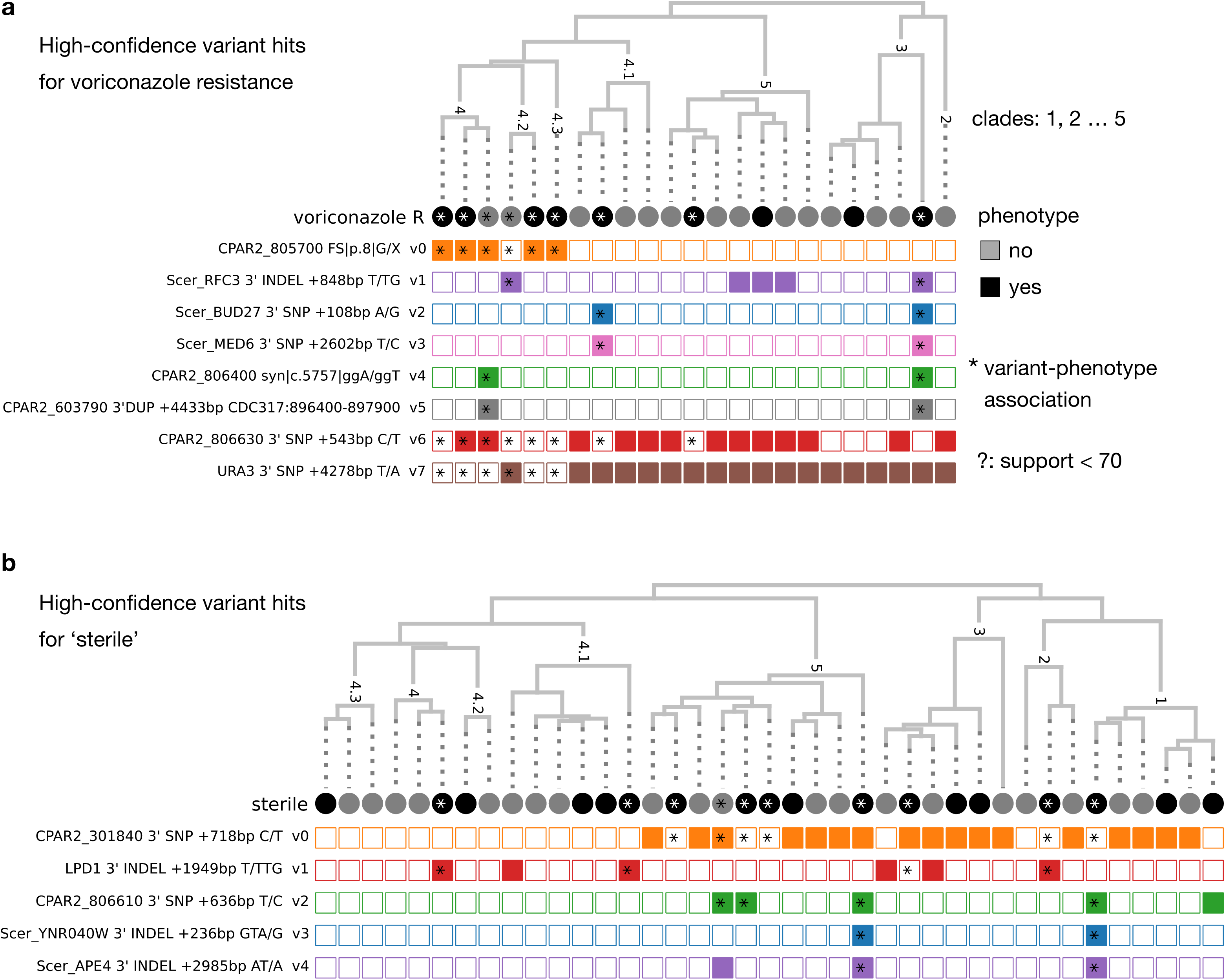
GWAS high-confidence variant hits for phenotypes in which recombination is unclear. Presence / absence pattern of high-confidence GWAS variant hits for ‘voriconazole resistance’ and ‘source sterile’, represented as in **Extended Data** Fig. 9.

**Extended Data Fig. 11.**
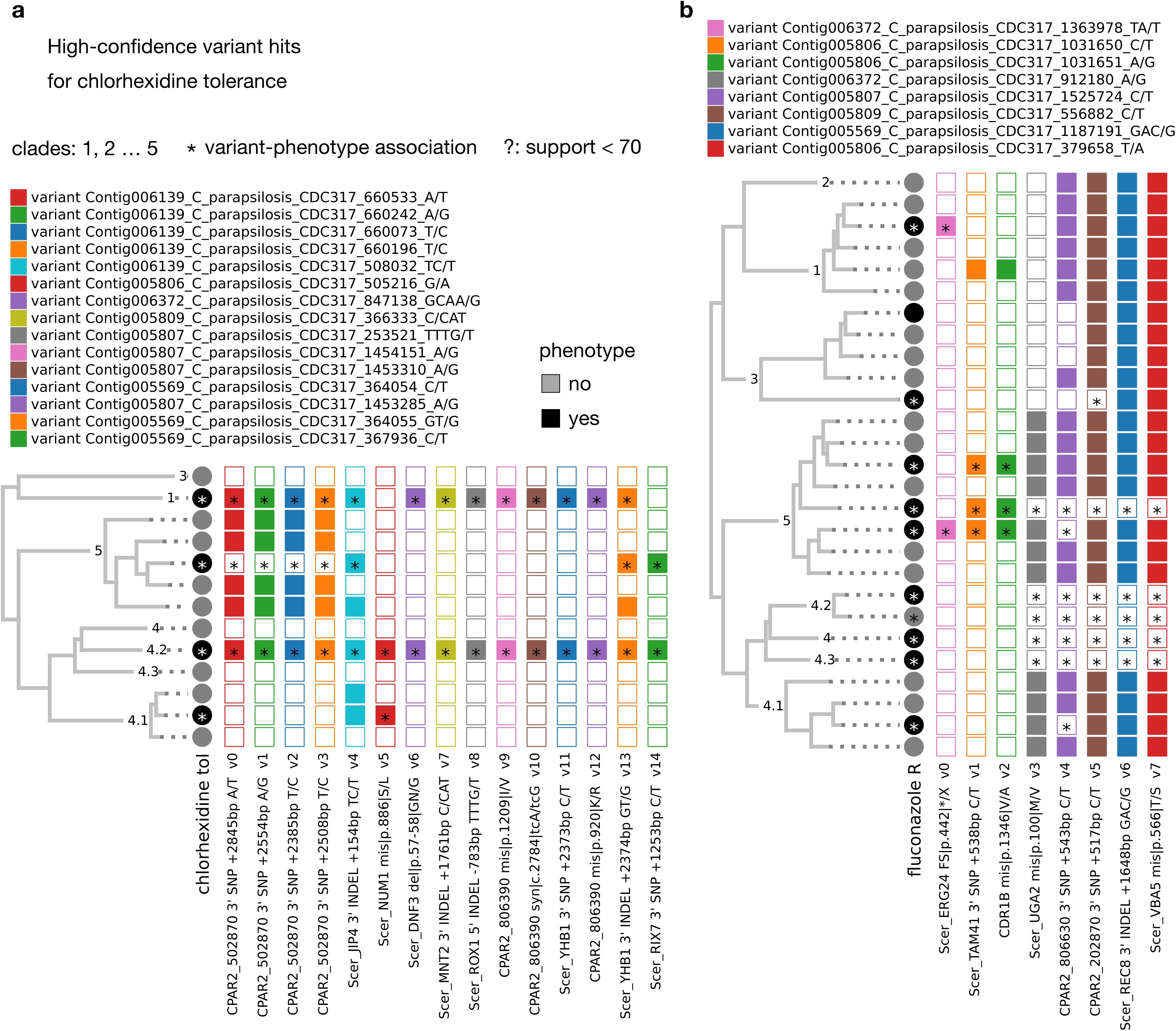
GWAS high-confidence variant hits for phenotypes in which recombination is plausible. Presence / absence pattern of high-confidence GWAS variant hits for ‘chlorhexidine tolerance’ and ‘fluconazole resistance’, represented as in **Extended Data** Fig. 9.

**Extended Data Fig. 12.**
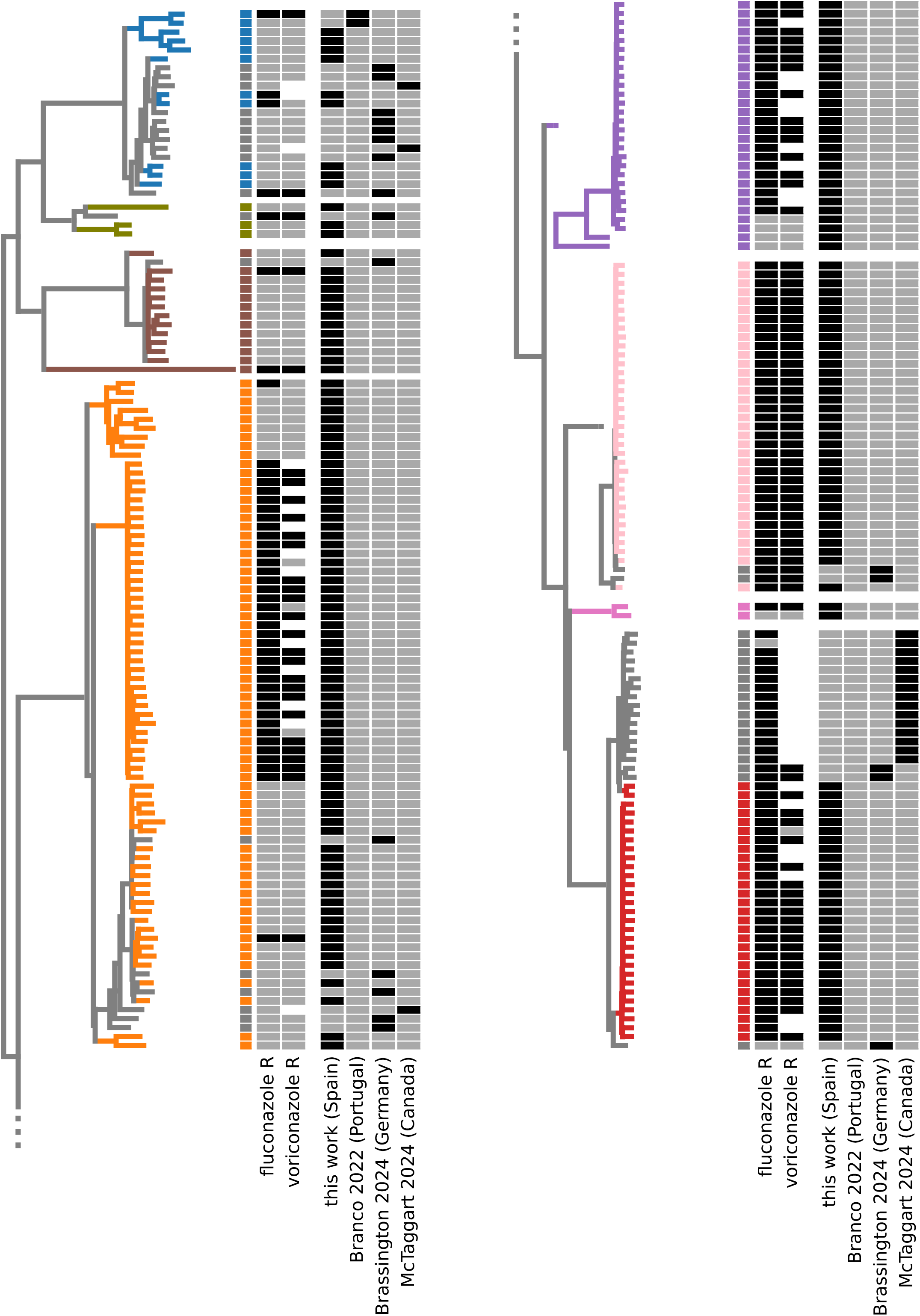
Phenotypic distribution of the strains used for ML validation. Strains tree for all isolates with clear 0/1 fluconazole resistance, including those sequenced here and those used for the ‘validation’ ML testing strategy (see **Main text**). This includes 188 / 189 strains sequenced here and 40 extra ones previously phenotyped in Branco et. al.^79^, Brassington et. al.^37^ and McTaggart et. al.^38^. Additionally, the heatmap shows the voriconazole resistance profile for our isolates and the subset of the extra strains where voriconazole susceptibility data was available. The colors of clades and phenotypes match those in Fig. 2. Note that some strains have no clade coloring as they were added to the analysis specifically for the ML validation (see **Methods**).

**Extended Data Fig. 13.**
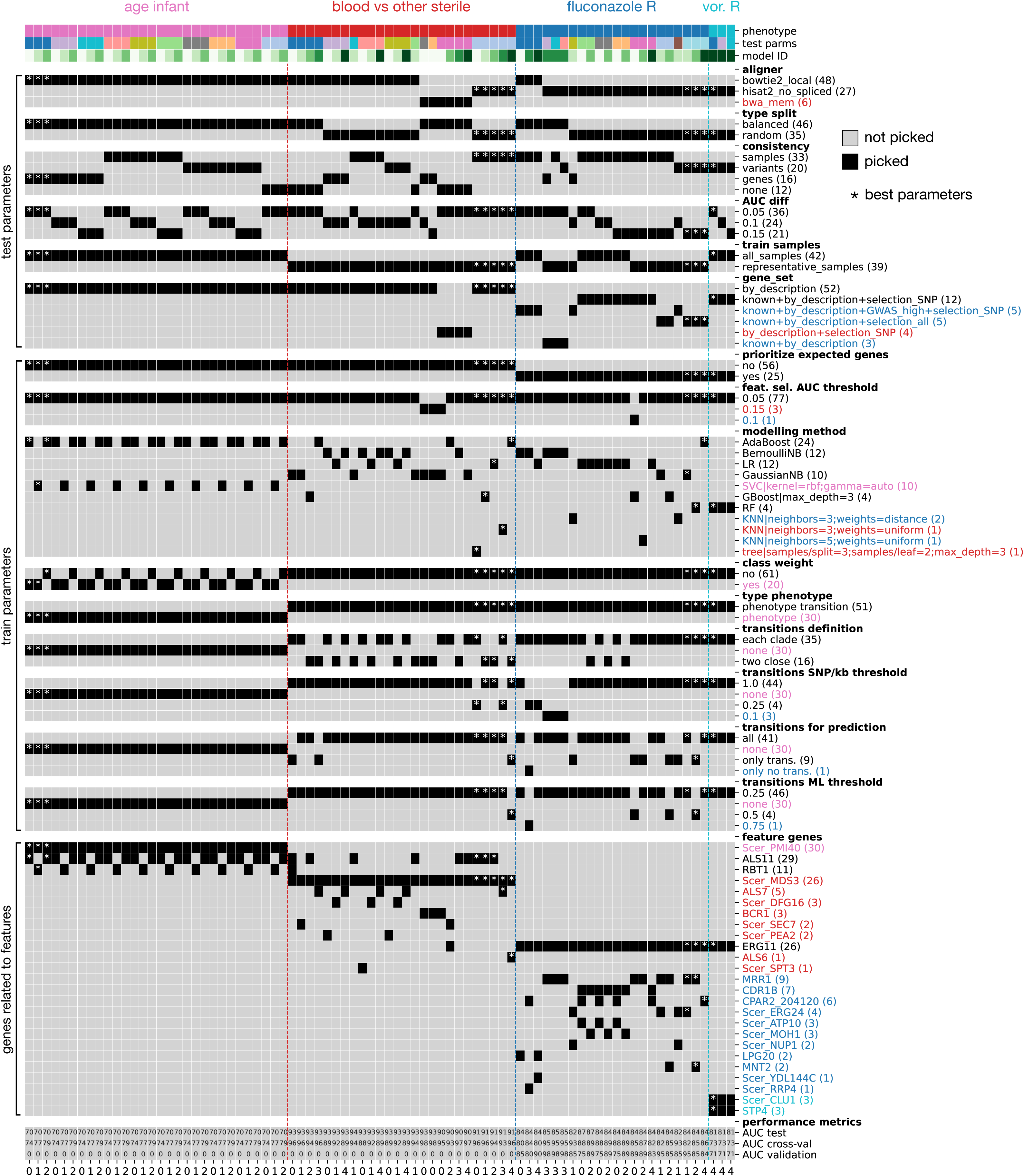
Chosen parameters, genes related to selected features, and performance for all top models. The performance metrics are the ROC AUC values as yielded by different testing strategies (see Fig. 5 **d**). Also, the colors indicate parameters or genes that are unique to one of the four phenotypes.

**Extended Data Fig. 14.**
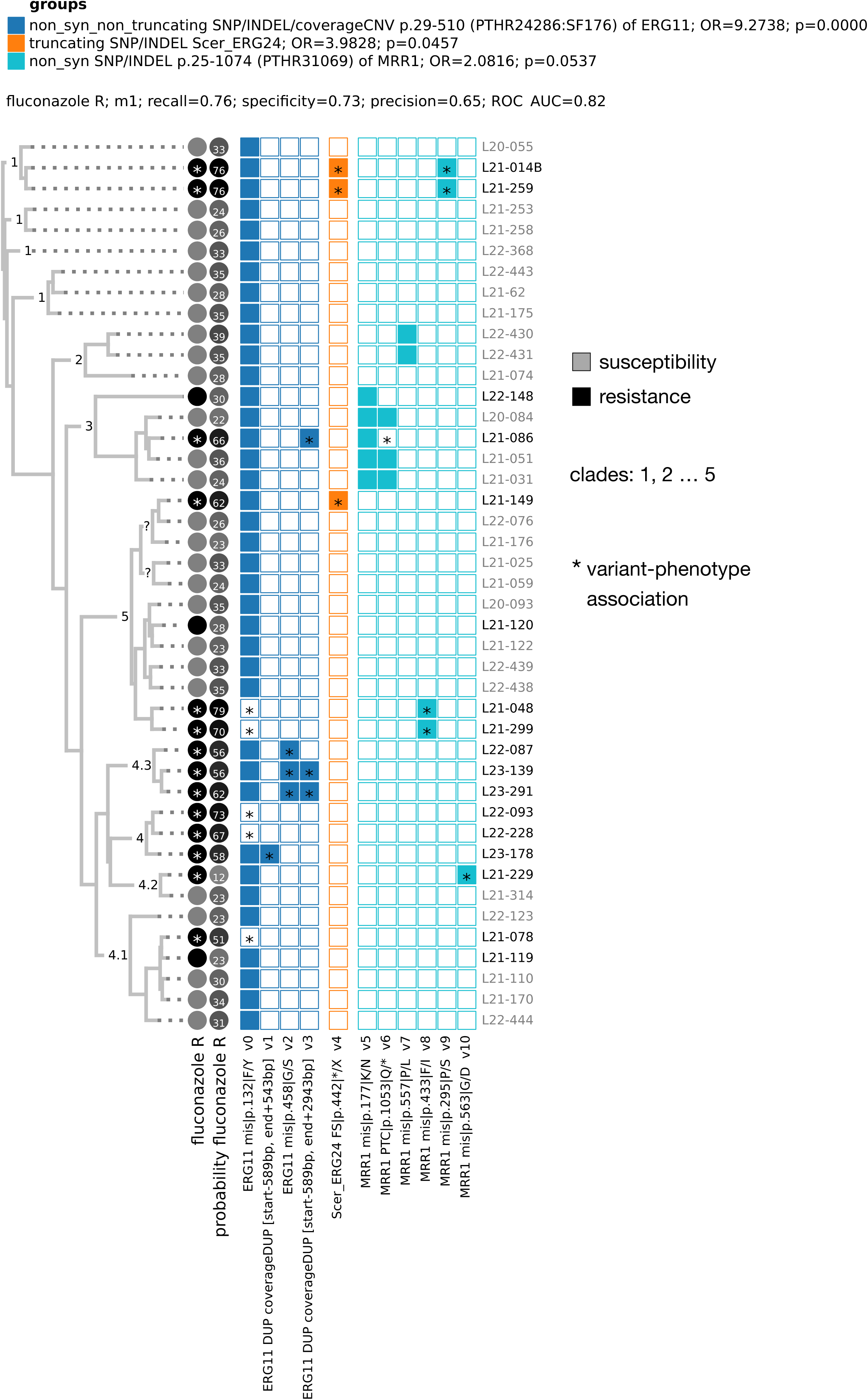
Features used by the best top model for fluconazole resistance. This representation is similar to Fig. 6, but showing the predicted phenotype probability. Note that the prediction was based on phenotype transitions (see **Extended Data** Fig. 15), and the indicated OR and p values indicate, for each feature, the association statistics between the feature transition and the phenotype transition. The “predicted phenotype” is based on a threshold balancing sensitivity and specificity (see **Methods**). Among the different m1-m5 models yielded by the best parameters for fluconazole, this is the one that has features more consistent with the most commonly used genes for fluconazole models (Fig. 6b**,c**). The probability reflects the two decimals of a 0-1 probability.

**Extended Data Fig. 15.**
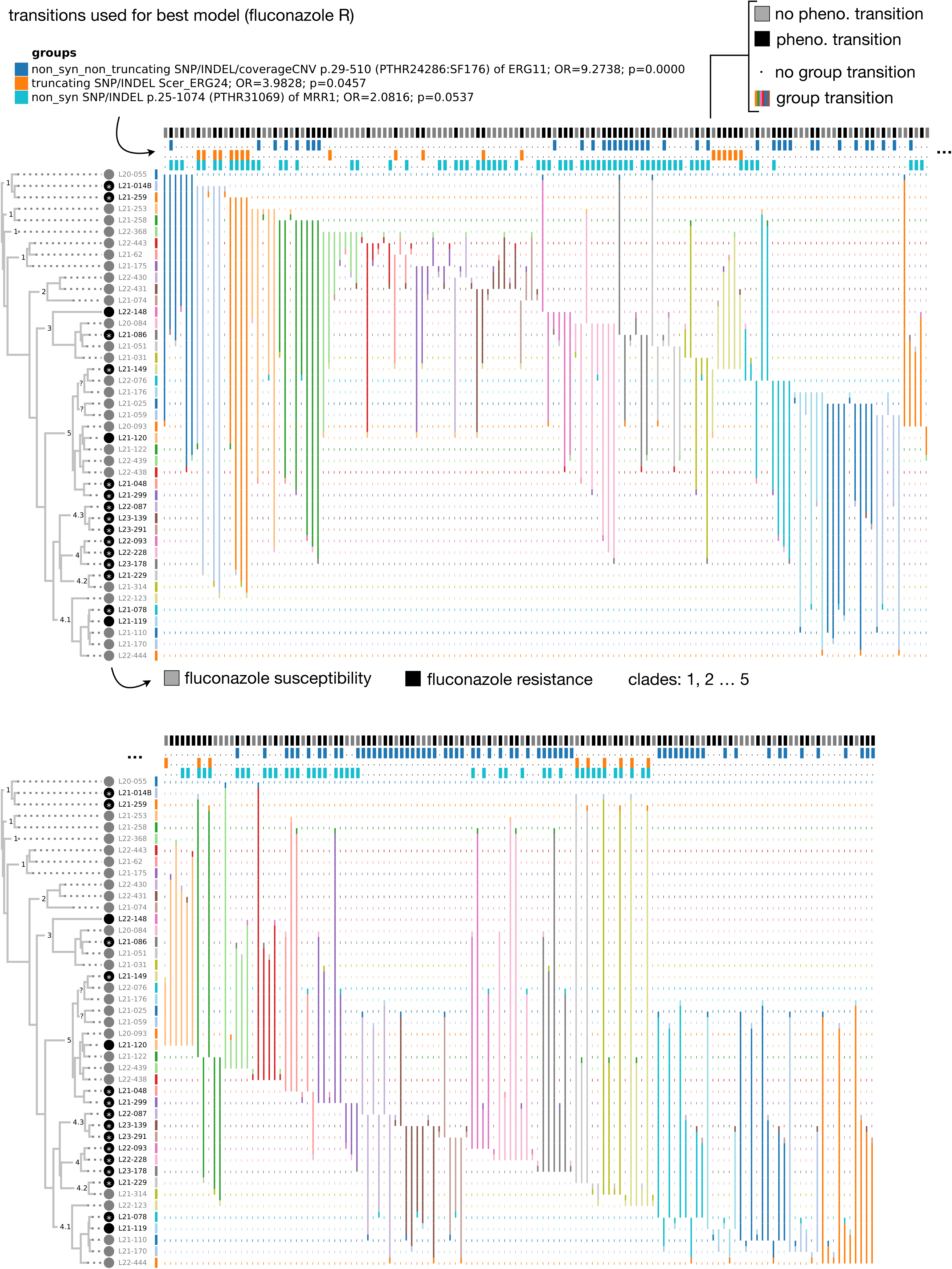
Transitions used for the best model for fluconazole resistance. This is related to **Extended Data** Fig. 14. The colors in rows indicate each sample, sorted by the phylogeny. The colors in columns indicate the group transitions (change in any variant of the group). The vertical lines indicate the pairs of strains compared for each transition / no-transition instance (columns).

**Extended Data Fig. 16.**
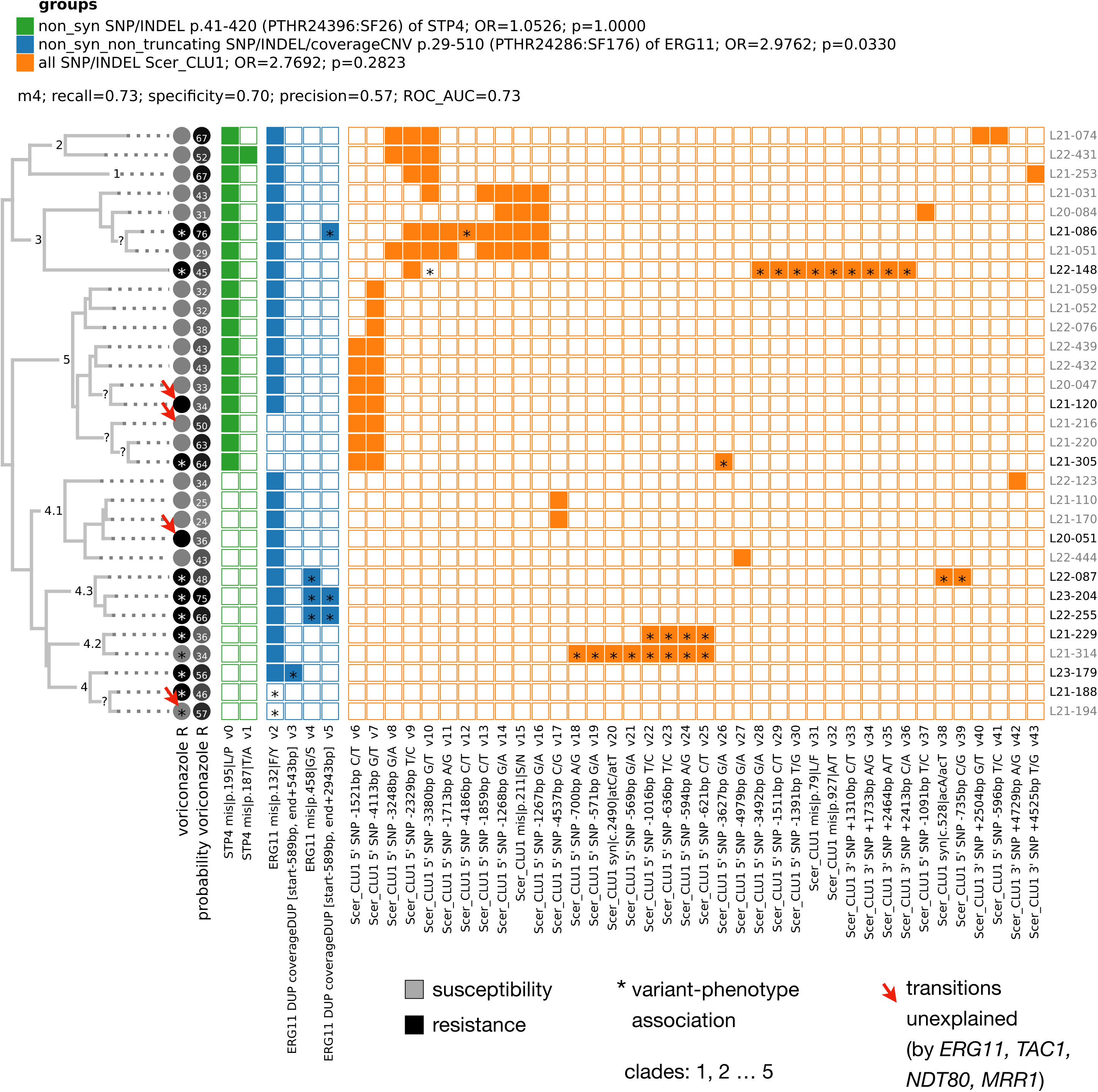
Features used by the best top model for voriconazole resistance. Equivalent to **Extended Data** Fig. 14 but for the top model for voriconazole. Note that all top models for this phenotype used *STP4*, *ERG11* and *Scer_CLU1* (Fig. 6a**, Extended Data** Fig. 13), so that this model is representative of the others. The red arrows indicate resistance transitions that are not explainable by the expected genes (Fig. 3).

**Extended Data Fig. 17.**
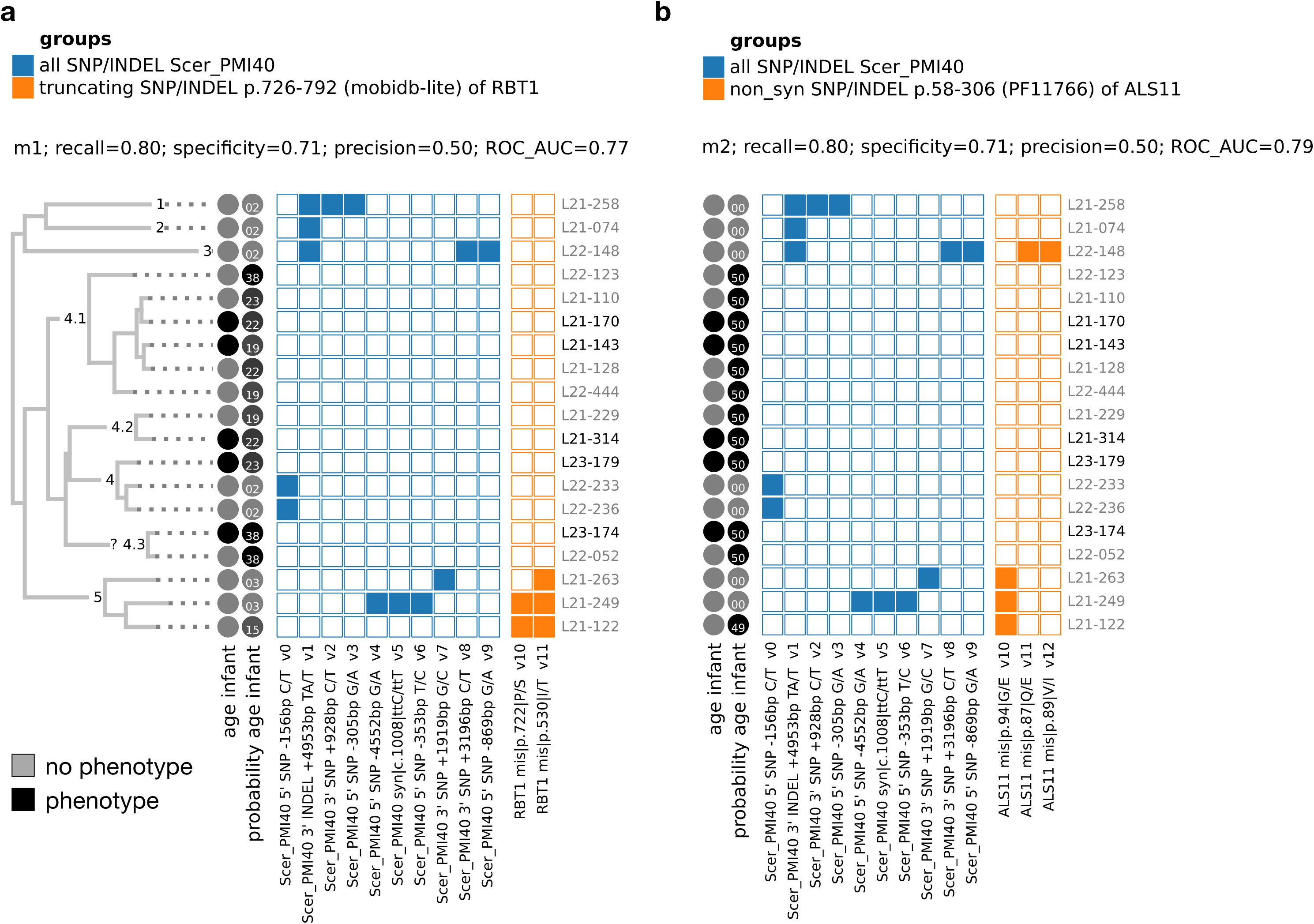
Features used by the representative top models for ‘age infant’. Equivalent to **Extended Data** Fig. 14 but for ‘age infant’. To visualize representative models among the top ones we took advantage of the fact that all of them used features related to *Scer_PMI40* and either *ALS11* (20/30 top models) or *RBT1* (10/30 top models) (Fig. 6a**, Extended Data** Fig. 13). Also, among the best parameters we had models using these combinations (m1 and m2, respectively, **Extended Data** Fig. 13), indicating that these are representative of each set of genes. The panels (a) and (b) show the results of each of the m1/m2 models.

**Extended Data Fig. 18.**
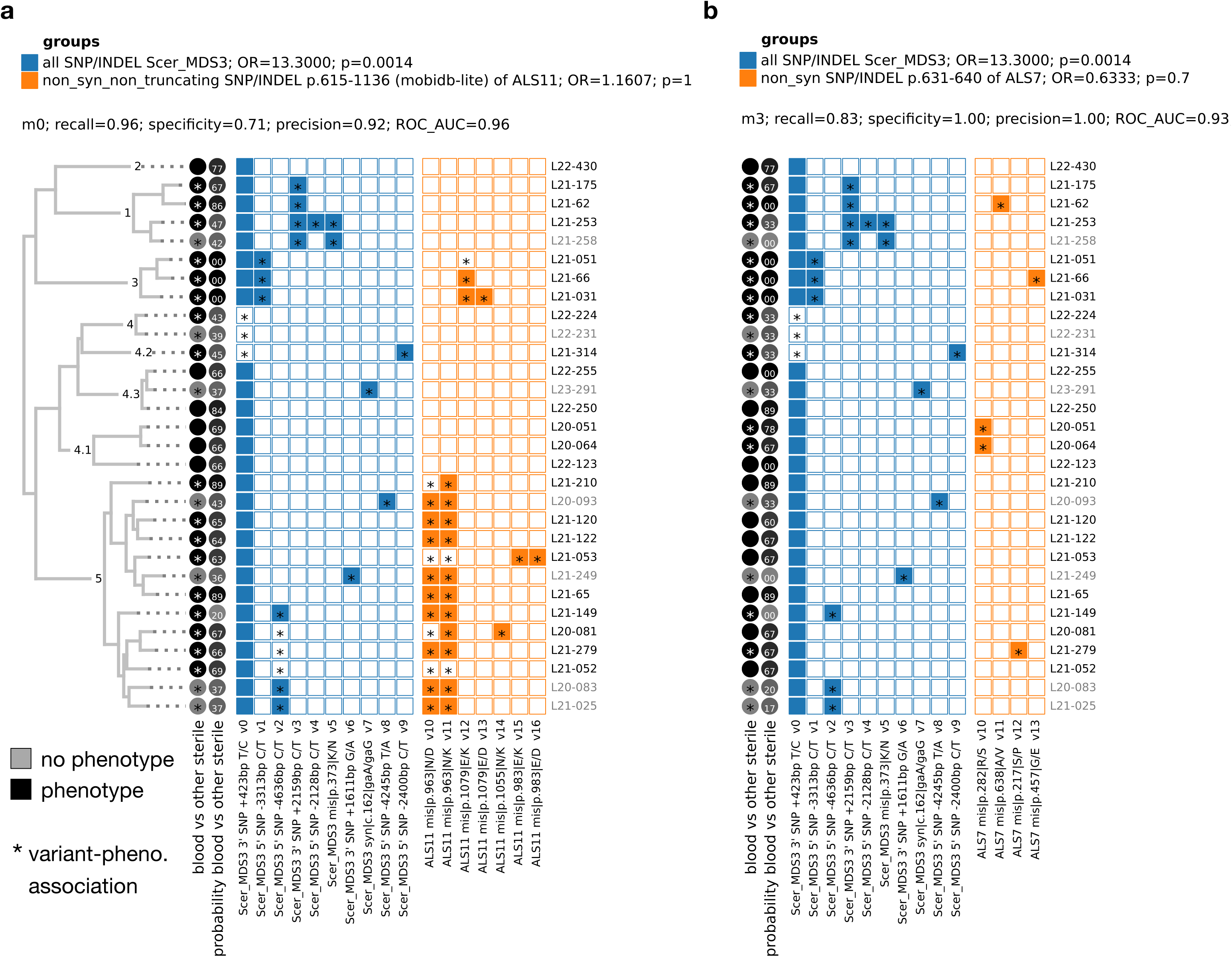
Features used by the representative top models for ‘blood vs other sterile’. Equivalent to **Extended Data** Fig. 17 but for ‘blood vs other sterile’. For this phenotype, all of the 26 top models used *Scer_MDS3* combined with one (24/26) or two (2/26) additional genes (**Extended Data** Fig. 13). Out of these additional genes, we focused on those that are used by at least three top models, including *ALS11*, *ALS7*, *BCR1* and *Scer_DFG16*. Also, given that we had top models found with all three aligners (**Extended Data** Fig. 13), we focused on genes found by at least two aligners, including *ALS11* and *ALS7*. Among the best parameters we had models using these combinations (**Extended Data** Fig. 13): *Scer_MDS3* and *ALS11* (a) or *Scer_MDS3* and *ALS7* (b).

**Extended Data Fig. 19.**
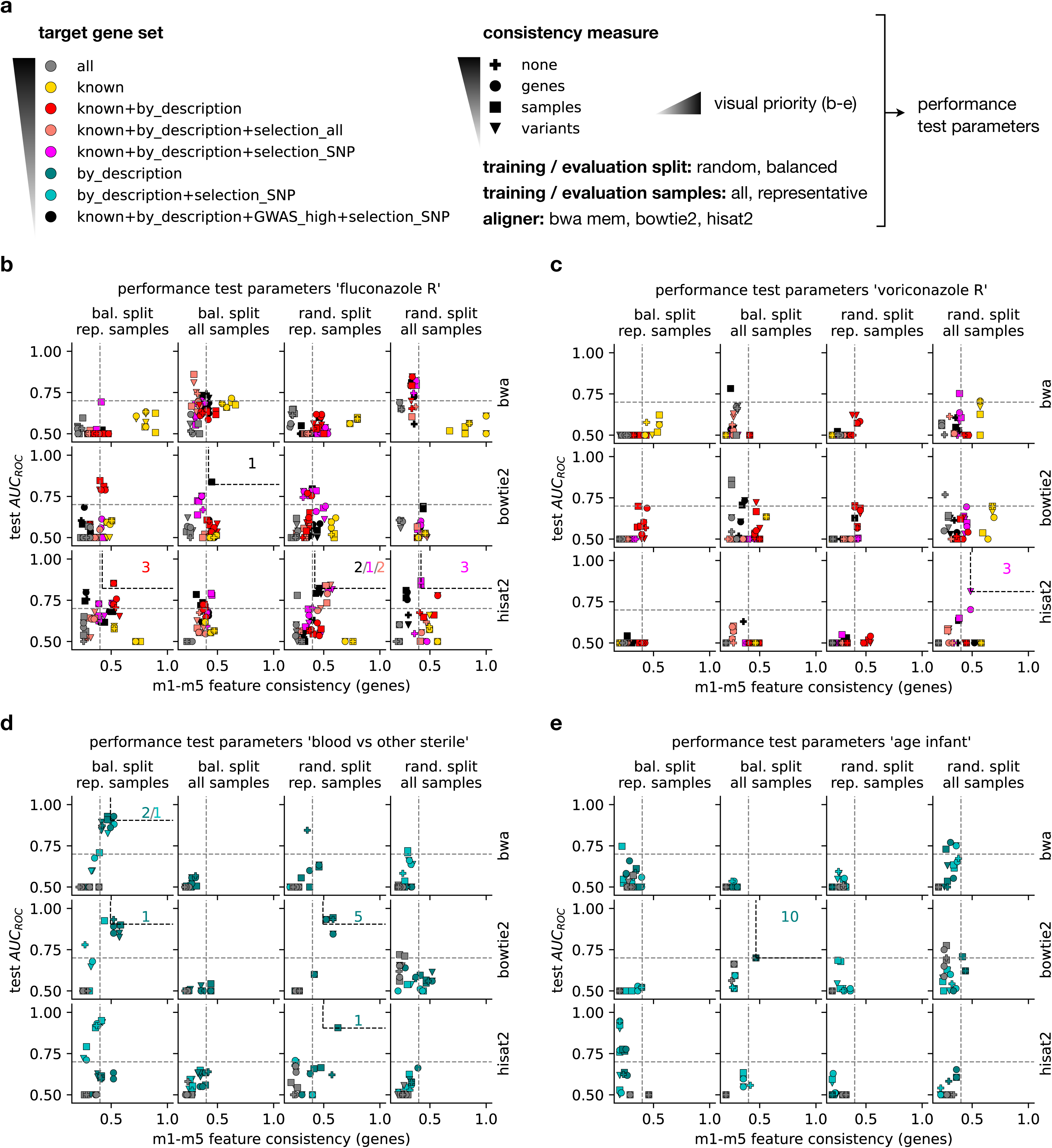
Performance of test parameters for each phenotype. (a) To understand the effect on performance of different test parameter values (Fig. 5b) we visualized their predictive performance (test AUC) and m1-m5 feature consistency, splitting the results by gene set (colors), consistency measures (symbols), train/evaluation split (columns), train / evaluation samples (columns) and aligner (rows). Only shown are parameter values that were taken in any of the top parameters. The gene set may be any combination of i) any gene, where initial feature selection is based on GWAS (all), ii) genes expected to be related to azole resistance, inlcuding “ERG11”, “ERG3”, “TAC1”, “MRR1”, “NDT80” and “UPC2” (known), iii) genes with a functional description in SGD that is related to virulence or resistance (see **Methods**) (by_description), iv) genes with orthologs (in other *Candida* species) having signs of recent genomic selection by any variant, according to ^9^ (selection_all), v) genes with orthologs having signs of recent genomic selection by SNPs (selection_SNP) or vi) genes with orthologs in *C. albicans*, *C. glabrata* or *C. auris* with ‘high-confidence’ GWAS hits on azole resistance (GWAS_high). Also, note that we only show certain gene sets of interst, including i) all, ii) known and iii) those used among top parameters. To enable visual comparisons we plotted scatterplots using a stacking priority as defined by the gray colorbar. (b-e) Scatterplots with all the information described in (a), for each phenotype. The numbers in the top right indicate the numbers of top parameters that have each parameter combination.

**Extended Data Fig. 20.**
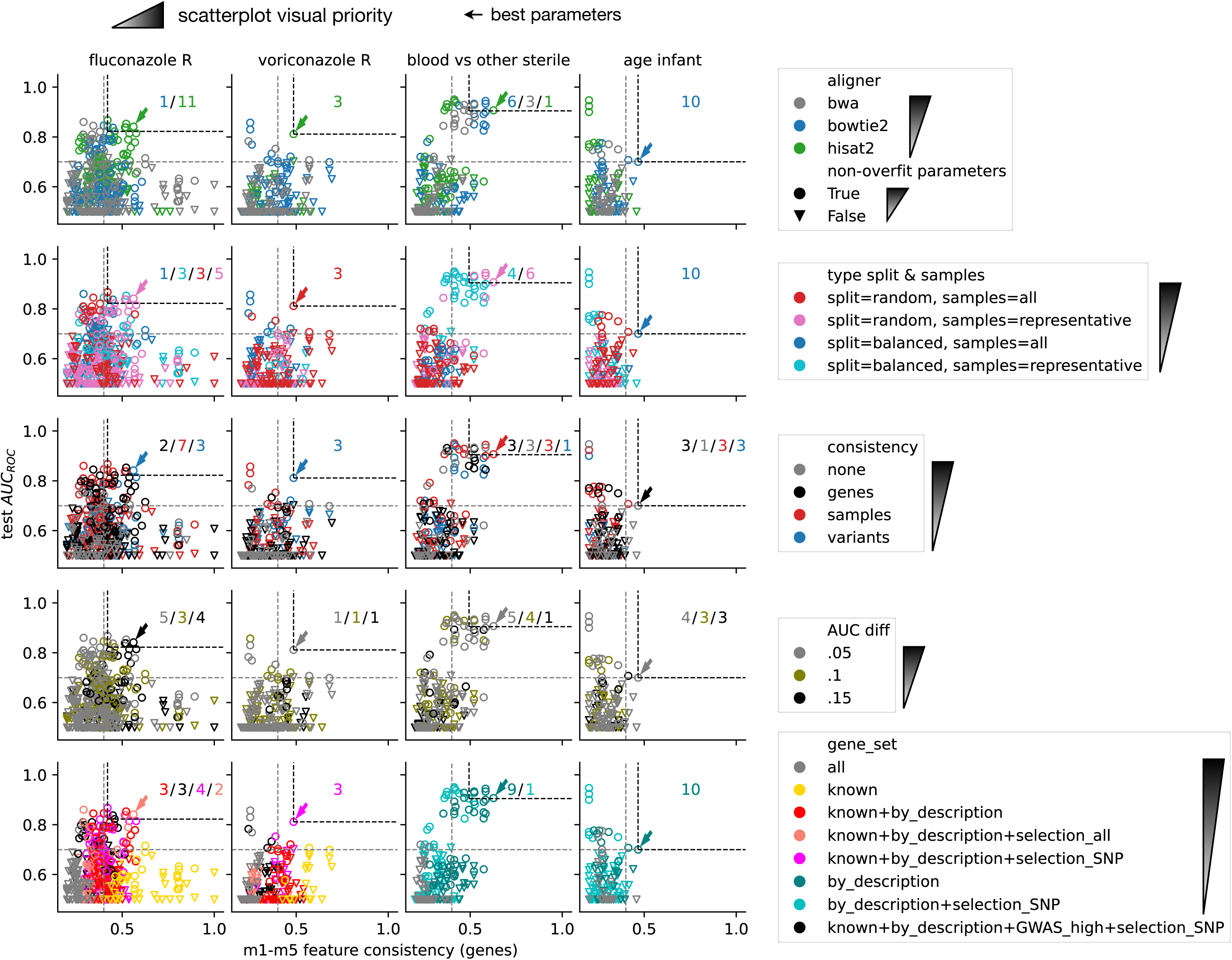
Performance of single test parameters for each phenotype. Performance of the test parameters as in **Extended Data** Fig. 19, but coloring each parameter values independently in rows. The symbols represent whether the parameters lack signs of overfitting (ROC AUC >= 0.7 on the training data and similar AUC in training and testing, see **Methods**), and the parameter values are plotted in the order shown in the legends

**Extended Data Fig. 21.**
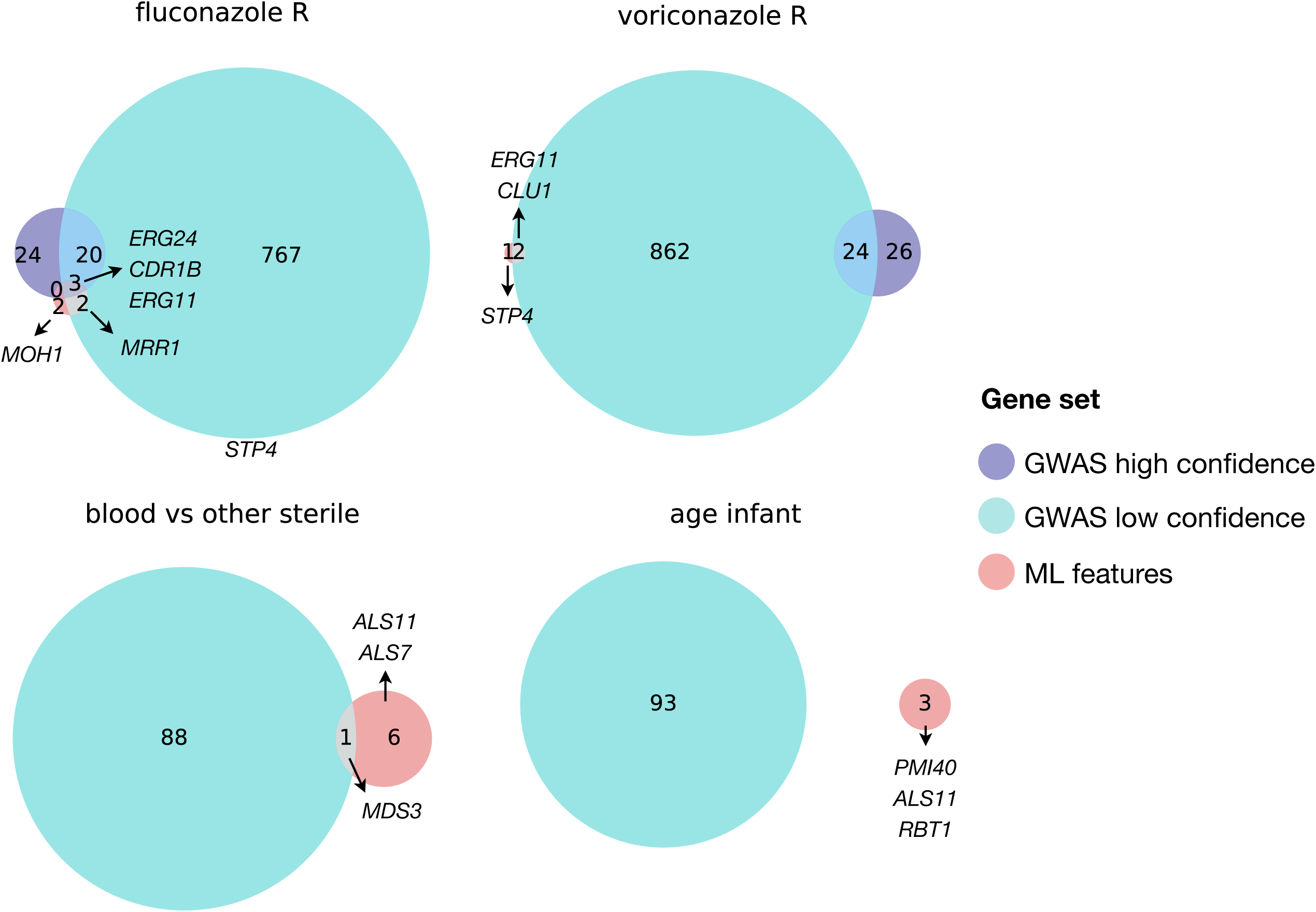
Overlap between the GWAS and ML genes. For each phenotype with considered ML predictors, this figure represents the overlap between the set of genes found by different approaches. The arrows point towards sets that include genes of interest.

**Extended Data Fig. 22.**
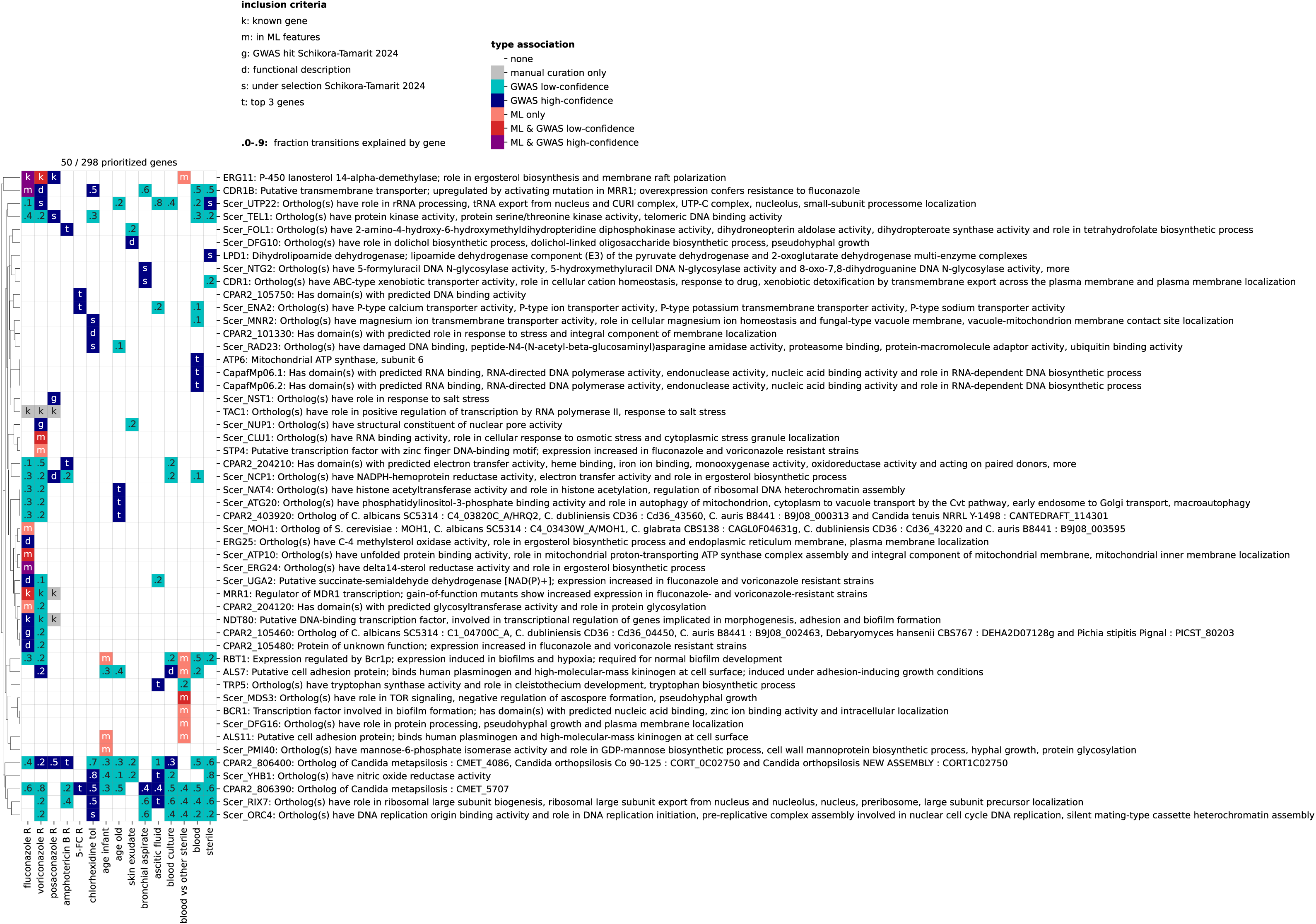
Descriptions of the functional genes. Equivalent to Fig. 7, but including the description of all genes.

