## Supplementary Material for "Dissection of genotype-phenotype relationships in *Candida parapsilosis* uncovers drivers of clinically-relevant traits"

### SUPPLEMENTARY RESULTS AND DISCUSSION

#### Choosing parameters for tree reconstruction

To select optimal parameters for SNP-based tree reconstruction we built six different strain phylogenies (**Fig. 1b**), varying i) the aligner used upstream of variant calling (bwa mem, hisat2 or bowtie2) and ii) the mode of tree reconstruction (diploid or diploid_homozygous). The mode refers to how we consider the zygosity of SNPs. In 'diploid' mode we used a common approach for diploid *Candida* species[1,2], by building a consensus tree from 100 resampled trees based on all homozygous SNPs and randomly selecting one of two alleles for heterozygous SNPs, i.e. haplotype sampling approach[3]. Conversely, in 'diploid_homozygous' mode we generate a single tree based only on homozygous SNPs while discarding positions that have some heterozygous SNP in any isolate, as done previously for other *Candida* species[4].

To select the best tree out of the six we assessed various quality metrics (**Extended Data Fig. 2a**), and compared their topologies (**Extended Data Fig. 2b**), which revealed various relevant trends. Given that we defined the *C. parapsilosis* outgroup clade based on a prior tree reconstruction with *C. orthopsilosis* homozygous strains set as outgroups (see **Methods**), we evaluated whether the final *C. parapsilosis*-only unrooted trees (resampled or not) had the expected outgroup. This was the case for all of them, as the metric fraction_trees_expected_outgroup is always 1.0 (**Extended Data Fig. 2a**), validating our strategy for defining the *C. parapsilosis* outgroup. Additionally, despite all trees having many (39-68%) nodes with low (< 90) branch support (see pct_collapsed_nodes, **Methods** and **Extended Data Fig. 2a**), this likely results from the inclusion of highly similar, clonal strains, which cannot be fully resolved in the tree. This is consistent with lowly supported nodes generally having very short branch lengths, with pct_collapsed_long_nodes always < 1.1% (see **Methods**), and/or few diverse child nodes, with pct_politomies_diverse always < 4.1% (see **Methods**) (**Extended Data Fig. 2a**).

Most importantly, various observations suggest that the diploid mode greatly outperforms the diploid_homozygous one. For instance, the diploid mode yields significantly higher numbers of parsimony informative sites for tree reconstruction (>=15,477) as compared to the diploid_homozygous mode (<=9,696), suggesting that it is more robust (**Extended Data Fig. 2a**), a large difference that can likely be explained by the removal of positions with heterozygous SNPs in diploid_homozygous mode. Accordingly, the mode largely impacts the topology itself, as trees generated by the same aligner and different modes have < 58% of equal nodes with support >= 90 (**Extended Data Fig. 2b**). Furthermore, we find that the trees generated in diploid mode with data from different aligners are more consistent (>97% identical nodes with support >= 90) as compared to those generated from different aligners and the diploid_homozygous mode (<72% equal nodes with support >= 90) (**Extended Data Fig. 2b**).

These findings show that even in a highly homozygous species like *C. parapsilosis* the haplotype sampling approach (diploid mode) may be more suited, so we used it in all our tree reconstruction pipelines. Beyond this application to our data, our findings are relevant for further studies of phylogenetic reconstruction in diploid individuals, showcasing the importance of considering heterozygous SNPs as proposed before[3]. Conversely, we could not find clear differences between the results of different aligners, so we used bwa mem-called variants to derive the tree including all isolates (**Fig. 1b**), as it is commonly used[1,2,5].

#### Phylogeographic context of our collection

To understand the phylogeographic context of the isolates analyzed here we assessed clade distribution across countries and Spanish hospitals (**Fig. 1c,d**). While the five major clades showed no clear country bias, some sub-clades displayed unbalanced distributions. For instance, sub-clade 1.1 was exclusive to Turkey, while most of the newly described Spanish sub-clades (4.1, 4.2, 4.3) were geographically restricted, with 4.1 found only in Barcelona and 4.3 predominantly in Sevilla. Also most Santander isolates belonged to clade 4. These findings suggest both global and local propagation among Spanish clinical isolates.

#### Extensive phenotyping of our collection

To study mechanisms of resistance, we retrieved various drug susceptibility phenotypes for the 189 newly sequenced isolates (**Fig. 2**, **Extended Data Figs. 4,5**). These include susceptibility in liquid media towards three echinocandins (caspofungin, micafungin, anidulafungin), 5-flucytosine (5-FC), the polyene amphotericin B and five azoles (fluconazole, voriconazole, itraconazole, posaconazole and isavuconazole), measured in [6]. Based on the distributions of minimum inhibitory concentrations (MIC) and clinical resistance breakpoints we categorized strains as resistant or susceptible (**Extended Data Fig. 4 a**). For echinocandins and amphotericin B, no isolates exhibited clinical resistance over the EUCAST defined breakpoint; therefore, we defined as “resistant” those isolates displaying reduced susceptibility within the clinically susceptible range. Additionally, we measured fluconazole susceptibility using E-test (see **Methods**) in 61 isolates, which allowed us to identify three relevant phenotypes, related to resistance in solid media, that are variable across isolates: heteroresistance, higher resistance in solid vs liquid media, and tolerance at 48h (**Extended Data Fig. 4b,c,d**).

To extend phenotypic insights to virulence-related and other clinically-relevant phenotypes we considered phenotype data for various additional traits, measured previously for our strains[7]. This included growth viability in six disinfectants (ethanol, hypochlorite, $H_2O_2$, chlorhexidine, and the detergents zwittergent 3-14 and surfanio) for 32 outbreak isolates, identifying strains with particularly high and low tolerance (**Extended Data Fig. 4e**). Also, for 77 isolates we measured their capacity for agar invasion, pseudohyphal growth, or biofilm formation (**Fig. 2**). Finally, for ten isolates we report variable behaviors in a microfluidics catheter-like device (i.e. adherence, formation and size of cell clumps) (**Fig. 2**).

To elucidate whether genetic variants modulate clinical pathogenesis, we considered as relevant 'traits' of an isolate the clinical information related to the infected patient. We defined patient age groups ("infant", "young" and "old") based on age distribution (**Extended Data Fig. 5a**), and considered isolation source (e.g. blood, environmental or respiratory) (**Extended Data Fig. 5b**). While these "traits" may not solely reflect strain genetics (patient status and clinical context can be very significant), defining them as binary phenotypes would enable us to find variants predisposing strains to infect specific patient populations (e.g. infants) or body sites (e.g. blood). All in all, this comprehensive collection of phenotypes from a single outbreak offers an unprecedented capacity to study genotype-phenotype relationships in *C. parapsilosis*. Furthermore, given that most similar studies in *Candida* focused only on drug resistance traits, our inclusion of virulence and clinical phenotypes makes our dataset particularly unique.

#### Choosing parameters for GWAS filtering

##### Defining high-confidence GWAS hits

To define adequate filtering parameters for GWAS hits we explored changing the i) aligner (bwa mem, bowtie2 or hisat2), ii) minimum branch support (10, 50, 70, 90), iii) ASR method (DOWNPASS, MPPA or MPPA,DOWNPASS consensus), iv) sample set (all, representative or balanced sets), v) minimum convergence level ($\varepsilon$) (0.1, 0.2 or 0.3), vi) minimum number of nodes with genotype and phenotype transitions ($n_{G\&P}$) (2 or 3) and vii) distinct combinations of p-values and corrections considered. Specifically, we considered using i) no p values, ii) uncorrected $p(X^2)$ or $p(n_{G\&P})$, iii)

FDR-corrected $p(X^2)$, $p(n_{G\&P})$ or both, iii) bonferroni-corrected $p(X^2)$, $p(n_{G\&P})$ or both and iv) $p(X^2)_{maxT}$, $p(\varepsilon)_{maxT}$ or both. Also, when p value correction was used (including when $p(X^2)_{maxT}$ or $p(\varepsilon)_{maxT}$ were used), we considered the effect of applying the correction across i) all of the tested groups (for a given combination of aligner, ASR method, min support, and sample set) (conservative 'correction scope') vs ii) all groups with the same variant grouping strategy (certain values for 'type_vars', 'type_mutations' and 'type_collapsing') (relaxed 'correction scope'). Furthermore, although we used a $p<0.05$ in most cases, we explored the effect of various thresholds (i.e. $p<10^{-4}$, $p<10^{-3}$ or $p<0.05$) when using uncorrected p values. Thus, for each phenotype we tested 18,144 parameter combinations, expanding those tested in [2].

To identify optimal parameter combinations we evaluated the effect of these filters on three relevant features. First, as done in [2], we assessed the amount of protein-altering hits in expected genes for resistance phenotypes (as defined in 'Assessing genes previously related to drug resistance'). We could find such instances for resistance towards fluconazole (with hits on *ERG11, MRR1* and *NDT80*), voriconazole (*ERG11, MRR1* and *NDT80*), posaconazole (*ERG11* and *TAC1*) and caspofungin (*FKS2*). Second, we scored the fraction of phenotype transitions that can be attributed to some protein-altering hit, which is a rough proxy for the recall of our filtering strategy. Third, we calculated the number of variants with hits that do not result from convergence across clades, which may be false positives (justified in the next paragraph). Ideally, we wanted to find filters that i) would yield most of the expected genes, ii) had hits explaining most of the phenotype transitions in a large fraction of phenotypes and iii) did not yield many 'false positive' variant hits (e.g. <10).

Various lines of evidence support the assumption that variant hits not found convergently across many clades are likely false positives. Variants appearing as GWAS hits may be attributed to either i) recombination between genetically-distinct isolates leading to phenotypic changes, ii) independent acquisition of the same variant by various *de novo* phenotype transition events or iii) spurious associations related to multiple testing and/or noisy ASR results (e.g. wrong ASR methods or poor tree resolution) leading to false positive hits. Given the mostly asexual diversification of *C. parapsilosis* (**Extended Data Fig. 3**), we expected recombination to be an infrequent driver of phenotypic change. Furthermore, in the absence of recombination, we expected genotype-phenotype convergence to be mostly detectable at the level of genes / domains / pathways rather at the variant level, as illustrated by the finding that many different mutations in similar genes lead to resistance in *Candida*[2,8,9]. Hence, we expected most variant hits to be false positives, unless there was clear evidence for independent variant acquisitions across multiple clades. To illustrate this, we show the variant hits obtained when using very relaxed filters (i.e. using min_support = 10 and no multiple testing correction) (**Extended Data Fig. 9**).

To perform the filter benchmarking while validating that the number of variants with hits is an indicator of false positive rate we visualized the effect of applying different filters in a sequential way. Initially, we visualized the effect of varying the p value types, which revealed relevant patterns (**Extended Data Fig. 8a**). First, *TAC1*-related hits can only be found in posaconazole, but only by a few filters that use no p value corrections, so we discarded it for further filter benchmarking. Second, we generally see the expected genes in the expected phenotypes, with the exception of a few filters (i.e. not using p values) yielding *ERG11* hits in caspofungin. This validates our approach of using expected genes, and shows the importance of considering p values to avoid false positives. Third, parameter sets using no or less conservative p values (e.g. with no or FDR correction) tend to yield higher numbers of variants with hits, supporting the idea that high numbers of such hits is an indicator of false positives. Fourth, we find strategies based on p value correction yielding most of the expected genes (except *TAC1* in posaconazole), suggesting that we should be using some correction. Fifth, there are no p value strategies that always yield optimal parameters (i.e. yielding all expected genes and low numbers of variant hits), showing the need for additional parameters.

In a second step, we visualized the effect of varying the min support on filters using some sort of p value correction (**Extended Data Fig. 8b**), now looking only at *ERG11, MRR1*, *NDT80* and *FKS2*. We found a clear correlation between a lower support and very high number of variants with hits, further suggesting these are most likely false positives, and supporting the use of higher support filters (70 or 90). Also, our analysis shows that a min support of 10 (very relaxed) is required to find *ERG11* or *MRR1* in voriconazole, so we have to accept missing these genes for getting high-confidence results. Finally, even within filters that use a min support of 70 or 90 we still find a large variability in the number of (likely false positive) variant hits, suggesting that more parameters should be explored. Thus, we kept parameters with a min support of 70 or 90 for further benchmarking.

Next, we checked the impact of varying the ASR method (**Extended Data Fig. 8c**), which had a phenotype-dependent effect. We need to use MPPA to find *FKS2* in caspofungin, but this comes at the cost of i) mostly having many variant hits and ii) missing *NDT80* in fluconazole / voriconazole and *ERG11* in posaconazole. Conversely, various MPPA, DOWNPASS consensus or DOWNPASS-based parameters result in good outcomes (even at the cost of missing *FKS2* in caspofungin), so we further explored parameters related to these methods. Also, we checked the impact of the min ε (**Extended**

**Data Fig. 8d**). This showed that all values for this parameter had redundant effects, so we kept the most conservative min $\varepsilon$=0.3.

Furthermore, we checked the combined impact of varying the sample set and the min $n_{G\&P}$ (**Extended Data Fig. 8e,f**). This suggested that using balanced or representative samples generally yielded better results than using all samples. The choice between representative and balanced samples was not trivial, as the former would miss *ERG11* in fluconazole, while the latter would miss *MRR1* in the same drug. However, parameters yielding *MRR1* with representative samples needed min $n_{G\&P}$ = 3, which is likely too conservative (i.e. missing *NDT80* in voriconazole and yielding 0 hits in caspofungin). Thus, even at the cost of missing *MRR1* in fluconazole, we used balanced samples and $n_{G\&P}$ = 2. Next, we evaluated the impact of the aligner (**Extended Data Fig. 8g**), which showed that bwa mem-based parameters perform worst (i.e. missing most expected genes). Also, we had to use hisat2 to find *ERG11* in fluconazole, while bowtie2 was necessary to find *NDT80* in voriconazole (but this was at the cost of having many variant hits). Thus, we chose hisat2 for the final parameters.

This hierarchical parameter selection allowed us to narrow down to 72 parameter combinations. To further select optimal filters we chose those that got all of the (remaining) expected genes (i.e. *ERG11* for fluconazole / posaconazole and *NDT80* for fluconazole), which are three combinations (**Supplementary Table 2**) using $p(n_{G\&P})$<0.05 that are either i) FDR-corrected with a conservative scope, ii) FDR-corrected with a relaxed scope or iii) bonferroni-corrected with a relaxed scope. Given that FDR correction may yield unexpected behaviors on highly interdependent tests[10], which is the case for our GWAS, we kept the filters using bonferroni correction. Thus, the final parameters used min_support = 70, aligner = hisat2, ASR method = MPPA,DOWNPASS, sample_set = balanced_samples, min $\varepsilon$ = 0.3, min $n_{G\&P}$ = 2, (only) bonferroni-corrected $p(n_{G\&P})$ < 0.05 and a relaxed correction scope.

##### Defining low-confidence GWAS hits

Given the tradeoffs that had to be made to achieve high-confidence hits across phenotypes (e.g. missing certain expected genes) we also designed a filtering strategy that would yield a wider range of hits, which constitute our 'low confidence' set of hits. To find these we focused on parameter combinations yielding all expected genes in all phenotypes (**Extended Data Fig. 8a**). We could not find such filters, but when discarding *TAC1* from consideration we could find 44 adequate parameter combinations. These yielded *MRR1* / *NDT80* / *ERG11* for fluconazole and voriconazole, *ERG11* for posaconazole and *FKS2* for caspofungin, as expected (**Extended Data Fig. 8a**). A closer look at these filters (**Supplementary Table 3**) revealed that all of them were based on min support = 10 and yielded >=95 (likely false positive) variant hits in some phenotype, indicating that they indeed yield various spurious results.

However, to still get a set of low-confidence hits we picked parameters yielding a number of variant hits close to the minimum possible (<110), while using uncorrected p values and a 0.05 threshold. We found three such parameters , yielding <= 99 variant hits and using various combinations of uncorrected $p(n_{G\&P})$ and/or $p(X^2)$. For consistency with the filters used for high-confidence filtering we used the ones based only on $p(n_{G\&P})$. These use min_support = 10, aligner = hisat2, ASR method = MPPA, sample_set = all samples, min $\varepsilon$ = 0.2, min $n_{G\&P}$ = 2, (only) uncorrected $p(n_{G\&P}) < 0.05$.

#### Parasexual recombination may underlie certain genotype-phenotype associations

To assess genetic architecture we inspected the type of the GWAS high-confidence hits obtained **(Fig. 4b)**. While most hits are groups of collapsed variants, suggesting independent *de novo* acquisition of such traits, for four phenotypes we could find >=5 hits on single variants **(Fig. 4b,c)**. This was unexpected given the mostly clonal nature of this species (**Extended Data Fig. 3**), where independent phenotypic transitions are unlikely to result from the same variant. Although these hits could include false positives (**Extended Data Fig. 9**), our conservative filtering suggests otherwise. Careful inspection of these cases revealed different scenarios. For 'voriconazole resistance' and 'source sterile', the variants associated to the phenotype were generally unlinked from each other (**Extended Data Fig. 10**), suggesting convergence through recombination-independent mechanisms. However, for 'chlorhexidine tolerance' and 'fluconazole resistance' tracts of 5 or more linked variants, often on different scaffolds, were associated with some phenotypic transitions (**Extended Data Fig. 11**). This unexpected result suggests an occasional role of (para)sexual recombination in the spread of these traits, as observed in other *Candida* species[2].

#### Challenges and opportunities for building ML classifiers in *Candida parapsilosis*

We anticipated the building of ML classifiers to be a challenging task, potentially leading to either inaccurate classification or model overfitting due to various reasons. First, our dataset has limited size (<200 isolates) but thousands of variants that may be used as predictive features, often correlated with each other. Second, the genetic diversification of the analyzed collection is most likely the result of clonal reproduction (**Extended Data Fig. 3**), suggesting that the independent acquisition of each phenotype (**Fig. 2**) does not necessarily result from resistance variants having spread sexually, but rather from independent mutations. Consequently, standard prediction from the presence/absence patterns of variants (as done in [11–14]) may not be fruitful. Third, the correlation between phenotype and phylogeny (**Fig. 2**) may complicate the analysis, as ML models may only learn the underlying phylogenetic association, and not the causal variants of each phenotype, leading to overfitting (as we propose happened in [13,14]). Fourth, some clades are likely overrepresented in our dataset (e.g. clades with azole resistance, **Fig. 2**), generating sample redundancy that may lead to models biased towards such clades. Fifth, our limited sample size means that a train/test splitting may leave insufficient samples for performance estimation.

At the same time, various reasons suggest that we can elucidate genotype-phenotype relationships through ML classifiers. First, several phenotypes (e.g. resistance) are expected to have a simple genetic architecture, where a few variants with high effect may underlie the phenotype[8,15,16]. Second, at least the experimentally-measured traits are likely to be genetically determined to a large degree (e.g. similarly to rare human diseases where ML classifiers have been successfully used[17]), fulfilling the assumptions of ML approaches. Third, the asexual nature of *C. parapsilosis* implies that we can precisely model population structure through a phylogeny (i.e. reticular evolution is not expected) (**Fig. 1**), which enables the use of phylogeny-aware ML (see **Main text**). Fourth, previous efforts could predict with high accuracy micafungin heteroresistance in *C. parapsilosis*, using a comparable sample size (n=219)[14], suggesting we may have sufficient power to build useful ML classifiers.

#### Optimization of ML classifier parameters inform about best practices for building predictive models in *Candida*

To understand parameter choice and learn how to build ML classifiers in *Candida* we initially examined the used 'test parameters' among the top-performing models (i.e. those yielded by the 'top parameters' in **Fig. 5c**), which provided interesting insights about parameter optimality (**Extended Data Fig. 19,20**). First, the aligner used slightly influences model performance, so that hisat2 and bowtie2 generally yield better results than bwa mem (**Extended Data Fig. 20**). Second, using some measure of feature consistency for training / evaluation seems necessary for the azoles, but not for the other phenotypes (**Extended Data Fig. 20**), which may reflect differences in genetic architectures between the phenotypes. Third, setting different values for the 'AUC diff' parameter (indicating how much we prioritize ROC AUC over feature consistency when selecting the optimal model during training / evaluation) had a very minor effect (**Extended Data Fig. 20**). Fourth, narrowing the feature selection of the model on a subset of genes seems to be essential for optimal performance in all phenotypes (**Extended Data Fig. 20**). Also, for azoles we find that using only the previously-expected genes for azole resistance ('known' genes *ERG11*, *ERG3*, *TAC1, MRR1, UPC2* and *NDT80*) yields poor performance for resistance phenotypes, supporting the idea that these genes are insufficient to explain such phenotypic variation.

Furthermore, the combined exploration of train and test parameters (**Extended Data Fig. 13,19,20**) provided insights into how relevant is to consider the phylogenetic structure of the dataset (i.e. either using only representative samples or splitting the data in a phylogenetically-balanced way (**Extended Data Fig. 6b**)). Such parameters only yield clearly better results for the phenotype 'blood vs other sterile' (**Extended Data Fig. 20**). We speculate that this is generated by the particular nature of class imbalance of each phenotype (**Fig. 2**). For fluconazole and voriconazole resistance we have a quite balanced dataset, which likely enables using all isolates. Furthermore, there are resistant / susceptible isolates across most clades, so that a random splitting is less likely to yield overfit results. The remaining two phenotypes ('age infant' and 'blood vs other sterile') have imbalanced classes, so that using all isolates may theoretically yield worse results. This is the case for 'blood vs other sterile', where we need to use representative samples to achieve high performance (**Extended Data Fig. 20**). However, for 'age infant' there is no clear benefit of using these. To better understand this difference we explored the behavior of the 'class weight' parameter, also evaluated on these phenotypes to address the class imbalance issue. While none of the top models used class weighing for 'blood vs other sterile', most of them (20/30) did for 'age infant' (**Extended Data Fig. 20**). This suggests that class weighing compensates for the imbalance in this phenotype, so that there is no clear benefit of using representative samples. This may be attributable to a higher number of representative samples in 'blood vs other sterile' (**Extended Data Fig. 18**) as compared to 'age infant' (**Extended Data Fig. 17**), which influences the viability of modelling with only such subsets of isolates. On another line, the similar behavior for random / balanced splitting for these two phenotypes (**Extended Data Fig. 20**) may be attributable to the fact that the minority class phenotype is not correlated to the phylogeny (**Fig. 2**), so that a random splitting is less likely to yield overfit results.

Finally, the train parameters used by top models offer interesting insights into how these models work (**Extended Data Fig. 13**). First, prioritizing models that include (among others) expected genes (*ERG11, TAC1, MRR1* and *NDT80*) is always the chosen strategy for azole phenotypes, showing the importance of using *a priori* knowledge. Second, the most used AUC threshold for feature selection is the minimum tried (0.05), showing that using features with marginal improvements on training/evaluation performance is an overly good strategy. Third, we observe all ML methods (e.g. random forest, logistic regression …) being used in some top model, without a clear preference among them. This suggests that most modelling strategies work well when provided with adequate features. Fourth, the modelling is based on predicting phenotype transitions for most phenotypes (all except 'age infant'), showing the benefit of our novel approach. Fifth, the transitions used are generally considering pairs of isolates spanning the whole tree (definition=each clade, SNP/kb threshold=1.0), which may be enabled by the low intraspecific diversity of *C. parapsilosis*. Sixth, most top models use all types of predicted transitions for modelling the phenotype (see **Methods**), showing that both transition and no-transition events are informative. Seventh, a low ML threshold (0.25) is generally applied to model phenotypes from transition probabilities (see **Methods**), indicating that a relaxed categorization of transitions is optimal.

These findings improve the explainability of our models, showcase the importance of parameter exploration, and provide an interesting benchmark for further studies.

#### Using ML classifiers to elucidate mechanisms of virulence and drug resistance

To infer evolutionary mechanisms of phenotypic transitions from our ML models, we analyzed the genes contributing to the predictive features of the 'top' m1-m5 models yielded by the 'top parameters' (**Fig. 5a,b, Extended Data Fig. 13**). Given that a top parameter may include several m1-m5 top models (see **Methods, Fig. 5a,b**), this included 22 'top' models for fluconazole resistance, 3 for voriconazole resistance, 30 for 'age infant', and 26 for 'blood vs other sterile'. We identified 16 genes contributing to more than three top models, each mostly associated to one phenotype, with the exception of *ALS11* (related to 'age infant' and 'blood vs other sterile') and *ERG11* (in fluconazole and voriconazole resistance) (**Fig. 6a**). Such reduced numbers of genes suggest an oligogenic basis for such phenotypes.

For fluconazole resistance, all 22 top models used *ERG11* which was combined with one to three other genes, depending on the model. Seven genes were used in three or more top models (*ERG11*, *Scer_MOH1* (i.e. ortholog of *Saccharomyces cerevisiae MOH1*), *CDR1B, Scer_ATP10, MRR1, Scer_ERG24* and *CPAR2_204120*) (**Fig. 6a, Extended Data Fig. 13**). Visualization of the presence/absence pattern of the mutations used by the models along the strains tree (**Fig. 6b,c**) indicated that these genes collectively explain all fluconazole resistance transitions. Most notably, we observe that variants in *CDR1B*, *Scer_MOH1* and *Scer_ERG24,* not typically associated to resistance in this species, are necessary to explain transitions that were not explained by changes in previously known mechanisms involving *ERG11*, *TAC1*, *MRR1* or *NDT80* (**Fig. 3, Fig. 6c**). This highlights our model's utility in leading to novel results. Furthermore, our findings support the previously proposed relevance of *CDR1B* in azole resistance[18].

The fact that different models for fluconazole resistance use distinct sets of genes (**Extended Data Fig. 13**) may suggest redundant predictive patterns (i.e. variants in different genes having correlated presence/absence patterns). However, while this is true to some extent, we also find that independent acquisitions of resistance are often explained by different genes (**Fig. 6c**), suggesting that several of the models may capture complementary mechanisms, rather than redundant ones. This is consistent with each individual model not having perfect predictive performance (AUC ranging 0.80-0.95), and the need to collectively consider all genes to explain all observed resistance transitions (**Fig. 6c**). For instance, the best top model for fluconazole resistance (**Extended Data Fig. 14**,**15**) is useful to explain transitions related to *ERG11*, *MRR1* and *Scer_ERG24*, but misses transitions that can be attributed to changes in *Scer_MOH1* and *CDR1B*. Hence, looking at all top models collectively, as done here, enhances interpretability.

Model interpretability for voriconazole resistance is more straightforward, with all three top models using the genes *STP4, ERG11* and *Scer_CLU1* (**Extended Data Fig. 13**). Therefore, we inspected how the features of the best top model (AUC=0.73) explained the presence / absence patterns of the phenotype (**Extended Data Fig. 16**). While most transitions can be explained by variants in *ERG11* or *Scer_CLU1,* there are four that remain unexplained by this model or by changes in *TAC1*, *MRR1* or *NDT80* (**Fig. 3, Extended Data Fig. 16**), suggesting there are additional drivers of voriconazole resistance.

Similarly, for 'age infant' we could also interpret the models based on the performance of two top models. All 30 top models used *Scer_PMI40* in combination with either *ALS11* (20/30) or *RBT1* (10/30). Among the best top test parameters we had two models using each of these combinations (**Extended Data Fig. 13**), respectively, so we used them as representatives (*RBT1* model AUC=0.77; *ALS11* model AUC=0.79). Based on their variant patterns (**Extended Data Fig. 17**), we infer that the absence of variants in *Scer_PMI40* and *ALS11/RBT1* (correlated to each other) increases the predisposition towards infecting infants. The high false positive rate of these models (**Extended Data Fig. 17**) might be due to unknown drivers or the possibility that strains with high potential to infect infants are additionally infecting other age groups.

Finally, for 'blood vs other sterile' all of the 26 top models used *Scer_MDS3* combined with one (24/26) or two (2/26) additional genes (**Extended Data Fig. 13**). Five genes appeared in at least three models (*Scer_MDS3, ALS11*, *ALS7*, *BCR1* and *Scer_DFG16*), so we focused on models considering these. In contrast to fluconazole resistance (discussed above), where many genes were complementary, for 'blood vs other sterile' we could find a set of 'best top parameters' yielding 5 highly accurate models (AUC from 0.93-0.97), of which four used *Scer_MDS3, ALS11* and *ALS7*, and not *BCR1* or *Scer_DFG16* (**Extended Data Fig. 13**). This suggests that the former three genes capture the relevant genotype-phenotype signal, whereas the other genes are redundant and not relevant to the phenotype. We inspected the variant patterns for two representative models, each using *Scer_MDS3* and either *ALS11* or *ALS7* (**Extended Data Fig. 18**), and concluded that several regulatory variants in *Scer_MDS3* increase the predisposition towards infecting non-sterile compartments outside the bloodstream, while the effect of *ALS11*/*ALS7* is more complex to interpret.

In summary, our analysis of predictive features enables a deep understanding of the classifier behavior, and suggests novel determinants of resistance and virulence that deserve further attention.

#### Comprehensive integration of all genotype-phenotype relationships characterized in this study

The analysis of expected genes, convergence GWAS and ML suggested 79 prioritized gene-phenotype mappings (50 genes and 15 phenotypes), including the most likely relevant genes for each phenotype (see **Main Text, Methods, Fig. 7, Supplementary Table 6**). In brief, these prioritized mappings include genes i) known to be associated with the phenotype (**Fig. 3**), ii) related to ML features, iii) associated to GWAS hits in other *Candida* species[2] for similar phenotypes, iv) with relevant functional descriptions in Candida Genome Database (CGD), v) under genomic selection in other *Candida* species[2] or vi) among the top three genes with strongest genotype-phenotype mappings. Also, to explore the similarities among ergosterol-related phenotypes (fluconazole R, voriconazole R, posaconazole R and amphotericin B R), we added genes i) with GWAS hits (high or low confidence) in the ergosterol-related phenotype of interest but not prioritized by the former six criteria and ii) prioritized in some ergosterol-related phenotype.

To interpret and validate these results we checked, for each gene, i) their functional annotations (**Supplementary Table 6**), ii) the Candida Genome Database (CGD)[19] 'phenotype' information for the

*C. parapsilosis* gene and *C. albicans* orthologs, iii) the CGD *C. albicans* orthologs' functional annotations, iv) the Saccharomyces Genome Database (SGD)[20] 'phenotype' information and v) existing literature about the genes or their orthologs in fungi. This curation for all 79 prioritized gene-phenotype mappings revealed various relevant trends, discussed below. Note that we refer to the genes by either their *C. parapsilosis* gene name if existing (e.g. *ERG11*), the name of the *Saccharomyces cerevisiae* ortholog if existing (e.g. *Scer_NCP1*), or the gene symbol if no gene name or *S. cerevisiae* orthologs exist (e.g. *CPAR2_806400*).

#### Azole resistance general insights

Beyond the known genes *ERG11*, *MRR1, TAC1* and *NDT80*, we identify 18 additional ones related to azole resistance, which shows the power of our approach for suggesting novel candidates even for such well-studied phenotypes (**Supplementary Table 6**). While 7/22 such genes can be related to pan-azole resistance (towards fluconazole, voriconazole and posaconazole), the others are associated to only fluconazole and voriconazole resistance (7/22), or to only one azole (8/22), suggesting that resistance mechanisms differ across azole drugs, as previously proposed in other *Candida* species[2,8,21]. However, fluconazole and voriconazole-resistance genes are more similar to each other than posaconazole-resistance genes, which may be explained due to the fact that the former two are short-tailed, while the latter is a long-tailed azole[22]. In the next sections we report the relevant evidence for each of these genes.

#### Pan-azole resistance genes

Regarding azole resistance traits, we find seven genes related to pan-azole resistance (at least voriconazole, fluconazole and posaconazole): *ERG11, NDT80, TAC1, MRR1, Scer_NCP1, Scer_TEL1* and *CPAR2_806400* (**Supplementary Table 6**). *ERG11* is the azole target, and mutations in this gene have been commonly associated with azole resistance, likely due to reduced drug binding due to missense mutations[23] or drug titration due to *ERG11* overexpression related to gene duplications[18]. Similarly, *NDT80, TAC1* and *MRR1* are regulators of azole efflux previously related to azole resistance[23]. Thus, the associations found on these four genes further confirm the importance of these genes. Conversely, the findings regarding *Scer_NCP1, Scer_TEL1* and *CPAR2_806400* are more novel. *Scer_NCP1* is a predicted NADPH-hemoprotein reductase, with a putative role in ergosterol biosynthetic process, and interacting with *ERG11* in *C. albicans* to enable azole tolerance[24]. Given that we find *Scer_NCP1* duplications associated with resistance, we propose that such variants lead to the hyperactivation of this tolerance mechanism, generating azole resistance.

Furthermore, *Scer_TEL1* has annotations of protein serine/threonine kinase activity, telomeric DNA binding and DNA damage responses, with signs of genomic selection in other *Candida* species[2]. Also, *TEL1* has been shown to be overexpressed in *Aspergillus fumigatus* biofilms during itraconazole exposure[25], so that its association to azole resistance may be related to biofilm formation and DNA damage stress responses. Finally, *CPAR2_806400* has no annotations nor *C. albicans* / *S. cerevisiae* orthologs (CGD), but it was reported as a cell wall / cell membrane protein acquiring mutations during echinocandin adaptation in *C. parapsilosis*[26]. Thus, the association of this gene with pan-azole resistance described here suggests that it may be an uncharacterized, but relevant mediator of multidrug-resistance (MDR) towards different drug families.

In summary, we find pan-azole resistance genes possibly related to impaired drug binding and/or titration (*ERG11*), regulation of azole efflux (*NDT80, TAC1* and *MRR1*), ergosterol biosynthesis (*Scer_NCP1*), biofilm formation and DNA-damage stress responses (*Scer_TEL1*). Also, we identify a novel gene, *CPAR2_806400*, with unknown function.

##### Fluconazole and voriconazole resistance genes

We find seven novel genes related to voriconazole and fluconazole resistance: *CDR1B, Scer_UTP22, Scer_UGA2*, *CPAR2_105460, CPAR2_105480, CPAR2_204120* and *CPAR2_204210* (**Supplementary Table 6**). *CDR1B* encodes an azole efflux pump, regulated by *MRR1*, so we may be detecting hyper-activating variants leading to resistance due to increased efflux via this pump. This is consistent with a previous study proposing a link between *CDR1B* duplications and fluconazole resistance in *C. parapsilosis*[18]. However, we find different types of mutations associated with each of the azoles, showing distinct mechanisms of resistance. We find *CDR1B* missense variants associated with fluconazole resistance, which may hyperactivate drug efflux leading to the phenotype. Conversely, we find a deletion overlapping the 5' CDS region of *CDR1B* associated with voriconazole resistance, which was puzzling due to the expectation of hyperactivating mutations leading to azole resistance. We propose that this variant actually reflects a gene conversion event, which is interpreted as a deletion due to assembly errors in the used CDC317 reference genome. A recent study[18] showed that CDC317 has two tandem *CDR1B* genes (*CDR1B.1, CDR1B.2*), but the reference genome has an erroneous fusion gene including the 5' region of *CDR1B.2* and the 3' region of *CDR1B.1*. Thus we interpret our findings as a gene conversion event where the *CDR1B.1* copy is kept, suggesting that this copy encodes a more efficient efflux pump. These findings further validate the previously proposed[18] importance of *CDR1B* variants in azole resistance.

Furthermore, *Scer_UTP22* has a predicted role in rRNA processing and tRNA export, with signs of genomic selection in other *Candida* species[2]. *UTP22* has been linked to amphotericin B adaptation in *C. auris*[27], and it is regulated by the azole-response transcription factor *NCB2* in *C. albicans* azole-resistant strains[28]. These findings suggest that the *NCB2*-regulated azole response[28] may also exist in *C. parapsilosis*, and that *UTP22* could be a relevant downstream effector that can be modulated by mutations leading to resistance. Similarly, *Scer_UGA2* is a putative succinate-semialdehyde dehydrogenase, overexpressed in fluconazole and voriconazole-resistant strains. Specifically, *MRR1*-mutated fluconazole-resistant *C. parapsilosis* strains have increased *Scer_UGA2* expression[29]. Together with our GWAS results on this gene, these findings suggest that the *MRR1*-regulated azole response may involve not only efflux pumps (*MDR1B* and *CDR1B*, see [29]), but also metabolic genes such as *Scer_UGA2*, which can also be hyperactivated by mutations leading to resistance. Also, null *UGA2* mutants in *S. cerevisiae* have increased miconazole resistance (SGD), further supporting the link of this gene with azole responses.

*CPAR2_105460* has aspartate and asparagine metabolism annotations, but it lacks *C. albicans* / *S. cerevisiae* orthologs (CGD) that would enable us to infer precise biological functions. However, this gene has orthologs in other *Candida* species associated with azole resistance in a recent GWAS[2], supporting the importance of this uncharacterized gene in azole resistance. In addition, *CPAR2_105480* has many inconclusive metabolism-related annotations (e.g. GDP-L-fucose biosynthesis, glycerol degradation, fatty acid beta-oxidation), but it has been found to be overexpressed in fluconazole and voriconazole resistant strains, which supports its link to such drugs. Furthermore, *CPAR2_204120* has a predicted role in protein glycosylation, but we could not find additional evidence explaining its relationship with azole resistance. Finally, *CPAR2_204210* has domains with predicted electron transfer activity, heme binding, iron ion binding, monooxygenase activity, oxidoreductase activity and acting on paired donors, and has annotations of an involvement in phenylacetate and xenobiotic metabolism. According to a recent study[30], this gene is a homolog of the *C. albicans* alkane-inducible cytochrome P450 gene *ALK8*, whose overexpression has been linked to fluconazole resistance (CGD), which validates the observed associations.

All in all, the genes associated to only fluconazole and voriconazole resistance are possibly related to drug efflux (*CDR1B*), rRNA processing and tRNA export (*Scer_UTP22*), aminoacid, energy and xenobiotic metabolism (*Scer_UGA2, CPAR2_105460, CPAR2_105480* and *CPAR2_204210*), and protein glycosilation (*CPAR2_204120*).

**Fluconazole, voriconazole and posaconazole-only resistance genes**

We find four genes associated only with fluconazole resistance: *Scer_ATP10, Scer_MOH1, Scer_ERG24* and *ERG25* (**Supplementary Table 6**). *Scer_ATP10* has a predicted role in mitochondrial proton-transporting ATP synthase complex assembly, and null mutants of this gene in *S. cerevisiae* have upregulated drug efflux transporters (SGD). Given the link between mitochondrial dysfunction and upregulation of azole efflux pumps[31], we propose that *Scer_ATP10* variants lead to fluconazole resistance via a hyperactivation of this physiological tolerance mechanism. Furthermore, *Scer_MOH1* has orthologs in *S. cerevisiae* with predicted activity in vacuolar transport (SGD), and mutations in this gene have been linked to caspofungin adaptation in *C. glabrata*[32]. Thus, while the specific mechanisms linking this gene to resistance remain elusive, the findings on *C. glabrata* show that it may be related to adaptation to different drug families. In addition, *Scer_ERG24* has a predicted role in ergosterol biosynthesis, so its link to resistance may be related to altered sterol composition due to mutations. Also, various observations validate its link to azole resistance, including i) its overexpression in *C. parapsilosis* azole-resistant strains with *ERG3* mutations[33], and ii) *C. albicans* null mutants generating increased azole resistance[34]. Finally, *ERG25* also has a role in ergosterol biosynthesis in the endoplasmic reticulum, so that its mutations may also change sterol composition leading to resistance. Accordingly, *ERG25* null mutants in *C. albicans* have increased fluconazole sensitivity[35].

Conversely, we find three genes associated only with voriconazole resistance: *Scer_CLU1, STP4* and *Scer_NUP1*. *Scer_CLU1* is a putative RNA binding protein with a predicted role in the cellular response to osmotic stress and cytoplasmic stress granule localization, but we could not find additional evidence linking it to azole resistance, making it a novel candidate. Conversely, *STP4* encodes a putative zinc finger transcription factor, overexpressed in fluconazole and voriconazole-resistant strains. Also, additional lines of evidence support the link of this gene to azole resistance, including i) its overexpression in fluconazole-resistant strains with *MRR1* mutations in *C. parapsilosis*[29], ii) its expression induced by fluconazole in *C. albicans*[36], iii) the fact that it is a target of *NDT80* (azole azole-related transcription factor) regulation in *C. albicans*[37] and iv) the overexpression of the *S. cerevisiae* ortholog (*STP3*) leading to fluconazole sensitivity (SGD). Specifically, due to the association with azole-response regulators *MRR1* and *NDT80*, we propose that *STP4* may be an additional transcriptional regulator of azole responses, which may be modulated through mutations leading to resistance. Furthermore, *Scer_NUP1* is a putative structural constituent of nuclear pore activity, and it was prioritized due to orthologs in other

*Candida* species being associated with azole resistance in a recent GWAS[2]. Accordingly, its orthologs in *Cryptococcus gattii* are differentially expressed upon fluconazole treatment[38], which supports its (uncharacterized) role in resistance.

Finally, we find one gene associated with only posaconazole resistance: *Scer_NST1*. This gene has a putative role in the response to osmotic stress, and has orthologs in other *Candida* species associated with azole resistance in a recent GWAS[2]. In addition, null mutants in *NST1* lead to thiabendazole tolerance in *S. cerevisiae*[39], which serves as a potential validation of this gene. As for *Scer_CLU1* (discussed above), our results on *Scer_NST1* suggest a link with osmotic stress and azole tolerance / resistance pathways.

In summary, the genes associated to resistance towards only one azole are possibly related to mitochondrial-activated drug efflux (*Scer_ATP10*), vacuolar transport (*Scer_MOH1*), ergosterol biosynthesis (*Scer_ERG24* and *ERG25*), osmotic stress responses (*Scer_CLU1* and *Scer_NST1*), transcriptional regulation of azole responses (*STP4*) and nuclear pore activity (*Scer_NUP1*).

##### Amphotericin B resistance genes

Our prioritization strategy suggested four relevant genes linked to amphotericin B resistance: *Scer_NCP1*, *CPAR2_806400*, *CPAR2_204210* and *Scer_FOL1* (**Supplementary Table 6**). The first three (*Scer_NCP1*, *CPAR2_806400*, *CPAR2_204210*) were also linked to azole resistance (see above), so their association with amphotericin B suggests common resistance mechanisms between these two drug families, perhaps due to common metabolic changes in ergosterol biosynthesis (targeted by azoles and polyenes[23]). Specifically, *Scer_NCP1* has a putative role in ergosterol biosynthesis, and we find that duplications of this gene are associated with amphotericin B resistance. Given that the toxicity of this drug is related to its binding to ergosterol[40], duplications in *Scer_NCP1* may generate resistance due to the (predicted) resulting increase in Ncp1p expression and higher ergosterol production, which may titrate away the drug.

Also, the uncharacterized gene *CPAR2_806400* may be particularly relevant due its association with amphotericin B resistance, pan-azole resistance (see above) and echinocandin adaptation[26]. Thus, we propose that this gene is a mediator of resistance towards all major antifungal families, and could be an interesting target for the development of co-adjuvant therapies. Furthermore, *CPAR2_204210* has a predicted role in xenobiotic metabolism, which may explain its relationship to azole and amphotericin B resistance. Conversely, *Scer_FOL1* has a predicted role in the tetrahydrofolate biosynthetic process. While we could not find specific literature linking this gene to polyene resistance, it has been proposed that *FOL1* could be a good antifungal target[41], and our results suggest such new drugs targeting this enzyme may be good amphotericin B co-adjuvants.

All in all, amphotericin B resistance is related to genes involved in ergosterol biosynthesis (*Scer_NCP1*), xenobiotic metabolism (*CPAR2_204210*) and tetrahydrofolate biosynthesis (*Scer_FOL1*). Also, we identify a novel gene, *CPAR2_806400*, with unknown function. Note that most of them (*Scer_NCP1, CPAR2_806400* and *CPAR2_204210*) were also related to azole resistance, which suggests common resistance mechanisms between the two drug families.

##### 5-flucytosine resistance genes

For 5-FC, our prioritization strategy yielded three genes: *CPAR2_105750, Scer_ENA2* and *CPAR2_806390* (**Supplementary Table 6**). *CPAR2_105750* has domains with predicted DNA binding activity, and annotations of glucose biosynthesis and trehalose degradation. Also, it is upregulated in *C. parapsilosis* azole-resistant strains with *MRR1* mutations[29], validating a potential link of this gene with drug responses. Conversely, *Scer_ENA2* has predicted P-type ion transporter activity for calcium, potassium and sodium. In addition, various lines of evidence support the link of this gene to stress or drug responses, including i) its induction by the antifungal ciclopirox olamine in *C. albicans* (CGD), and ii) its involvement in salt osmotic responses in *S. cerevisiae* (SGD). Finally, *CPAR2_806390* has no annotations or orthologs in *C. albicans* / *S. cerevisiae* that may allow us to understand its function, so it represents a novel, unconfirmed result.

In summary, 5-flucytosine resistance is associated with genes involved in regulation of carbohydrate metabolism (*CPAR2_105750*) and transmembrane ion transport (*Scer_ENA2*). Also, we identify a novel gene, *CPAR2_806390*, with unknown function.

##### Chlorhexidine tolerance genes

For chlorhexidine tolerance we prioritized four genes: *CPAR2_101330, Scer_ORC4, Scer_RAD23* and *Scer_MNR2* (**Supplementary Table 6**). *CPAR2_101330* has domains with a predicted role in response to stress and integral component of membrane localization, so we hypothesize that its role in stress responses may be modulated by mutations that lead to such tolerance. In addition, *Scer_ORC4* has a predicted role in DNA replication and heterochromatin assembly, and has signs of genomic selection in other *Candida* species[2]. Also, null mutants in *S. cerevisiae* change susceptibility towards various chemicals (staurosporine, dichlorophen, dichloroaniline and methylaniline) (SGD), which support the role of this gene in chlorhexidine responses. Furthermore, *Scer_RAD23* has predicted binding to damaged DNA, proteasome and ubiquitin, and has a putative role in protein deglycosylation and handling of endoplasmic reticulum stress. Thus, it may be related to the response to stress exerted by chlorhexidine, as further indicated by i) it having signs of genomic selection in other *Candida* species[2], ii) null mutants in *C. albicans* having decreased UV resistance (CGD), and iii) null mutants in *S. cerevisiae* leading to reduced tolerance towards various chemical stressors (e.g. $H_2O_2$, cycloheximide, methyl methanesulfonate, mechlorethamine, flucytosine, cisplatin) (SGD).

In addition, *Scer_MNR2* has a predicted magnesium ion transmembrane transporter activity and a putative role in cellular magnesium ion homeostasis, with signs of genomic selection in other *Candida* species[2] that reinforce its importance. Also, consistent with the role of this gene in the adaptation to chemical stress, null mutants in *C. albicans* have increased amphotericin B susceptibility (CGD), while null mutants in *S. cerevisiae* have increased ethanol tolerance and decreased calcium dichloride tolerance (SGD).

On another line, beyond these prioritized genes, we found that the *CDR1B* efflux pump, related to azole resistance (see above) had also high-confidence GWAS hits for chlorhexidine tolerance (**Fig. 7**, **Supplementary Table 4**). While the variants in this gene associated to azole resistance (partial transcript deletion, syn|c.4038|gtG/gtA, mis|p.846|A/T, mis|p.1346|V/A, mis|p.1355|A/G and mis|p.1352|M) are different to those correlated to chlorhexidine tolerance (mis|p.621|F/S and FS|p.563|F/X), it suggests that this efflux pump may play a role in both azole and disinfectant tolerance mechanisms. Accordingly, *C. glabrata* (*Nakaseomyces glabratus*) strains evolved in chlorhexidine acquire *PMA1* and *PDR1* mutations, leading to azole cross-resistance (likely) due to overexpression of *CDR1*[42].

All in all, chlorhexidine tolerance is associated with changes in stress responses (*CPAR2_101330, Scer_RAD23*), DNA replication (*Scer_ORC4*), magnesium homeostasis (*Scer_MNR2*) and drug efflux (*CDR1B*).

##### Patient age groups' genes

For 'age infant' we find associations on three genes, *Scer_PMI40, RBT1* and *ALS11* (**Supplementary Table 6**), which all have clear virulence-related functions. *Scer_PMI40* has a predicted role in cell wall mannoprotein biosynthetic process, hyphal growth, and protein glycosylation. The role in mannoprotein biosynthesis is notable due to the immunogenic nature of mannoproteins[43], suggesting that the variants in this gene may be related to immune evasion. In addition, the ortholog in *C. albicans* is involved in invasive growth, adherence to polystyrene, phagocytosis and spider media-induced biofilm formation (CGD), which all are virulence-related functions. Similarly, *RBT1* is induced in hypoxic growth and biofilm formation, which are important virulence-related processes. Finally, *ALS11* is a putative cell adhesion protein binding to human plasminogen, which is a recognized invasion mechanism in other *Candida* species[44]. Also, null mutants in *C. parapsilosis* have decreased virulence in mouse vaginal infection models (CGD), and this gene is involved in the attachment to human endothelial and epithelial cells[45], which all validate the importance of *ALS11* for virulence.

In contrast, for 'age old' we prioritized associations on three other genes: *Scer_NAT4*, *Scer_ATG20* and *CPAR2_403920* (**Supplementary Table 6**). *Scer_NAT4* has a predicted role in histone acetylation and regulation of ribosomal DNA heterochromatin assembly, with annotations related to protein glycosylation that may be important for the interaction with the immune system[46]. Also, in *C. albicans* this gene is related to white-to-opaque phenotype switching (CGD), and it is regulated by the virulence-associated transcription factor *RLM1*[47], validating its importance. Furthermore, *Scer_ATG20* has a predicted role in autophagy and cytoplasm to vacuole transport by the Cvt pathway, but we could not find any additional evidence linking this gene with virulence mechanisms. Finally, *CPAR2_403920* has no functional annotations, but various lines of evidence suggest its relationship with virulence in the *C. albicans* ortholog *HRQ2*. *HRQ2* is i) induced in rat catheters and spider media- related biofilms (CGD), ii) downregulated by (morphogenesis-regulating) acetylcholine[48], and iii) co-expressed with *PGA34*[49], a virulence factor related to epithelial cell damage (CGD).

In summary, the predisposition towards infecting infants is associated to genes involved in mannoprotein biosynthesis (*Scer_PMI40*), biofilm formation (*Scer_PMI40, RBT1*), invasion (*RBT1, ALS11*), hypoxic growth (*RBT1*) and adhesion to human cells (*ALS11*). Conversely, the predisposition towards infecting older patients is associated with genes involved in protein glycosylation (*Scer_NAT4*), autophagy (*Scer_ATG20*), biofilm formation (*CPAR2_403920*) and morphogenesis (*CPAR2_403920*). Although the sets of genes related to each age group are non-overlapping, they affect certain similar functions (biofilm formation, protein glycosylation and adhesion), suggesting common properties of strains that infect such particularly susceptible patients.

##### Isolation sources' genes

We found 17 genes associated to the predisposition towards infecting various isolation sources ('blood vs other sterile', 'skin exudate', 'bronchial aspirate', 'ascitic fluid', 'blood culture', 'blood' and ‘sterile’) (**Supplementary Table 6**). To keep the most relevant ones, for associations related to ML modelling (for ‘blood vs other sterile’) we discuss below the three genes found by the best parameters: *Scer_MDS3, ALS11* and *ALS7* (**Extended Data Fig. 13**). Also, for associations related to GWAS hits we only discuss genes with a virulence-related functional description (*Scer_DFG10* and ALS7) or with signs of genomic selection in other *Candida* species[2] (*Scer_NTG2, CDR1, LPD1* and *Scer_UTP22*). This means that, for the sake of synthesizing our results, we do not discuss the roles of *Scer_YHB1, Scer_RIX7, TRP5, BCR1, Scer_DFG16, ERG11, RBT1, CapafMp06* and *ATP6*, whose annotations can be found in (**Supplementary Table 6**).

The predisposition to infect blood vs other sterile compartments (trait ‘blood vs other sterile’) can be attributed to variants in *Scer_MDS3* combined with mutations in *ALS11* and/or *ALS7*, which all have clear virulence-related functions. *Scer_MDS3* has a predicted role in TOR signaling, negative regulation of ascospore formation, pseudohyphal growth. In *C. albicans*, it is required for virulence in a mouse model of systemic infection, while null mutants lead to decreased biofilm formation and decreased hyphal growth (CGD), showing that it may be a regulator of key virulence factors. In addition, *ALS11* is related to human cell adhesion and virulence, as described above. Similarly, *ALS7* is a putative cell adhesion protein binding to human plasminogen. Also, various lines of evidence linked this gene to virulence in *C. parapsilosis*, including i) mutations linked to changes in biofilm formation[50], ii) upregulation in adhesion-inducing conditions[51], and iii) adhesion to host epithelial cells[52,53]. Most importantly, we propose that the binding to human plasminogen of *ALS11* and *ALS7* could underlie differential invasion of human tissues. Similarly, we find that *ALS7* variants are associated with the ‘blood culture’ source, which may be related to redundant results on these sources.

Conversely, the predisposition for being found in ‘skin exudate’ is associated with variants in *Scer_DFG10*. This gene has a predicted role in dolichol-linked oligosaccharide biosynthetic process and pseudohyphal growth, which are both roles that may be linked to virulence, as justified next. On the one hand, pseudohyphal growth promotes host tissue disruption and invasion[54], which is key for infection dispersion. On the other hand, the role in dolichol-linked oligosaccharide biosynthetic process is relevant due to the link between dolichol and protein glycosylation[55], which is relevant for interactions with the immune system[46]. Accordingly, genes downregulated during gastrointestinal colonization in *C. albicans* were enriched in protein glycosylation annotations, including *DFG10* among others[56]. This is particularly relevant because glycosylation-defective *C. albicans* mutants generated a decreased inflammatory response[57], and mannosylation-defective strains are more resistant to phagocytosis by neutrophils[58]. Thus, we propose that the observed *Scer_DFG10* variants may alter glycosylation patterns leading to selective colonisation through immune evasion mechanisms.

Furthermore, the predisposition for being found in 'bronchial aspirate' is associated with variants in *Scer_NTG2* and *CDR1*. *Scer_NTG2* has predicted DNA N-glycosylase activity and annotations of DNA damage response. Its *C. albicans* ortholog *NTG1* is induced in spider media-related biofilms and involved in the Base Excision Repair (BER) DNA-damage response pathway (CGD). The relationship of this gene with body compartment selectivity could be explained by the link between BER and immune-related oxidative stress responses[59], and/or by its putative role in biofilm formation. Conversely, *CDR1* (different from *CDR1B*) has xenobiotic transporter activity and role in cellular cation homeostasis. In *C. albicans* it is induced in spider media-related biofilms (CGD), but we could not find additional evidence supporting the role of this gene in virulence.

Finally, the predisposition for invading 'sterile' compartments is associated with variants in *Scer_UTP22* and *LPD1*. These variants are particularly relevant because the capacity to infect sterile compartments is a key virulence property of such strains. *Scer_UTP22* has a predicted role in rRNA processing and tRNA export, and is also related to azole resistance (see above). In *C. albicans* this gene is regulated by the *RLM1* transcription factor, involved in cell wall biogenesis and virulence[47], which may explain why it could be related to differential compartment invasiveness. Conversely, *LPD1* encodes a lipoamide dehydrogenase component (E3) of the pyruvate dehydrogenase and 2-oxoglutarate dehydrogenase multi-enzyme complexes. In *C. albicans* this gene is highly antigenic in human oral infection and murine systemic infection, and induced in macrophages (CGD), so it may be related to immune activation.

In summary, the genes modulating the predisposition towards infecting specific body compartments are involved in pseudohyphal growth (*Scer_MDS3, Scer_DFG10*), host adhesion (*ALS7, ALS11*), biofilm formation (*Scer_MDS3, ALS7, Scer_NTG2, CDR1*), protein glycosylation (*Scer_DFG10*), DNA-damage responses (*Scer_NTG2*), xenobiotic transmembrane transport (*CDR1*), rRNA processing and tRNA export (*Scer_UTP22*) and antigenic properties (*LPD1*).These are all virulence-related functions that may be modulated through mutations, leading to selectivity for specific compartments in certain isolates.

#### On the importance of considering various variant types

Regarding the types of variants, we find that lots of different variants, and not only the traditionally-studied missense mutations, are relevant (**Supplementary Table 6**). On the one hand, we find that only 55 of the 79 prioritized gene-phenotype mappings can be attributed small variants altering the protein sequence (missense mutations, frameshifts or premature termination codons) or CNVs / SVs overlapping transcript coordinates, while the other are related to upstream, downstream or CDS synonymous variants. Specifically, these are related to novel gene-phenotype mappings, including i) *CPAR2_105460* and *CPAR2_105480* with fluconazole and voriconazole resistance, ii) *Scer_ATP10* and *Scer_MOH1* with fluconazole resistance, iii) *CPAR2_204120* and *Scer_TEL1* with voriconazole resistance, iv) *CPAR2_806400* with voriconazole, posaconazole and amphotericin B resistance, v) *CPAR2_105750* and *Scer_ENA2* with 5-FC resistance, vi) *Scer_ORC4* with chlorhexidine tolerance, vii) *Scer_ATG20*, *CPAR2_403920* and *Scer_NAT4* with 'age old', viii) *Scer_YHB1* and *Scer_RIX7* with 'ascitic fluid' source, ix) *ALS7* with 'blood culture' source, x) *RBT1* and *Scer_MDS3* with 'blood vs other sterile' source, and xi) *LPD1* and *Scer_UTP22* with 'sterile' source. This suggests that non-coding variants, perhaps through altering gene regulation, may play a significant role in shaping these clinically-relevant phenotypes. Also, this observation highlights the importance of moving beyond the traditional focus on exploring only non-synonymous variation in the study of genotype-phenotype relationships in *Candida*.

On the other hand, we find 20 of 79 gene-phenotype mappings related to CNVs and SVs, showing the importance of considering these complex, often overlooked, variants. Some of these involve the association between *ERG11* whole-gene duplication and pan-azole resistance (**Supplementary Table 6, Fig. 3**). Furthermore, we find some novel gene-phenotype mappings involving CNVs and SVs. These include i) *CPAR2_204210* partial transcript deletion associated with fluconazole, voriconazole and amphotericin B resistance, ii) *Scer_NCP1* whole-gene duplication associated with pan-azole and amphotericin B resistance, iii) *CDR1B* partial transcript deletion associated with voriconazole resistance iv) *Scer_NST1* whole-gene duplication associated with posaconazole resistance v) *Scer_FOL1* whole-gene duplication associated with amphotericin B resistance, vi) *CDR1* partial transcript deletion associated with 'bronchial aspirate' source, vii) *BCR1* transcript-breaking SV associated with 'blood vs other sterile' source, viii) *ERG11* whole-gene duplication associated with 'blood vs other sterile' source, ix) *RBT1* 5' duplications associated with 'blood vs other sterile' source, x) *CapafMp06* partial transcript duplication associated with 'blood' source, and xi) *ATP6* partial transcript duplication associated with 'blood' source.

These findings highlight the importance of regulatory variants, CNVs and SVs, and showcase the power of our approach for analyzing them in a systematic straightforward manner.

### SUPPLEMENTARY MATERIALS AND METHODS

#### Read depooling pipeline

To verify that the sequencing datasets analyzed here contain only *C. parapsilosis* reads we ran a depooling pipeline on a subset of 100,000 read pairs for each strain, mapping them to the genomes of five *Candida* species (*C. parapsilosis*, *C. auris*, *C. orthopsilosis*, *C. metapsilosis* and *C. albicans*) (see **Methods**). In brief, we used bwa mem (v0.7.17) to align the reads to a concatenated reference genome including the five pooled species, generated using biopython (v1.76). The references for individual species were obtained as described in the **Methods** section "Obtention of genomes, annotations and databases". We next separated the reads uniquely mapping to each taxa with samtools (v1.9), and used python to calculate the fraction of reads belonging to each species based on these uniquely-mapped reads.

#### Convergence GWAS methodological details

This section explains in detail how we performed GWAS analyses. For this, we developed and used the ancestral_GWAS toolkit, based on [2], available at https://github.com/Gabaldonlab/ancestral_GWAS_toolkit. We ran the scripts included in this toolkit on the ancestral_GWAS_env conda environment available in this repository (see main **Methods**). Note that the scripts of this toolkit mostly rely on calling functions from the python module ancestral_GWAS_functions.py, and in the sections below we refer to specific functions in this module as GWASfun.<function name>. Also, as in **Methods**, we refer to functions inside Cparapsilosis_popgen_functions.py, from https://github.com/Gabaldonlab/Cparapsilosis_GenoPheno, as CPfun.<function name>.

##### Preparation of annotation datasets

For running the GWAS we generated various annotation datasets with scripts of the ancestral_GWAS toolkit. To enable variant collapsing at the domain level and to get per-gene Reactome / MetaCyc annotations we generated InterProScan annotations for the proteins encoded in the gff annotations with run_interproscan_from_gff.py. Specifically, to enable proper InterProScan running we first added the InterProScan_env directory path to the $LD_LIBRARY_PATH variable with 'export LD_LIBRARY_PATH=<InterProScan_env dir>/lib:$LD_LIBRARY_PATH'. Also, we defined the environmental variable $INTERPROSCAN_SH, used in run_interproscan_from_gff.py to run the tool, with 'export INTERPROSCAN_SH="source <conda dir>/etc/profile.d/conda.sh && conda activate InterProScan_env && <InterProScan v5.52-86.0 dir>/interproscan.sh"'. Then, we ran run_interproscan_from_gff.py with arguments '--gDNA_code 12 --mtDNA_code 4' (env:ancestral_GWAS_env), which generates the protein domain annotations into 'interproscan_annotation.out'.

Furthermore, to transform pathway annotations (from GO, Reactome and MetaCyc, described above) to a compatible format we used prepare_pathway_annotations.py. Specifically, to enable processing of MetaCyc annotations we executed this script while the Pathway Tools software was running (with '<Pathway Tools dir>/pathway-tools -lisp -python'). Also, to enable filtering out MetaCyc annotations outside Ascomycota taxons we obtained the NCBI Taxonomy database[60] files ('taxa.sqlite' and 'taxa.sqlite.traverse.pkl') by running 'python -c "from ete3 import NCBITaxa; ncbi = NCBITaxa(); ncbi.update_taxonomy_database()"' on 28 June 2024. Then, we ran prepare_pathway_annotations.py with arguments '--interproscan_output interproscan_annotation.out --reactome_pathways ReactomePathways.txt --reactome_pathway_relations ReactomePathwaysRelation.txt --max_fraction_genes_pathway 0.05 --reactome_pathways_keep Candida --MetaCyc_pathways_clade Ascomycota --obo_file go-basic.obo --GOterms gene_association.20231003.cgd.gz --GOterms_type gene_association_CGD --NCBI_taxa_db taxa.sqlite' (env:ancestral_GWAS_env).

This run of prepare_pathway_annotations.py generated various relevant datasets. The files 'annotations_GO.tab', 'annotations_MetaCyc.tab' and 'annotations_Reactome.tab' contain the per-gene pathway annotations, with various relevant information added, including i) the GO namespace (BP, CC or MF), ii) the description, iii) the fraction of genes with the annotation, iv) the organism from which the annotations come in Reactome, and v) various pathway metrics (number of children, number of parents, GO term depth and GO term level). Also, the files 'MetaCyc_pathway_info.tab' and 'MetaCyc_pathway_relations.tab' contain the MetaCyc pathway descriptions, NCBI taxon IDs ('MetaCyc_pathway_info.tab') and child-parent MetaCyc pathway relations as generated by Pathway Tools ('MetaCyc_pathway_relations.tab').

Note that while GO terms originate from CGD annotations, Reactome and MetaCyc terms derive frominterproscan_annotation.out passed through --interproscan_output. Also, note that this script filtered out pathways that are i) too general, in >5% of genes due to --max_fraction_genes_pathway 0.05, ii) not annotated in Ascomycota taxons according to the NCBI Taxonomy (MetaCyc annotations only) due to --MetaCyc_pathways_clade Ascomycota, iii) coming from an organism that is not *Schizosaccharomyces pombe* or *Saccharomyces cerevisiae* (Reactome annotations only) due to

--reactome_pathways_keep Candida, or iv) not biologically meaningful for *Candida* species (Reactome annotations only) due to --reactome_pathways_keep Candida.

##### Tree reconstruction for each GWAS

To generate optimal strain trees for subsequent GWAS, for each phenotype and aligner (target_aligner) we reconstructed a strain tree for the subset of the isolates that have a clearly defined phenotype (with values of 1 or 0), called “target runs”. For this, we ran CPfun.generate_tree_one_phenotype_and_aligner (see **Methods**) with arguments min_coverage_pos = 12 and target_runs = set of target runs for a given phenotype. This function first loads the tree of all 367 high-confidence isolates generated with diploid mode and the corresponding target_aligner (see ‘Tree reconstruction for all isolates’ from **Methods**), and generates a pruned tree with only the target runs. Next, this pruned tree is used to define an outgroup clade within the tree of target runs. Then, the function runs CPfun.get_cmds_tree_reconstruction_one_configuration (see **Methods**) on the target runs (argument runIDs_para) and the identified outgroup (argument outgroup), with mode = “diploid”, aligner = target_aligner, min_coverage_pos = 12, min_support_trees = 90. Finally, from the resulting tree reconstruction metrics, this function validates that either fraction_trees_expected_outgroup or fraction_trees_expected_most_frequent_outgroup (defined in **Methods**) are 1.0 (for quality control of the outgroup), and keeps the resulting rooted tree (raw, with no collapsed nodes) for further analyses.

##### Defining GWAS sample sets

We considered various subsets of isolates for GWAS for each phenotype and aligner. Initially, we considered all isolates for which the phenotype was clearly defined (with values 1 or 0), with their corresponding tree (generated as described in the previous section), which corresponds to sample_set = all_samples. Also, to avoid the biases generated by clonal redundancy (**Fig. 2**), we considered a subset of such isolates where we only keep three samples for each clade that is monophyletic for a given phenotype value (1 or 0), corresponding to sample_set = representative_samples (**Extended Data Fig. 6a**). To find these representative samples we ran get_balanced_samples.py with arguments ‘--tree_file <tree all samples phenotype> --min_support 70 --ASR_method DOWNPASS --nrepresentative_samples 3’ (env:ancestral_GWAS_env), using CPfun.get_balanced_samples_for_GWAS. Note that this script also outputs the number of phenotype transitions in the tree of representative samples, used to quantify phenotypic variation

(**Fig. 2**), calculated in a way that all nodes are considered for the calculation. Specifically, for nodes where the DOWNPASS ASR yields uncertain estimates, the most common phenotype across the leaves is assigned, enabling the inference transition / no transition in all nodes. Furthermore, we considered the sample_set = balanced_samples, which includes a similar subset of isolates where we only keep one sample per monophyletic clade, generated with an equivalent get_balanced_samples.py command, but using '--nrepresentative_samples 1'.

Finally, to enable convergence GWAS running we obtained a strains tree for each of these subsets of isolates (representative and balanced) with CPfun.generate_tree_one_phenotype_and_aligner (see **Methods**), run with aligner = target_aligner, target_runs = representative or balanced set of runs, min_coverage_pos = 12. All code to define these sample sets and reconstruct their corresponding trees can be found in CPfun.get_cmds_convergence_GWAS.

##### Preparing GWAS input files

To prepare the input files for run_GWAS.py, for each phenotype, aligner and sample set (all_samples, representative_samples, balanced_samples) we ran the script prepare_GWAS_inputs.py from the ancestral_GWAS toolkit (env:ancestral_GWAS_env). Specifically, this script generates the filtered variants (small, SV, CNV and aneuploidies), variant group definitions (e.g. variants affecting each domain, gene or pathways, see **Fig. 4b**), properly formatted tree, phenotype ASR and variant ASR calculations, and we ran it with various custom arguments explained next. To focus the analysis on the sample_set of interest we provided the phenotypes and tree for each subset of samples through --phenotypes and --treefile. In addition, to get high-confidence filtered small variants we set '--integrated_small_vars integrated_small_variants.tab --small_vars_filtering typical_diploid --small_vars_min_cov_pos 12 --repeats_file <simple repeats>'. These arguments configure the pipeline to i) load the raw variants 'integrated_small_variants.tab' (described in **Methods**) and ii) filter them in a diploid-like manner, keeping variants that are similar to 'variants_atLeast2PASS_ploidy2.vcf', but discarding those overlapping simple repeats.

Similarly, to get high-confidence filtered SVs and CNVs we set the arguments '--integrated_SV_CNVs integrated_SVs_CNVs.tab --SV_min_minAF 0.1 --SV_min_maxAF 0.3 --CNV_max_relative_CN_deletion 0.5 --CNV_min_relative_CN_duplication 1.5 --CNV_max_relative_coverage_deletion 0.6 --CNV_min_relative_coverage_duplication 1.3 --CNV_min_len 600'. These arguments configure the pipeline to i) load the SVs and CNVs from ‘integrated_SVs_CNVs.tab’ (described above) and ii) filter them in a diploid-like manner, similar to previous similar approaches[2]. Specifically, we kept SVs that have a VAF >=0.1 in all underlying breakends (--SV_min_minAF 0.1), and that have at least one breakend with VAF >= 0.3 (--SV_min_maxAF 0.3). Also, we kept all types of CNVs (complete deletions, monosomies, trisomies and tetrasomies) (--CNV_max_relative_CN_deletion 0.5 --CNV_min_relative_CN_duplication 1.5 ) of at least 600 bp (--CNV_min_len 600), but ensuring deletions have a maximum relative coverage of 0.6 (--CNV_max_relative_coverage_deletion 0.6) and duplications have a minimum relative coverage of 1.3 (--CNV_min_relative_coverage_duplication 1.3).

Furthermore, to get calls of whole-chromosome aneuploidies we set arguments ‘--paths_CNV_calling <paths to call_CNVs output directory> --aneuploidies_max_fraction_N_bases 0.1 --aneuploidies_max_fraction_repeats 0.1 --aneuploidies_min_median_mappability 0.75 --min_fraction_chromosome_aneuploidy 0.5 --aneuploidies_cov_thresholds 0.6,1.4’. These arguments configure the pipeline to i) load the relative coverage per window measurements generated by call_CNVs from --paths_CNV_calling (described above), ii) filter out regions with a fraction of N bases > 0.1 (--aneuploidies_max_fraction_N_bases 0.1), a fraction of repeats > 0.1 (--aneuploidies_max_fraction_repeats 0.1) or a median mappability < 0.75 (--aneuploidies_min_median_mappability 0.75) and iii) identify aneuploid chromosomes with at least 50% of the filtered windows (--min_fraction_chromosome_aneuploidy 0.5) deleted (relative coverage < 0.6) or duplicated (relative coverage > 1.4) (--aneuploidies_cov_thresholds 0.6,1.4).

To consider functional annotations for small variants, SVs and CNVs we passed the previously generated files (see above) with ‘--small_vars_annot small_vars_annot/annotated_variants_corrrectedGene.tab --SV_CNVs_annot SV_CNV_annotation/annotated_variants_corrrectedGene.tab’. Also, to process the annotations with the adequate translation codes we set ‘--gDNA_code 12 --mtDNA_code 4’. Furthermore, we used the default values for --truncating_consequences, --non_truncating_consequences, --synonymous_consequences and --non_synonymous_consequences, which are related to how we define truncating and non-synonymous variants in variant collapsing (see **Fig. 4 b**).

Furthermore, to consider InterProScan annotations for domain-level variant grouping we provided the ‘interproscan_annotation.out’, described above, through --interproscan_output. Similarly, to enable pathway-level collapsing (GO, Reactome and MetaCyc), we provided the output directory of prepare_pathway_annotations.py (see above) through --pathway_annotations_dir. Conversely, to keep only groups of at least 2 variants, and that have variants in at least 2 samples, we set the arguments '--min_nmuts_per_group 2 --min_nsamples_per_group 2'.

This prepare_GWAS_inputs.py run generated various relevant files. The table 'all_groupings.tab' contains the link between groups and variants, where each group can be tested in a subsequent GWAS. Each row corresponds to a combination of a variant (variantID_across_samples, e.g. SNP Contig005504_C_parapsilosis_CDC317_103394_C/A), a grouping strategy (grouping_ID, e.g. collapsing non synonymous small variants at the gene level) and a group name (e.g. gene CPAR2_600430). Also, the fields type_vars, type_mutations and type_collapsing indicate the important aspects of each grouping strategy **(Fig. 4a,b**). Specifically, type_vars refers to the variant types considered, and it can be small variants, SVs, CNVs or any combination thereof. Also, type_mutations refers to the types of mutational effects considered, and it may be all, non synonymous, non synonymous non truncating or truncating. Furthermore, type_collapsing reflects the level at which the variants were collapsed, including variant effects (e.g. *ERG11* variant G458S), domains, genes, GO, Reactome, MetaCyc or none (for single-variant groups). Finally, the column 'linkage_groupID' contains an ID reflecting groups that are equivalent, i. e. having the exact same presence / absence pattern of the underlying variants, which is useful for further avoiding unnecessary computational burdens.

Furthermore, the table 'variants_IDmapping.tab' includes a mapping between each variant and a corresponding numerical ID, which is useful to speed up downstream analysis. Also, the file 'mutation_to_representative_mutation.tab' includes a definition of groups of variants with equivalent presence / absence pattern across the considered isolates, i.e. fully linked, which is useful to avoid redundant computations in downstream analyses. In addition, 'corrected_rooted_tree.nw' contains a tree that is properly formatted for further GWAS, with i) a support of 100 for the root and each of its child nodes and ii) branch length with an added pseudocount (10% of the minimum branch length across tree leaves) to avoid having branches with length = 0.0. Also, the file 'phenotypes_ASR.out' and and folder 'ASR_all_variants/' contain the ASR results for the phenotypes and all variants, respectively, for different ASR methods, including the DOWNPASS maximum parsimony[61] and the MPPA maximum likelihood[61] methods. Finally, the file 'resampled_phenotypes_pastml_df.py' contains equivalent ASR results for 10,000 randomly resampled phenotypes, as generated by get_resampled_phenotypesASR.py of the ancestral_GWAS toolkit (env:ancestral_GWAS_env), which are necessary for further empirical p value calculations.

All of the code to prepare these GWAS input files can be found in CPfun.get_cmds_convergence_GWAS.

**Running GWAS**

For each phenotype, aligner, sample_set and various combinations of min_support (10, 50, 70, 90) and ASR_method ("MPPA", "DOWNPASS", "MPPA,DOWNPASS"), we ran run_GWAS.py (env:ancestral_GWAS_env) with various arguments, described next. To run the GWAS on the generated input files (groups, corrected tree, variant ID mappings and ASR results) of each sample set we used the arguments '--input_variant_groups all_groupings.tab --treefile corrected_rooted_tree.nw --ASR_phenotypes phenotypes_ASR.out --ASR_variants_dir ASR_all_variants/ --varID_mapping variants_IDmapping.tab --varID_mapping_rep mutation_to_representative_mutation.tab --resampled_phenotypes_pastml_out resampled_phenotypes_pastml_df.py'.

Also, to obtain GWAS results on different branch supports and ASR methods we set them through '--min_support <min_support> --ASR_method <ASR_method> --classification_method_ASR all_ignoreNaN'. Specifically, min_support refers to the minimum branch support that a node should have to be considered for GWAS, and we tried values 10, 50, 70 and 90. Also, --ASR_method was either MPPA, DOWNPASS (defined above) or a consensus between the two (MPPA,DOWNPASS). Due to '--classification_method_ASR all_ignoreNaN', the consensus state would be 1 (if both MPPA and DOWNPASS were 1, MPPA was 1 and DOWNPASS was NA or MPPA was NA and DOWNPASS was 1), 0 (if both MPPA and DOWNPASS were 0, MPPA was 0 and DOWNPASS was NA or MPPA was NA and DOWNPASS was 0) or NA if none of these conditions were met.

On another line, to record different types of genotype-phenotype associations we set '--interesting_gwas_methods synchronous,phyC,phylogeny_agnostic'. For gwas_method = 'synchronous', our pipeline measures whether nodes with genotype transitions (with gain or loss of a variant in the tested group) are more likely to have phenotype transitions (gain / loss of the phenotype) than expected by chance, which is equivalent to the 'synchronous' method of hogwash[62,63]. Thus, for this gwas_method the 'phenotype' is phenotype transition, and the 'genotype' is genotype transition. Conversely, for gwas_method = 'phyC' it measures whether the variants of the tested group are more likely to be acquired in nodes with the phenotype vs those without the phenotype, following the 'phyC' algorithm[63,64]. Thus, for this gwas_method the 'phenotype' is phenotype presence, and the 'genotype' is genotype acquisition. Finally, for gwas_method = 'phylogeny_agnostic' our pipeline calculates whether there is an association between having a variant in the tested group and having the phenotype, only across isolates and ignoring their phylogeny, so it is equivalent to classical allele-counting GWAS[65]. Thus, for this gwas_method the 'phenotype' is phenotype presence, and the 'genotype' is genotype presence. Also, we set '--min_n_geno_transitions 2 --min_n_geno_gains 2 --min_n_geno 2' to only test groups of variants with >=2 genotype transitions (gwas_method = 'synchronous'), >=2 variant acquisitions (gwas_method = 'phyC') or >=2 isolates with a variant (gwas_method = 'phylogeny_agnostic').

Additionally, to enable further parameter benchmarking and hit interpretability (described below) we used '--cladeID_info cladeID_Bergin2022_info.tab' to record the clades (according to the file 'cladeID_Bergin2022_info.tab' described in **Methods**) where convergence was observed. Specifically for 'synchronous' and 'phyC' methods, this pipeline identifies nodes with genotype-phenotype convergence (defined differently for each method, as described above), and records the clades of these nodes if they don't include isolates of various clades. This allowed us to identify hits which show convergence across multiple clades, that are of higher confidence as compared to hits that only show convergence within a single clade, which was useful for further parameter benchmarking. Furthermore, to only calculate association empirical p values that are based on phenotype resampling we set '--all_pval_methods phenotypes'. Note that, due to how we define transitions in convergence GWAS[2], loss-of-heterozygosity events were not considered as separate instances and thus were not analyzed specifically unless they involve a variant gain or loss.

Most importantly, run_GWAS.py calculates various association metrics for each gwas_method and group, based on the tree nodes in which phenotype and genotype (which depend on the gwas_method, described above) are properly resolved (with 0 or 1). For instance, nodes with properly resolved 'phenotype presence' or 'genotype presence' are those that have support >= min_support, and have 0 or 1 ASR state. Conversely, nodes with properly resolved 'phenotype transition', 'genotype transition' or 'genotype acquisition' are those where the 'phenotype presence' or 'genotype presence' are properly resolved in them and their parent nodes. At the end, our pipeline generates the 'phenotypes_info.tab' file with the phenotype and phenotype transition information for all tree nodes, used for GWAS filter benchmarking (see **Methods**) and for visualization of numbers of nodes (**Fig. 4c,d**). Also, it generates the 'integrated_GWAS_stats.tab' file with all GWAS association metrics, with one row for each combination of gwas_method, grouping strategy (column grouping_ID) and group ID (column group). In the following paragraphs we describe the relevant columns of this table, used in subsequent analyses.

The columns 'nodes_withGeno_clearNodes' ($n_G$) and 'nodes_withPheno_clearNodes' ($n_P$) indicate the number of nodes with the genotype or the phenotype, respectively. For instance, for gwas_method = 'synchronous' it reflects the number of nodes with genotype transitions or phenotype transitions, respectively. Note that, due to the arguments '--min_n_geno_transitions 2 --min_n_geno_gains 2 --min_n_geno 2' (see above), we only tested groups with $n_G >= 2$. In addition, the column 'nodes_GenoAndPheno' ($n_{G\&P}$) indicates the number of nodes with both genotype and phenotype. For instance, for gwas_method = 'synchronous' it reflects the number of nodes with both genotype transition and phenotype transition. Similarly, the columns 'nodes_noGenoAndPheno' ($n_{NG\&P}$), 'nodes_GenoAndNoPheno' ($n_{G\&NP}$) and 'nodes_noGenoAndNoPheno' ($n_{NG\&NP}$) indicate the numbers of nodes with no genotype and phenotype, genotype and no phenotype and lack of both genotype and phenotype, respectively. Also, the column 'nodes_withoutGeno_clearNodes' ($n_{NG}$) indicates the number of nodes without the genotype.

Furthermore, the column 'chi_square' ($X^2$) includes the $X^2$ statistic of the contingency table [[$n_{G\&P}$ , $n_{G\&NP}$] , [$n_{NG\&P}$ , $n_{NG\&NP}$]], calculated with scipy.stats.chi2_contingency (1.7.3), reflecting genotype-phenotype associations. Similarly, the column 'OR' (*OR*) includes the odds-ratio of this contingency table. Also, the column 'epsilon' ($\varepsilon$) includes the convergence level between genotype and phenotype[2,63], calculated as $\varepsilon = (2 * n_{G\&P}) / (n_G + n_P)$, as proposed before[2,63]. For instance, gwas_method = 'synchronous' conceptually reflects how often a genotype transition is associated with a phenotype transition and *vice versa*. Also, the columns 'cladeID_names_GenoAndPheno' and 'node_names_GenoAndPheno' reflect the clade IDs and node names in which the genotype-phenotype convergence was observed.

The columns 'pval_chi_square_phenotypes' ($p(X^2)$) and 'pval_GenoAndPheno_phenotypes' ($p(n_{G\&P})$) include empirical p values reflecting probability of getting the observed $X^2$ or $n_{G\&P}$ by chance, respectively. Specifically, they reflect the fraction of 10,000 randomly resampled phenotypes (from 'resampled_phenotypes_pastml_df.py') that lead to a $X^2$ or $n_{G\&P}$ above or equal to the observed one, while setting the 0.0 to a pseudocount of 5e-05. Similarly, the column 'pval_fisher' ($p_{FISHER}$) includes the Fisher p value of the [[$n_{G\&P}$ , $n_{G\&NP}$] , [$n_{NG\&P}$ , $n_{NG\&NP}$]] table, calculated with scipy.stats.fisher_exact. Also, the columns 'pval_chi_square_phenotypes_bonferroni', 'pval_chi_square_phenotypes_fdr_bh', 'pval_GenoAndPheno_phenotypes_bonferroni' and 'pval_GenoAndPheno_phenotypes_fdr_bh' include the bonferroni or FDR-corrected $p(X^2)$ $p(n_{G\&P})$,

respectively. For these, the correction is applied across independent groups (with different linkage_groupID) of a given grouping strategy (grouping_ID).

The columns 'pval_chi_square_phenotypes_maxT' ($p(X^2)_{maxT}$) and 'pval_epsilon_phenotypes_maxT' ($p(\varepsilon)_{maxT}$) include the maxT-corrected p values[2,66]. These are calculated as the fraction of 1,000 randomly resampled phenotypes that lead to a maximum $X^2$ or $\varepsilon$ across groups of a given grouping strategy (grouping_ID) that is above or equal to the observed one.

The columns 'pval_chi_square_phenotypes_bonferroni_allGroups', 'pval_chi_square_phenotypes_fdr_bh_allGroups', 'pval_GenoAndPheno_phenotypes_bonferroni_allGroups', 'pval_GenoAndPheno_phenotypes_fdr_bh_allGroups', 'pval_chi_square_phenotypes_maxT_allGroups' and 'pval_epsilon_phenotypes_maxT_allGroups' include equivalent corrected p values, but where the correction is applied across all independent groups (with different linkage_groupID) together, which is a more conservative correction scope.

All the necessary code can be found in CPfun.get_cmds_convergence_GWAS.

##### Defining non-redundant GWAS hits

The 'raw' lists of filtered GWAS hits (see **Methods**) included many redundant groups, as the same variant may be found in different groups. To address this and generate a final set of non redundant hits, as found in **Supplementary Table 4,5**, we ran CPfun.get_NR_GWAS_tables on the lists of raw low and high confidence hits. This function runs, for the GWAS hits obtained on each phenotype and type of filtering (low or high confidence) the ancestral_GWAS toolkit script generate_NR_groups.py (env:ancestral_GWAS_env). Specifically, we used arguments '--gwas_hits <raw filtered hits> --custom_geneID_maps Scer_HAL9:TAC1 --gwas_inputs_dir <prepare_GWAS_inputs.py output directory> --pathway_annotations_dir <prepare_pathway_annotations.py output directory> --CGD_chromosomal_features <CGD tabular file> --interproscan_output interproscan_annotation.out --initial_sorting_fields epsilon,OR,nodes_GenoAndPheno --initial_sorting_ascending False,False,False --reactome_organism_priority Saccharomyces_cerevisiae,Schizosaccharomyces_pombe --keep_variants_w_transitions --min_support <70 (high confidence) or 10 (low confidence)> --ASR_method <MPPA,DOWNPASS (high confidence) or MPPA (low confidence)>'.

With these arguments, generate_NR_groups.py first calculates, for all variants belonging to groups with raw GWAS hits (raw groups), the maximum *ε*, *OR* and $n_{G\&P}$ obtained across raw groups including each variant. Then, it iterates through each of these variants hierarchically sorted by the maximum *ε*, *OR* and $n_{G\&P}$ values (in a descending manner) to progressively create a set of non-redundant groups, so that a given variant will be in only one non-redundant group. Specifically, our pipeline performs various steps in each new iteration, yielding a single new non-redundant group related to each new variant. First, it keeps raw groups that have no variants already within some defined non-redundant group, ending the redundancy-reduction iterations if there are no such raw groups left. Second, out of these raw groups, it identifies those that have the variant, hierarchically sorting them by *ε*, *OR* and $n_{G\&P}$ in a descending manner, and keeping the ones that have the same *ε*, *OR* and $n_{G\&P}$ values as the first group of the sorted list. For instance, two fully-redundant groups will yield the exact same statistics. Third, our pipeline keeps one representative out of these fully-redundant raw groups and adds it to the final list of non-redundant groups, using the function GWASfun.get_NR_group_series. In brief, this function applies a hierarchical redundancy reduction pipeline equivalent to the one in [2], where the most specific groups are prioritized over the more general ones.

Finally, generate_NR_groups.py generates a table where each row corresponds to one combination of GWAS non-redundant group, related variant and affected genes, such as those in **Supplementary Tables 4,5**. Many of these steps were modulated by the arguments mentioned next. To consider, across the whole pipeline, only variants within a group that transition (are gained / lost) with the phenotype we set '--keep_variants_w_transitions'. Similarly, to enable these calculations of transitions we set '--min_support 70 --ASR_method MPPA,DOWNPASS' (high-confidence hits) or '--min_support 10 --ASR_method MPPA' (low-confidence hits). Also, to perform a descending hierarchical sorting of redundant groups based on their *ε*, *OR* and $n_{G\&P}$, we set '--initial_sorting_fields epsilon,OR,nodes_GenoAndPheno --initial_sorting_ascending False,False,False', but these could be changed in further analyses. Furthermore, to re-define the gene name *Scer_HAL9* to *TAC1*, which are equivalent, in the output tables we set '--custom_geneID_maps Scer_HAL9:TAC1'.

Conversely, many arguments are related to the processing of functional annotations and their relationship with the redundant group prioritization performed in GWASfun.get_NR_group_series. For instance, to add gene metadata and enable gene-based prioritization we passed to --CGD_chromosomal_features the CGD tabular gene annotations file (see section 'Obtention of genomes, annotations and databases'). Similarly, to prioritize *S. cerevisiae* over *S. pombe* Reactome pathways, in the case of redundancy, we set '--reactome_organism_priority Saccharomyces_cerevisiae,Schizosaccharomyces_pombe'. Also, to process groups related to InterProScan annotations (domain-level collapsing) we provided the 'interproscan_annotation.out', described above, through --interproscan_output. Finally, to process groups related to pathway-level collapsing (GO, Reactome and MetaCyc), we provided the output directory of prepare_pathway_annotations.py (see above) through --pathway_annotations_dir.

#### Methodological details for the ML classifiers

This section explains in detail how we performed all ML analyses, referring to the relevant GWASfun functions, CPfun functions, and ancestral_GWAS toolkit scripts. Also, note that we refer to various datasets generated for the GWAS (see above), which were key inputs of the ML pipeline. The code to reproduce all of these analyses is in CPfun.run_ML_predictors_phenotypes. To follow the sections below, make sure to carefully read first the **Methods** section 'Building Machine Learning (ML) classifiers', which provides a necessary overview.

##### Defining target gene sets

To enable prioritization of redundant features during ML training (train parameter 'prioritize_expected_genes', see **Methods**) we defined a set of expected genes for azole and echinocandin resistance phenotypes, using CPfun.get_phenotype_to_expected_genes. Specifically, this function defines as 'expected genes' for azole resistance phenotypes the genes *ERG11*, *MRR1*, *Scer_HAL9 (TAC1)* and *NDT80*. Similarly, it defines as 'expected genes' for echinocandin resistance traits the genes *FKS1* and *FKS2*. Also, this function returns a mapping between these gene names and their gene IDs, as found in the CGD tabular features file.

To reduce overfitting, we explored whether building the classifiers on features related to only certain genes of interest would yield good models (test parameter 'gene_set', see **Methods**). To define such genes of interest we considered different combinations of the gene sets 'known', 'by_description', 'GWAS_high', 'GWAS_low', 'selection_all' and 'selection_SNP', described next. The 'known' set includes genes expected to be related to resistance towards azoles (*ERG11, ERG3, TAC1, MRR1, NDT80* and *UPC2*) or echinocandins (*FKS1, FKS2* and *ERG3*), which only apply for these phenotypes. Also, the 'by_description' set includes genes with a functional description in CGD that is related to the phenotype of interest, which we defined based on manual curation. For azole resistance traits, this included genes with the text 'azole' or 'ergosterol' in the description, excluding (after manual curation) the genes *Scer_THI6, Scer_ADE17, Scer_ADE13, HIS3, Scer_THI4, Scer_ADE1, Scer_HIS6, HIS7* and *Scer_ADE2*. For echinocandin resistance traits, this included genes with matching any of the text 'echinocandin', 'fungin', 'beta-d-glucan', 'beta-glucan', 'beta glucan' or 'drug', excluding *Scer_ERG1*. Finally, for virulence and clinical phenotypes (invasiveness, patient age groups, microfluidic behavior, isolation source, biofilm and pseudohyphae formation), this included genes matching any of the text 'virulence', 'virulen', 'biofilm', 'invasive', 'morphology', 'invasion', 'adhesion', 'adhesin', 'adherence', 'pseudohyphae', 'hypha', 'filamentous', 'immun' or 'host'.

Conversely the sets 'GWAS_high' and 'GWAS_low' include genes with orthologs in *C. albicans, C. glabrata* or *C. auris* with GWAS hits on azole / echinocandin resistance[2], depending on the phenotype. The 'GWAS_high' / 'GWAS_low' set refers to the usage of high-confidence or low-confidence hits, respectively, as defined in [2]. Finally, the set 'selection_all' includes genes with orthologs (in other *Candida* species) having signs of recent genomic selection by SNPs, INDELs, duplications or deletions, according to [2]. Similarly, 'selection_SNP' is a subset of such genes, with orthologs having signs of selection by SNPs. To define all these gene sets for each phenotype we used CPfun.get_phenotype_to_all_possible_genes_for_ML.

Finally, we used CPfun.get_all_gene_sets_combinations to define the actual combinations of these gene sets, resulting in the different gene sets tried for training and evaluation (test parameter 'gene_set'). For resistance phenotypes we tried the combinations of sets "known", "known+by_description", "known+GWAS_high", "known+GWAS_high+GWAS_low", "known+by_description+GWAS_high", "known+by_description+GWAS_high+GWAS_low", "known+selection_all", "known+selection_SNP", "known+by_description+selection_all", "known+by_description+selection_SNP", "known+by_description+GWAS_high+ selection_SNP" and "known+by_description+GWAS_high+GWAS_low+selection_all". For virulence and clinical phenotypes we tried the combinations "by_description", "by_description+selection_all" and "by_description+selection_SNP". These differences between gene_set across phenotypes explain why there are different 'test parameters'.

##### Preparing ML input files

To prepare the necessary input files for ML training and evaluation we used prepare_ML_classifier_inputs.py (env:ancestral_GWAS_env). To enable further hyperparameter tuning we ran this for each phenotype and combinations of various parameters, including the aligner, min_support (50, 70 or 90), ASR_method (MPPA, DOWNPASS or 'MPPA,DOWNPASS') and transition_definition ('each_pheno_clade' or 'two_close_balanced'), described in **Methods** and referred below. The main input for this pipeline was 'input_paths.tab', a table with one row per sample_set (all or representative samples), with columns corresponding to files and folders generated for GWAS for the corresponding aligner, min_support, ASR_method and sample_set. These include i) a file with phenotype presence / absence pattern across isolates, ii) the output directory of prepare_GWAS_inputs.py (see above) and iii) the 'integrated_GWAS_stats.tab' raw GWAS results (see above).

For each phenotype, aligner, min_support, ASR_method and transition_definition we set the following arguments for prepare_ML_classifier_inputs.py: '--min_support <min_support> --ASR_method <ASR_method> --classification_method_ASR all_ignoreNaN --paths_file 'input_paths.tab' --min_pheno_transitions 5 --pathway_annotations_dir <prepare_pathway_annotations.py output directory> --interproscan_output interproscan_annotation.out --CGD_chromosomal_features <CGD tabular file> --reactome_organism_priority Saccharomyces_cerevisiae,Schizosaccharomyces_pombe --n_phenotype_resamples 10 --n_CVset_resamples 5 --min_geno_transitions_GWAS_hit 2 --min_geno_pheno_transitions 2 --transition_definition <transition_definition>'. This pipeline processes the groups, variants and trees generated for GWAS (see above), to generate features and train / evaluation / test splits necessary for further ML modelling. In the next paragraphs we describe how it works with the specified arguments.

Initially, prepare_ML_classifier_inputs.py processes the input files provided by --paths_file with GWASfun.check_df_paths_ML_predictors and GWASfun.get_df_gwas_hits_filtered_for_ML_predictors, yielding a set of variant groups with sufficient genotypic variation to be possible ML predictive features. These include those that i) come from a sample_set with >= 5 phenotype transitions (due to '--min_pheno_transitions 5') and ii) for gwas_method = 'synchronous' have $n_G$ >= 2 (due to '--min_geno_transitions_GWAS_hit 2') and $n_{NG}$ >= 1. Then, it runs a redundancy-reduction procedure on these groups (see GWASfun.get_df_groups_NR_by_linkage_each_sample_set) to keep a representative subset of groups, for each sample set, that have a unique presence / absence pattern of the underlying variants across isolates, using the function GWASfun.get_NR_group_series for prioritizing the most specific groups. This results in the initial set of non-redundant possible predictive features for the ML classifiers, and their group-to-variant mappings, saved into 'df_groups_variant_info.py'.

Next, the pipeline prepare_ML_classifier_inputs.py defines train / evaluation / test splits for each sample set, as implemented in GWASfun.get_phenotype_and_phenotype_transitions_df_all_sample_sets. First, for each sample_set it first splits the whole dataset into five folds (due to --min_pheno_transitions 5), each corresponding to a different ~80 / 20 % split used for training and evaluation / testing, thus leading to one of the m1-m5 models. These folds, each corresponding to a 0-4 'test_split_ID', i) have a class imbalance equivalent to the one of the original data and ii) are generated to be phylogenetically balanced, including different clades in different folds (**Extended Data Fig. 6b**). For this, the pipeline uses GWASfun.get_phenotypes_df_with_balanced_split_IDs with n_splits = 5, min_support = 70 and ASR_method = "DOWNPASS", leading to the same folds for different min_support / ASR_method 'train parameters'. Then, within each of these folds ('test_split_ID' 0-4) and different 'training_type_split' values, our pipeline defines five independent training / evaluation 4x cross-validation folds (due to '--n_CVset_resamples 5', related to the identifier 'CVset_resample_I' 0-4), which may result from a splitting that is random (training_type_split = 'random') or phylogenetically-balanced with GWASfun.get_phenotypes_df_with_balanced_split_IDs (training_type_split = 'balanced'). This train / evaluation split is performed with GWASfun.get_phenotypes_df_CVset_with_resamples. Also, note that our pipeline saves the splits used for training / evaluation into 'phenotypes_df_all_CVset.py', and the ones for testing into 'phenotypes_df_all_test.py'.

Also, to enable prediction of phenotypes from prediction of phenotypic transitions (see parameter 'type_phenotype' above), our pipeline defines various pairwise isolate comparisons within each of these splits, and whether they constitute phenotypic transitions or not, as implemented within GWASfun.get_phenotype_and_phenotype_transitions_df_all_sample_sets. For this, it first runs GWASfun.get_df_pairwise_SNP_distances to calculate the genetic distance between all pairs of isolates (different SNPs / kb), enabling further filtering based on such distances. Then, it runs GWASfun.get_phenotype_transitions_df_ML_predictors on the set of isolates of a given training / evaluation / testing split, keeping certain comparisons depending on the value of 'transition_definition'. Specifically, for transition_definition = 'two_close_balanced' this function keeps, for each isolate, comparisons to the two closest isolates (with least SNPs/kb) that have the same or a different phenotype. Conversely, for transition_definition = 'each_pheno_clade' this function identifies monophyletic clades for each phenotype value and keeps, for each isolate, a comparison with one representative (with least SNPs / kb) isolate of each monophyletic clade. At the end of these procedure, the files 'phenotype_transitions_df_all_CVset.py' and ‘phenotype_transitions_df_all_test.py’ contain the comparisons and phenotype transition information for all the training / evaluation or test folds, respectively.

Furthermore, to generate a table with the presence / absence pattern of each non-redundant predictive feature (group of variants generated in GWAS, defined above) in each isolate prepare_ML_classifier_inputs.py runs GWASfun.get_ML_classifiers_features_df_all. Conversely, to generate a similar features table for each isolate comparison, reflecting whether there are variants of a given feature appearing in one of the isolates of the comparison vs the other, our pipeline runs GWASfun.get_ML_classifiers_features_transitions_df_all. All of these feature tables, called ‘features_df_all.py’ and ‘features_df_transitions_all.py’, were useful for further ML modelling.

Finally, to generate a list of features (variant groups) with genotype transitions associated with phenotype transitions within each training / evaluation set, which would further serve as an initial set for feature selection when gene_set is not ‘all’, we used GWASfun.generate_feature_files_each_CVset. This function runs, for the isolates of each sample_set and training / evaluation fold, the pipeline run_GWAS.py (described above) with arguments ‘--resampled_phenotypes_pastml_out skip --min_support <min_support> --ASR_method <ASR_method> --classification_method_ASR all_ignoreNaN --all_pval_methods skip --interesting_gwas_methods synchronous --min_n_geno_transitions 2 --skip_QQ_plots --skip_ASR_integration --filter_groups_raw_correlation’ (env:ancestral_GWAS_env). This generates, for features (groups) where the presence / absence pattern of variants across isolates is associated with the phenotypic presence / absence pattern (Fisher $p < 0.1$, due to --filter_groups_raw_correlation), calculations of $n_{G\&P}$ for gwas_method = ‘synchronous’ (described above). At the end, our pipeline generates a ‘target_features_CVsets.tab’ including the ‘synchronous’ $n_{G\&P}$ and $p_{FISHER}$ for all features with $n_{G\&P}$ >= 2 (due to ‘--min_geno_pheno_transitions 2’) in each each sample_set and training / evaluation fold.

On another line, the arguments ‘--pathway_annotations_dir --interproscan_output --CGD_chromosomal_features --reactome_organism_priority’ are set to process the features related to domain, gene and pathway annotations, as done for the GWAS (see above). Also, the ‘--n_phenotype_resamples 10’ was set to generate 10 training / evaluation folds with randomly resampled phenotypes for further ML maxT-like p value calculations (see above), but we finally decided not to use these calculations due to their computational cost and a lack of evidence of its usability for ML. Furthermore, for some combinations of phenotype, aligner, min_support and ASR_method we could not find any sample_set yielding >= 5 phenotypic transitions, and we discarded these combinations. This explains why the numbers of ‘train parameters’ differ across phenotypes.

##### Running training and evaluation

To run the training and evaluation process we ran the ancestral_GWAS toolkit script train_ML_classifier_CVset.py (env:ancestral_GWAS_env), for each combination of phenotype, aligner, min_support, ASR_method, transition_definition, sample_set, type_phenotype (‘phenotype_transition’ or ‘phenotype’), test_split_ID (0-4), gene_set (defined above) and type_filtering (‘p < 0.05’ or ‘all’ when gene_set = ‘all’, see **Methods**). Specifically, we set the arguments ‘--sample_set <sample_set> --test_split_ID <test_split_ID> --type_phenotype <type_phenotype> --features_max_pval_GWAS <p value threshold> --input_files <prepare_ML_classifier_inputs.py output directory> --pathway_annotations_dir <prepare_pathway_annotations.py output directory> --interproscan_output interproscan_annotation.out --CGD_chromosomal_features <CGD tabular file> --reactome_organism_priority Saccharomyces_cerevisiae,Schizosaccharomyces_pombe --expected_genes <expected genes> --SNP_kb_thresholds 0.1,0.25,1.0 --threshold_definition_transitions 0.25,0.5,0.75 --default_probability_pheno <default phenotype probability> --min_AUC_feature_sel 0.6 --min_n_features 2 --target_genes_training <set of genes of each gene_set>’.

There are various arguments that are related to the various train and test parameters. First, the p value threshold passed to --features_max_pval_GWAS was 0.05 (for type_filtering = ‘p < 0.05’), 1.0 (for type_filtering = ‘all’) or ‘-1’ (for gene_set different from ‘all’). Second, to --expected_genes we pass the comma-separated ordered list of expected genes for azole and echinocandin phenotypes (see ‘Defining target gene sets’ above), or ‘skip’ for the others, which are related to the ‘prioritize_expected_genes’ parameter (described below and in **Methods**). Third, the arguments ‘--pathway_annotations_dir --interproscan_output --CGD_chromosomal_features --reactome_organism_priority’ are set to process the features related to domain, gene and pathway annotations, as done for the GWAS (see above). Fourth, to --default_probability_pheno we pass, for each phenotype and sample set, the fraction of isolates that have phenotype = 1 divided by the total number of isolates, which is used for setting a default phenotype probability. Fifth, to --target_genes_training we pass either ‘all’ (for gene_set = ‘all’) or a file that contains a list of the target gene IDs of each gene_set used for training and evaluation (see ‘Defining target gene sets’). Sixth, --input_files contains the output directory of prepare_ML_classifier_inputs.py (see above) for the corresponding phenotype, aligner, min_support, ASR_method and transition_definition, and train_ML_classifier_CVset.py uses the following files within it: df_groups_variant_info.py, phenotypes_df_all_CVset.py, features_df_all.py, phenotype_transitions_df_all_CVset.py, features_df_transitions_all.py and target_features_CVsets.tab.

With the set arguments, the pipeline train_ML_classifier_CVset.py initially loads the phenotypes or phenotype transitions to be predicted (referred below as 'target_phenotype') with their corresponding train / evaluation splits information, coming from either 'phenotypes_df_all_CVset.py' (type_phenotype = 'phenotype') or 'phenotype_transitions_df_all_CVset.py' (type_phenotype = 'phenotype_transition'). Also, it loads the ML feature tables, coming from either 'features_df_all.py' (type_phenotype = 'phenotype') or 'features_df_transitions_all.py' (type_phenotype = 'phenotype_transition'). Then, for settings where the initial feature filtering is based on GWAS (--features_max_pval_GWAS <0.05 or 1.0> and --target_genes_training all), it keeps those features from 'target_features_CVsets.tab' with a $p_{FISHER}$ < features_max_pval_GWAS in the corresponding sample_set and test_split_ID. Conversely, for settings where the initial features are those affecting only a subset of genes (--features_max_pval_GWAS -1 and --target_genes_training <set of genes of each gene_set>), our pipeline keeps features collapsed at the variant effect, domain or gene level (see type_collapsing above), involving genes within the target gene set.

Next, for the corresponding training / evaluation set (depending on --test_split_ID), train_ML_classifier_CVset.py calculates classifier performance and selected features for different combinations of CVset_resample_I, training_type_split, feature_selection_AUC_threshold, class_weight and model, using GWASfun.generate_models_one_CVset. Initially, this function generates a set of non-redundant, possibly predictive features for each combination of CVset_resample_I and training_type_split (see above), using GWASfun.generate_NR_features_one_combination_of_resamples. In brief, this procedure keeps features that have the following properties. First, they should have non-redundant presence / absence variant patterns across the isolates of the given training / evaluation set, prioritizing them through GWASfun.get_NR_group_series (see above). Second, they should be found in >=2 isolates or isolate comparisons (depending on type_phenotype). Third, they should be defined so that each representative variant (see 'mutation_to_representative_mutation.tab' above) is mapped to only one final feature, prioritizing those that have higher predictive power of the target_phenotype and are more specific, as implemented in GWASfun.get_df_feat_info_each_rep_mut_only_one_feature.

After this feature filtering, GWASfun.generate_NR_features_one_combination_of_resamples also runs GWASfun.get_df_features_with_interactions_NR to add new features reflecting two-way interactions between these non-redundant features. For this, the latter function iteratively adds interaction features that i) do not have a presence / absence variant pattern redundant with the ones of previously added features, ii) are in >=2 isolates or isolate comparisons (depending on type_phenotype) and iii) are associated with the target_phenotype (Fisher p value < 0.05). This results in the initial set of features to be used for subsequent ML modelling and feature selection, in each combination of CVset_resample_I and training_type_split. Note that these interaction features were added so that the logistic regression models (see below) could consider interactions.

Next, GWASfun.generate_models_one_CVset runs GWASfun.get_series_info_one_model to run feature selection and obtain predictive performance metrics for different combinations of CVset_resample_I, training_type_split, feature_selection_AUC_threshold (0.05, 0.1 or 0.15), class_weight (balanced or none) and model (see **Methods**, e.g. logistic regression, AdaBoost or random forest). Specifically, we build nine different interpretable ML models from scikit-learn (sklearn) (AdaBoost, RF, GBoost, KNN, BernoulliNB, GaussianNB, LR, SVC and tree), and in some of them we try different configuration hyperparameters (GBoost, KNN, SVC, tree), as described next. First, 'AdaBoost' (sklearn.ensemble.AdaBoostClassifier) refers to an ada boost tree ensemble method, run with 50 estimators, a decision tree of depth=1 as base estimator, and SAMME.R as the optimization algorithm. Second, 'RF' (sklearn.ensemble.RandomForestClassifier) indicates a random forest tree ensemble method, run with 100 estimators and bootstrapping. Third, 'GBoost' (sklearn.ensemble.GradientBoostingClassifier) refers to a gradient boosting tree ensemble method, run with 100 estimators and a maximum depth of 3, 5, 7 or 9. Fourth, 'KNN' (sklearn.neighbors.KNeighborsClassifier) indicates a K-nearest neighbors algorithm, run with leaf size of 30, and varying n_neighbors (3, 5) and weights (distance, uniform). Fifth, 'BernoulliNB' (sklearn.naive_bayes.BernoulliNB) refers to a Bernoulli naive bayes classifier run with default parameters. Sixth, 'GaussianNB' (sklearn.naive_bayes.GaussianNB) indicates a Gaussian naive bayes classifier run with default parameters. Seventh, 'LR' (sklearn.linear_model.LogisticRegression) refers to a logistic regression run with 'liblinear' solver and 'l2' penalty. Eight, 'SVC' (sklearn.svm) is a support vector classifier, run with either a linear or a rbf kernel, and for the rbf kernel, we also modified gamma parameter, taking values 'auto' and 'scale'. Ninth, 'tree' (sklearn.tree.DecisionTreeClassifier) is a decision tree classifier, with no pre-set number of maximum features, using 'gini' as the criterion for measuring the quality of splits, and changing i) the minimum samples per split (2 and 3), ii) the minimum samples per leaf (1 and 2) and iii) the max depth (3, 5, 7, 9 and none).

The function GWASfun.get_series_info_one_model first builds the unfitted sklearn model object with GWASfun.get_predictor_model depending on the model type, configuration hyperparameters and the set class weighing (argument 'class_weight' of the sklearn object). Then, to perform sequential feature selection using this model it runs sklearn.feature_selection.SequentialFeatureSelector with n_features_to_select = "auto", direction = "forward" and tol = <feature_selection_AUC_threshold>, using GWASfun.get_integrated_ROC_AUC_across_CVs_CVset_feat_selection as the scoring function. With these settings, in each iteration the feature selection pipeline adds a new feature that maximizes ROC AUC, calculated by running sklearn.metrics.roc_auc_score on the aggregated target_phenotype probabilities of different evaluation folds, until the increase in ROC AUC is below feature_selection_AUC_threshold. This results in the minimal set of predictors yielding maximum predictive performance. Finally, this function runs GWASfun.get_integrated_ROC_AUC_across_CVs to get the final ROC AUC measurement (called 'CVset_ROC_AUC') of these minimal sets of features on the corresponding training / evaluation set.

There are some steps of this general pipeline that are adjusted depending on the specific dataset and modelling procedure. First, we skip KNN modelling settings where the n_neighbors is higher than the minimum number of isolates / isolate comparisons across all training sets. Second, we only consider two-way interaction features for LR models, as the others already model interactions natively. Third, for some training / evaluation folds with insufficient variability in training data (i.e. only one isolate / isolate comparison or all features having the same presence / absence pattern ) we assigned a default value to the corresponding validation target_phenotype probabilities. This was either the value passed to --default_probability_pheno (see above) for type_phenotype = 'phenotype', or a simulated default transition probability based on --default_probability_pheno (see GWASfun.get_pheno_trans_prob_from_pheno_prob) for type_phenotype = 'phenotype_transition'.

Fourth, the class_weight was only applied to certain datasets. To check whether to use it, our pipeline calculates the mean fraction of isolates with target_phenotype = 1 across different training / evaluation sets of the corresponding sample_set (see GWASfun.get_mean_fraction_1_real_phenos). Then, only if this number was above 0.8 or below 0.2 it runs the modelling with class weighting (class_weight = 'balanced') or without it (class_weight = 'none'). Conversely, for other datasets where this was not the case our pipeline only runs the modelling with class_weight = 'none'. All these processes result in a set of selected predictors and 'CVset_ROC_AUC' for each CVset_resample_I, training_type_split, feature_selection_AUC_threshold, class_weight and model. After modelling, to add gene and variant information to these different models, train_ML_classifier_CVset.py runs GWASfun.get_df_models_CVset_with_affected_genes_info. This function records, for each set of predictors selected by each model, their related i) set of genes, ii) set of genes within the 'expected genes' (defined above) for azole and echinocandin phenotypes, iii) linkage_groupID for further redundancy reduction, and iv) underlying variants. All of these were obtained by parsing the ML input files as provided by --input_files.

Next, train_ML_classifier_CVset.py runs GWASfun.generate_models_one_CVset_integrated to get the best model and related features, with optimal CVset feature consistency and ROC AUC across CVset_resample_I resamples, for different combinations of training_type_split, feature_selection_AUC_threshold, class_weight and model. Also, this function considers some additional parameters (consistency_measure, prioritize_expected_genes, AUC_diff_threshold, transition_SNPkb_threshold, transition_used_prediction and transition_ML_threshold, see **Methods**) for best model selection. Thus, it results in a final dataset with the best model and features for each combination of training_type_split, feature_selection_AUC_threshold, class_weight, model, consistency_measure, prioritize_expected_genes, AUC_diff_threshold, transition_SNPkb_threshold, transition_used_prediction and transition_ML_threshold.

This function calculates, for each feature set yielded by models using different CVset_resample_I values for each of these combinations, the CVset feature consistency and the CVset_min_AUC. CVset feature consistency is calculated as the average jaccard index between the set of variants, strains or genes (depending on consistency_measure) related to the CVset_resample_I's features vs the features yielded by other CVset_resample_I values. Conversely, CVset_min_AUC is calculated as the minimum ROC AUC obtained with sklearn.metrics.roc_auc_score, when using such features across CVset_resample_I folds. In this case, the AUC is calculated considering phenotype probabilities that come from either i) direct phenotype predictions (type_phenotype = 'phenotype'), or ii) predictions of phenotype transitions (type_phenotype = 'phenotype_transition') transformed to phenotype probabilities depending to the parameters transition_SNPkb_threshold (0.1, 0.25, 1.0), transition_used_prediction ('all', 'only_transitions', 'only_no_transitions') and transition_ML_threshold (0.25, 0.5, 0.75) (see **Methods**, **Extended Data Fig. 15**, GWASfun.get_df_predicted_phenotypes_from_transitions). Finally,

GWASfun.generate_models_one_CVset_integrated keeps the best model as the one that has i) a CVset_min_AUC within 0.05, 0.1 or 0.15 (AUC_diff_threshold) of the maximum CVset_min_AUC across CVset_resample_I values , ii) features including some of the expected genes if prioritize_expected_genes is set and any such models are found, and iii) the first model when hierarchically sorting them by CVset feature consistency (if consistency_measure is different from 'none') (prioritizing higher values), CVset_min_AUC (prioritizing higher values), number of features (prioritizing lower values) and number of variants related to the features (prioritizing lower values). Also, note that, due to '--min_AUC_feature_sel 0.6 --min_n_features 2', this function omits parameter combinations where none of the CVset_resample_I models had at least 2 features and a CVset_ROC_AUC >= 0.6.

In summary, we used train_ML_classifier_CVset.py to generate all the performance estimates and features selected when using different models and parameters.

##### Integration and testing of ML models

To integrate all of the training and evaluation measurements and get performance estimates on the testing data we ran integrate_ML_classifier_results.py (env:ancestral_GWAS_env) on each combination of phenotype, aligner and gene_set. Specifically, we ran it with arguments '--pathway_annotations_dir <pathway annotations directory> --paths_training <paths training file>', defined next. First, --pathway_annotations_dir is set to process the features related to domain, gene and pathway annotations, as done for other GWAS and ML analyses (see above). Second, to --paths_training we passed a file including the paths to the different training and evaluation results, mentioned above.

This script initially runs GWASfun.get_df_best_models_each_CVset to select the optimal training parameters for each combination of testing parameters (aligner, gene_set, training_type_split, sample_set, AUC_diff_threshold and consistency_measure) and test_split_ID. For this, it keeps training parameters with a CVset_min_AUC within 0.05, 0.1 or 0.15 (depending on AUC_diff_threshold) of the maximum CVset_min_AUC across training parameters. Then, the optimal training parameters are first ones in a list of hierarchically sorted ones according to the following criteria. First, parameters prioritizing expected genes are prioritized (parameter prioritize_expected_genes), in a way that gives more importance to genes that are first in the list of expected genes (i.e. *ERG11*, *MRR1*, *Scer_HAL9* (*TAC1*), *NDT80* for azoles; *FKS1, FKS2* for echinocandins). Second, parameters with the highest CVset feature consistency are prioritized.

Third, predictions of phenotypes are prioritized over predictions of transitions (parameter type_phenotype). Fourth, for parameter transition_definition, the 'each_pheno_clade' is prioritized over 'two_close_balanced'. Fifth, for parameter transition_SNPkb_threshold, higher values are prioritized over lower ones. Sixth, for parameter transition_used_prediction, 'all' is prioritized over 'only_no_transitions', and 'only_no_transitions' is prioritized over 'only_transitions'. Seventh, models with fewer features are prioritized. Eighth, models with highest CVset_min_AUC are prioritized. Ninth, lower p values for GWAS feature filtering are prioritized (parameter type_filtering). Tenth, parameters using higher min_support values are prioritized. Eleventh, for feature_selection_AUC_threshold, 0.1 is prioritized over 0.05, and 0.05 is prioritized over 0.15. Twelfth, for class_weight, no weighting is prioritized over balanced weighting. Thirteenth, ML models and settings (parameter model) are prioritized according to their order of definition in GWASfun.get_all_ML_models_list, which prioritizes different models in this order: RF, GBoost, AdaBoost, tree, LR, GaussianNB, BernoulliNB, SVC and KNN. Fourteenth, for ASR_method, DOWNPASS is prioritized over MPPA,DOWNPASS, and MPPA,DOWNPASS is prioritized over MPPA. Fifteenth, for transition_ML_threshold lower thresholds are prioritized.

Then, integrate_ML_classifier_results.py runs i) GWASfun.get_classifier_df_models_test to calculate the performance of each of these optimal sets of training parameters on the testing data, ii) GWASfun.get_df_models_test_with_fields_from_df_metrics to add diverse performance metrics, and iii) GWASfun.generate_integrated_performance_df to create the final datasets of ML results called 'best_models_test_performance_integrated_min_nmodels=<min_nmodels>.tab. We used the performance integrated dataset with min_nmodels = 4 for all further analyses, defined below, including one row for each combination of phenotype, testing parameters, and test_sample_set (i.e. whether the performance is calculated considering the all samples or only representative ones within the testing data, see **Extended Data Fig. 6a**). This table contains performance metrics integrated across all training / test splits (4 or 5, for models m1-m5) where the model was fit, i.e. with at least one CVset_resample_I having >=2 features and CVset_ROC_AUC >= 0.6 (see above). In the next paragraph we describe relevant fields in it, used for our ML analyses hereafter.

The column 'nmodels_fit' includes the number of training / test splits (up to 5) with some model fit. Rows where nmodels_fit is below 4 (as we used min_nmodels = 4) have their performance metrics to -1, flagging them to be skipped. In addition, the column 'ROC_AUC_test_integrated_onlyCorrectModels' includes the ROC AUC related to the 'test' testing strategy (**Fig. 5d**), calculated with sklearn.metrics.roc_auc_score on the aggregated phenotype probabilities for the m1-m5 fit models (called 'AUC_test' below). Similarly, the column 'mean_min_ROC_AUC_CVset_resamples' includes the average CVset_min_AUC across fit models, described above, and reflects the training ROC AUC (called 'AUC_train' below). Also, the column 'mean_cross_splits_ROC_AUC' includes the average 'cross_splits_ROC_AUC' across fit models, related to the 'cross-val' testing strategy (**Fig. 5d**). The 'cross_splits_ROC_AUC' metric is calculated for each m1-m5 independently, training on each of the training splits, getting phenotype probabilities on the testing splits, and aggregating them to calculate ROC AUC with sklearn.metrics.roc_auc_score. Conversely, the column 'mean_cross_splits_consistency_jaccard_genes' is the average feature consistency related to the genes linked to each predictor (called 'final_consistency' below), calculated with the jaccard index average as for CVset feature consistency (defined above). This is the feature consistency per testing parameters referred to above (see **Fig. 5c**). Finally, the columns 'min_n_train_parameters' and 'max_n_train_parameters' indicate the range of training parameters explored across fit models.

The script integrate_ML_classifier_results.py also outputs two other relevant files that we used. First 'best_models_test_performance.tab' includes the performance metrics for each combination of phenotype, testing parameters, test_sample_set and test_split_ID. Relevant fields are 'ROC_AUC_test_integrated_onlyCorrectModels' (i.e. AUC_test defined above) and 'cross_splits_ROC_AUC' (i.e. the cross-val ROC AUC for each m1-m5 model). Second, 'best_models_test_threshold_metrics.tab' includes the precision, recall, fpr and specificity across all possible phenotype probability thresholds for each combination of phenotype, testing parameters, test_sample_set, test_split_ID and eval_data (evaluation data), calculated as implemented in GWASfun.get_r_df_metrics_with_added_vals. Most importantly, the true phenotypes / predicted probabilities evaluation data may be the one used to calculate either i) 'ROC_AUC_test_integrated_onlyCorrectModels' (eval_data = 'test_integrated_onlyCorrectModels') for the 'test' strategy metrics, or ii) 'cross_splits_ROC_AUC' (eval_data = 'cross_splits') for the 'cross-val' strategy metrics.

To focus on relevant testing parameters, for each phenotype we defined as 'correct' parameters those with test_sample_set == 'representative_samples', AUC_test >= 0.7, AUC_train >= 0.7, final_consistency >= 0.4, and less than 0.2 difference between AUC_test and AUC_train (to avoid under or underfitting). Also, we defined as 'top' parameters those correct parameters that had an AUC_test within 0.05 of the maximum AUC_test across correct parameters, and had the top 10 final_consistency values out of them. Finally, we defined as 'best' parameters those with top final_consistency, or with highest AUC_test and AUC_train if there was >1 top parameter with the same highest final_consistency. Also, for the 'cross-val' testing strategy and visualization of single-model (m1-m5) results (e.g. **Fig. 5e,f**, **Extended Data Fig. 13**) we only considered those with a cross_splits_ROC_AUC within 0.05 of the maximum cross_splits_ROC_AUC across test_split_ID values.

To infer an optimal probability threshold for calculating precision, recall, fpr and specificity, we took one with highest recall while being within the range 0.05-0.95 and having fpr <= 0.3. Then, we obtained the corresponding metrics from 'best_models_test_threshold_metrics.tab', using eval_data = 'test_integrated_onlyCorrectModels' for the 'test' strategy and eval_data = 'cross_splits' for the 'cross-val' strategy.

#### Visualization of variants and genes

To visualize the variant presence / absence pattern and their associated genes along the strain tree (**Fig. 3, 6b,c, Extended Data Fig. 7,9,10,11,14,15,16,17,18**) we used script plot_tree_variants_multiple_phenos.py from the ancestral GWAS toolkit (env:ancestral_GWAS_env).

First, to plot information about genes previously linked to resistance towards azoles (**Fig. 3**), echinocandins, amphotericin B (**Extended Data Fig. 7**) or 5-flucytosine, we ran this script with various arguments defined next, as implemented in CPfun.plot_trees_expected_genes_variants_grouped_by_type_drug. To --variants_file and --variant_annotations_file we passed the variants' files from the bwa mem GWAS inputs folder (defined above) for the phenotype 'source blood' and sample_set=all_samples, including variants for all strains. To --phenotypes_file we passed the binary resistance phenotypes of each types of drugs, while to --phenotype_info_file we passed a file mapping each phenotype name to a shorter version for the visualization (e.g. Fl for fluconazole resistance). To --treefile we passed a tree pruned to include representative samples for all plotted phenotypes, defined as for the GWAS, and excluding 14 redundant samples for azoles' plot. To --group_info we passed a table with the target group IDs, referring to the non synonymous variants of each target gene (defined in **Main text**), and to --target_geneIDs we passed the set of genes to analyze. We also set '--min_support 70 --ASR_method DOWNPASS --classification_method_ASR all_ignoreNaN ' to parametrize how the transitions and nodes are visualized. Furthermore, we set '--gff <gff> --ref_genome <reference genome> --pathway_annotations_dir <prepare_pathway_annotations.py output directory>

--CGD_chromosomal_features <CGD tabular file> --interproscan_output interproscan_annotation.out --custom_geneID_maps Scer_HAL9:TAC1,Scer_ERG5:ERG5,Scer_FCY1:FCY1,Scer_FCY2:FCY2,Scer_FUR1:FUR1,Scer_CHS7:CHS7 --gDNA_code 12 --mtDNA_code 4' to configure how functional variant annotations are processed. To --sampleID_mapping we passed a table with information about each sample, including the ID, color and clade. Finally, we set --only_vars_correlated_phenotype to only visualize variants that appear together with any of the phenotypes, or in nodes that have some of the phenotypes. Note that we could not find any variants correlated to resistance in any of the expected genes for 5-flucytosine, so we skipped it.

Second, to visualize single variant GWAS hits for various phenotypes (**Extended Data Fig. 9,10,11**) we ran plot_tree_variants_multiple_phenos.py as implemented in CPfun.plot_GWAS_hits_in_trees. We used similar arguments as for the plots of expected genes, with a few differences. For instance, to --variants_file and --variant_annotations_file we passed variants generated for the hisat2 GWAS inputs for each phenotype, using those of 'balanced_samples' for high-confidence hits, and those of 'all_samples' for low-confidence hits, matching the filtering parameters that yielded such hits (see **Methods**). Similarly, to --treefile we passed the hisat2 tree of balanced or all samples, for high or low-confidence hits, respectively, generated for the GWAS inputs. Also, to --group_info we passed only single-variant hits. Furthermore, to --ASR_method we passed 'MPPA,DOWNPASS', as it was selected in the GWAS parameter optimization (see **Supplementary Results**). Also, instead of using --only_vars_correlated_phenotype, we set --variant_correlation_only_transitions to only visualize variants whose transition is associated with a phenotype transition.

Third, to visualize all features associated with fluconazole resistance by the top ML models (**Fig. 6b,c**), we ran the script as implemented in CPfun.visualize_top_ML_models_in_tree_all_genes_together, using equivalent arguments as for the expected genes but considering features related to genes found in >=3 top models. Fourth, to visualize the features of each of the top models (**Extended Data Fig. 14, 15, 16, 17, 18**) we ran plot_tree_variants_multiple_phenos.py as implemented in CPfun.visualize_top_ML_models_in_trees. This uses the equivalent arguments to those of expected genes, but without --only_vars_correlated_phenotype, and passing information about the phenotype transitions to --sampleID_mapping enabling the transition plots from **Extended Data Fig. 15**).

#### SUPPLEMENTARY TABLES

**Supplementary Table 1.** Table with all 409 strains analyzed in this study. It contains information on whether the strains are from the 365 initial or 44 extra (source_data), whether it has the "smiley" coverage pattern (has_smiley, see **Methods**), clinical information (country_collection, hospital, patient_age, patientID, year_collection), inferred genotypes and clades (microsatellite_genotype, clade_Bergin2022, clade_basedOn_Bergin2022), measured phenotypes (e.g. MICs), and binarized phenotypes (columns starting with 'binaryPheno').

**Supplementary Table 2.** Final GWAS parameters explored after hierarchical filtering. See **Supplementary Results** for more details on how these were obtained.

**Supplementary Table 3.** GWAS parameters that yield all expected genes (*MRR1/NDT80/ERG11* for fluconazole and voriconazole, *ERG11* for posaconazole and *FKS2* for caspofungin, see **Extended Data Fig. 8a**). Note that we could not find any such parameters if we had to find *TAC1* for posaconazole, so it was not considered. See **Supplementary Results** for more details on how these were obtained.

**Supplementary Table 4.** Non-redundant, filtered, high confidence GWAS hits. Each row corresponds to one combination of GWAS group (column unique_groupID), variant (column variantID_across_samples) and gene (columns Gene and final_name), which are associated with the phenotype of interest. The column 'group description' includes the short description of the GWAS group tested. The columns type_vars, type_mutations and type_collapsing indicate the variant grouping strategy (see **Fig. 4**). The column n_transitions_var indicates the number of phenotype transitions that are associated to a transition in each variant, and nodes_transitions indicates the comma-separated node IDs where these transitions happened. The columns nodes_GenoAndPheno, nodes_noGenoAndPheno, nodes_GenoAndNoPheno, nodes_noGenoAndNoPheno indicate, for a given group the numbers of nodes with and without phenotype and/or genotype transitions. The columns , OR, chi_square indicate the groups' association statistics. The column nodes_withPheno indicates the number of phenotype transitions for a given phenotype. The columns starting with 'pval_' indicate various association statistics (see **Supplementary Methods**). The columns, start, end and strand indicate the location of the gene, while the column description indicates its functional description. The column 'variant effect' indicates the variant effect on the gene. Finally, the columns starting with 'GO_', 'Reactome_' and 'MetaCyc_' indicate functional annotations of the genes.

**Supplementary Table 5.** Non-redundant, filtered, low confidence GWAS hits. The columns are as in **Supplementary Table 4**.

**Supplementary Table 6.** 79 gene-phenotype mappings including the most important genes for each phenotype. See **Methods** for more information about how we generated these, and to interpret the columns. The colors reflect genes that are found in multiple phenotypes.

35. Lu, H. *et al.* A Small Molecule Inhibitor of Erg251 Makes Fluconazole Fungicidal by Inhibiting the Synthesis of the 14α-Methylsterols. *mBio* **14**, e02639-22.

36. Choudhary, S., Mundodi, V., Smith, A. D. & Kadosh, D. Genome-wide translational response of Candida albicans to fluconazole treatment. *Microbiol Spectr* **11**, e02572-23.

51. Bliss, J. M. *et al.* Transcription Profiles Associated with Inducible Adhesion in Candida parapsilosis. *mSphere* **6**, e01071-20.

52. Neale, M. N. *et al.* Role of the Inducible Adhesin CpAls7 in Binding of Candida parapsilosis to the Extracellular Matrix under Fluid Shear. *Infect Immun* **86**, e00892-17 (2018).

53. Bertini, A. *et al.* Targeted gene disruption in Candida parapsilosis demonstrates a role for CPAR2_404800 in adhesion to a biotic surface and in a murine model of ascending urinary tract infection. *Virulence* **7**, 85–97 (2015).

54. Tóth, R. *et al.* Investigation of Candida parapsilosis virulence regulatory factors during host-pathogen interaction. *Sci Rep* **8**, 1346 (2018).

55. Dolichol-Linked Oligosaccharide - an overview | ScienceDirect Topics. https://www.sciencedirect.com/topics/chemistry/dolichol-linked-oligosaccharide.
